# ASCARIS: Positional Feature Annotation and Protein Structure-Based Representation of Single Amino Acid Variations

**DOI:** 10.1101/2022.11.03.514934

**Authors:** Fatma Cankara, Tunca Doğan

## Abstract

**Motivation:** Genomic variations may cause deleterious effects on protein functionality and perturb biological processes. Elucidating the effects of variations is critical for developing novel treatment strategies for diseases of genetic origin. Computational approaches have been aiding the work in this field by modeling and analyzing the mutational landscape. However, new approaches are required, especially for accurate and comprehensive representation and data-centric analysis of sequence variations.

**Results:** In this study, we propose ASCARIS (Annotation and StruCture-bAsed RepresentatIon of Single amino acid variations - SAVs), a method for the featurization (i.e., quantitative representation) of SAVs, which could be used for a variety of purposes, such as predicting their functional effects or building multi-omics-based integrative models. In ASCARIS representations, we incorporated the correspondence between the location of the SAV on the sequence and 30 different types of positional feature annotations (e.g., active/lipidation/glycosylation sites; calcium/metal/DNA binding, inter/transmembrane regions, etc.) from UniProt, along with structural features such as protein domains, the location of variation (e.g., core/interface/surface), and the change in physico-chemical properties using models from PDB and AlphaFold-DB. We also mapped the mutated and annotated residues to the 3-D plane and calculated the spatial distances between them in order to account for the functional changes caused by variations in positions close to the functionally essential ones. Finally, we constructed a 74-dimensional feature set to represent each SAV in a dataset composed of ∼100,000 data points.

We statistically analyzed the relationship between each of these features and the consequences of variations, and found that each of them carries information in this regard. To investigate potential applications of ASCARIS, we trained variant effect predictor models that utilize our SAV representations as input. We carried out both an ablation study and a comparison against the state-of-the-art methods over well-known benchmark datasets. We observed that our method displays a competing performance against widely-used predictors. Also, our predictions were complementary to these methods which is probably due to fact that ASCARIS has a rather unique focus in modeling variations. ASCARIS can be used either alone or in combination with other approaches, to universally represent SAVs from a functional perspective.

**Availability and implementation:** The source code, datasets, results, and user instructions of ASCARIS are available at https://github.com/HUBioDataLab/ASCARIS.

## 1. Introduction

Nonsynonymous single nucleotide variations have been associated with diseases (Hindorff et al., 2009; Manolio et al., 2008) due to their effects, such as perturbing biological processes and impairing molecular functions of proteins by changing their stability or interactions (Fariselli, et al., 2015; Datta et al., 2015; Farh et al., 2015; Halushka et al., 1999; Hindorff et al., 2009; Khurana et al., 2016; Presnyak et al., 2015; Sauna and Kimchi-Sarfaty 2011; Supek et al., 2014; Zwart et al., 2018; Dincer et al., 2019). Interpreting the effect of variations is important for understanding diseases of genetic origin, proposing effective treatment strategies, and developing novel biotechnological products (Dincer et al., 2019). High-throughput technologies have been producing vast amounts of variation data that awaits interpretation. However, experimental investigation of these variations remains challenging due to extensive resource-centric requirements such as labor and time. For this reason, accurate computational methods are necessary for prioritizing variants to direct experimental analysis and expedite validation. Computational approaches have been used in variation analysis, yet most of the research so far only focused only on predicting the effect of variation. On the other hand, recent developments in artificial intelligence-related technologies led to a surge of new algorithms and methods to be used in bioinformatics and computational biology (Unsal et al., 2022). In order for these algorithms/methods to be able to process biological entities and phenomena, these entities must be represented numerically. Therefore, there is a current need for new approaches to yield accurate, comprehensive and reusable numerical representations (i.e., feature vectors) of sequence variations.

There are numerous computational methods/tools for predicting the effects of variations at the level of genes or proteins (Calabrese et al., 2009; Adzhubei et al., 2010 & 2013; Bromberg et al., 2007 & 2008; Carter et al., 2009; Chennen et al., 2020; Choi et al., 2015; Clifford et al., 2004; Fariselli, et al., 2015; Kaminker et al., 2007; Ng et al., 2003; Pandurangan et al., 2020; Pires et al., 2014; Quan et al., 2016; Rentzsch et al., 2019; Savojardo et al., 2016; Schwarz et al., 2014; Tavtigian et al., 2008; Topham et al., 1997; Worth et al., 2011; Yang et al., 2013; Yue et al., 2006). Most of these methods/tools aim to answer the question of whether a given substitution (or indel) can be deleterious at the molecular level and/or at the whole organism scale, by utilizing the available biological information. These methods differ in terms of the employed features and implemented algorithmic techniques (please see Supplementary Information S1 for a review of variant effect prediction methods/tools). One of the current and prevailing issues in this regard, especially associated with machine learning-based approaches, is the interpretability of results (Unsal et al., 2022). It is crucial for a researcher, the tool’s user, to comprehend why the method predicted that specific outcome. Another critical point here is the input data and its featurization. The choice of source/input feature type(s) and the dataset are at least as important as the algorithmic approach used. In this regard, types of features that are unexplored in the framework of variant modeling are potential subjects of investigation, where they can be combined with traditional approaches, with the ultimate aim of constructing new models with performances that are sufficiently high to effectively aid clinical decision-making.

Residue/region specific annotations of proteins (e.g., nucleotide/DNA binding regions, active sites, modified residues, motifs, domains, etc.) provides crucial information about both their molecular functions and the biological mechanisms in which they are involved. This knowledge is collected, organized, and presented to the user in a standard format in protein-centric resources such as the UniProt database (The UniProt Consortium, 2019). Especially, the ontology-based and protein-centric versions of these annotations (e.g., Gene Ontology term associations of whole proteins) have been studied within the context of predicting functions (Kulmanov & Hoehndorf, 2020; Rifaioglu et al., 2018) and phenotypic implications (Doğan, 2018) of proteins; however, they are under-explored in the framework of variant modeling, with only a few examples (Calabrese et al., 2009; Capriotti et al. 2013). Essentially, the correspondence between a SAV of interest and a positional annotation on the sequence may reveal its effect at the molecular level.

In this study, we propose ASCARIS, a new function centric featurization approach to represent single amino acid variations (SAVs) based on their spatial correspondence with the residues/regions with functional importance, such as domains, active sites, binding sites, disulfide bridges, etc. Our hypothesis is simply that mutations that directly correspond to functionally important sites/regions in proteins (or mutations that are proximally located to these functional regions in the 3-D plane) are more susceptible to causing deleterious effects since these localized roles can easily be disrupted by the respective amino acid change.

For this, we incorporated 30 different types of protein sequence annotations from UniProtKB (the full list is provided in Table 1). Due to the fact that variations do not often directly correspond to annotated positions, we also took the spatial distance between the SAV and the annotated positions/regions into account by using the coordinates in the 3-D structure of the protein and incorporated this distance-based information into our variation feature vectors. Furthermore, we included amino acid specific physicochemical and structural changes caused by these SAVs such as polarity, volume, and accessible surface area, together with the location of mutations in the structure, with the aim of characterizing variations in a more context-dependent manner. To obtain 3-D features, we utilized two different tracks; (i) PDB + homology modeling, and (ii) AlphaFold tool’s structure predictions. The latter allowed the extension of our variant featurization method to proteins with completely unknown 3-D structures.

**Table 1.** Types of positional annotations from the UniProt database that are incorporated into the proposed SAV representations.

| Annotation Class | Annotation Type | Description |
| --- | --- | --- |
| Region | Coiled Coin | Positions of regions of coiled coin within the protein |
|  | Motif | Short sequence motif of biological interest |
|  | Region | Region of interest in the sequence |
|  | Repeat | Positions of repeated sequence motifs or repeated domains |
|  | Zinc Finger | Position(s) and type(s) of zinc fingers within the protein |
|  | Calcium Binding | Position(s) of calcium binding region(s) within the protein |
|  | DNA Binding | Position and type of a DNA-binding domain |
|  | Nucleotide Binding | Nucleotide phosphate binding region |
|  | Intramembrane | Extent of a region located in a membrane without crossing it |
|  | Transmembrane | Extent of a membrane-spanning region |
|  | Topological Domain | Location of non-membrane regions of membrane-spanning proteins |
| **Sites** | Active Site | Amino acid(s) directly involved in the activity of an enzyme |
|  | Binding Site | Binding site for any chemical group |
|  | Metal Binding | Binding site for a metal ion |
|  | Site | Any interesting single amino acid site on the sequence |
| **Amino Acid Modification** | Cross-link | Residues participating in covalent linkage(s) between proteins |
|  | Disulfide Bond | Cysteine residues participating in disulfide bonds |
|  | Glycosylation | Covalently attached glycan group(s) |
|  | Lipidation | Covalently attached lipid group(s) |
|  | Modified Residue | Modified residues excluding lipids, glycans and protein cross-links |
| **Variants** | Natural Variant | Description of a natural variant of the protein |
|  | Mutagenesis | Site which has been experimentally altered by mutagenesis |
| **Secondary Structure** | Beta Strand | Beta strand regions within the experimentally determined protein structure |
|  | Helix | Helical regions within the experimentally determined protein structure |
|  | Turn | Turns within the experimentally determined protein structure |
| **Molecule Processing** | Peptide | Extent of an active peptide in the mature protein |
|  | Pro-peptide | Part of a protein that is cleaved during maturation or activation |
|  | Signal | Sequence targeting proteins to the secretory pathway |
|  | Initiator methionine | Cleavage of the initiator methionine |
|  | Transit Peptide | Extent of a transit peptide for organelle targeting |

ASCARIS is not a variant effect predictor, in particular. Nonetheless, as an example potential application of the proposed method, we trained machine learning-based classification models (utilizing the 68-dimensional numerical features as input vectors, which are obtained from our SAV representations by removing the meta-data columns) to predict the effects of SAVs as deleterious or neutral. We trained and validated prediction models with more than 100,000 variation data points collected from the UniProtKB (The UniProt Consortium, 2019), ClinVar (Landrum et al., 2018), and PMD (Kawabata et al., 1999) databases, and compared the predictive performance with state-of-the-art variant effect predictors. The schematic representation of the study is given in Figure 1. One of the main advantages of our method is that it practically incorporates structure-related information for variant modelling using fundamental properties and residue/region-based functional features without costly molecular calculations and sequence alignments. Another advantage of our approach is that it generates completely interpretable feature vectors, where each dimension corresponds to a pre-defined structural or annotation-based property. Our alignment-free featurization approach can be used to represent SAVs as concise numerical vectors to be used in various types of computational approaches, e.g., combining them with traditional conservation-based features under ensemble methods to predict the effects of variations with elevated performance and/or coverage, or as part of multi-omics-based datasets in large-scale integration and modeling of biomedical data.

**Figure 1.**
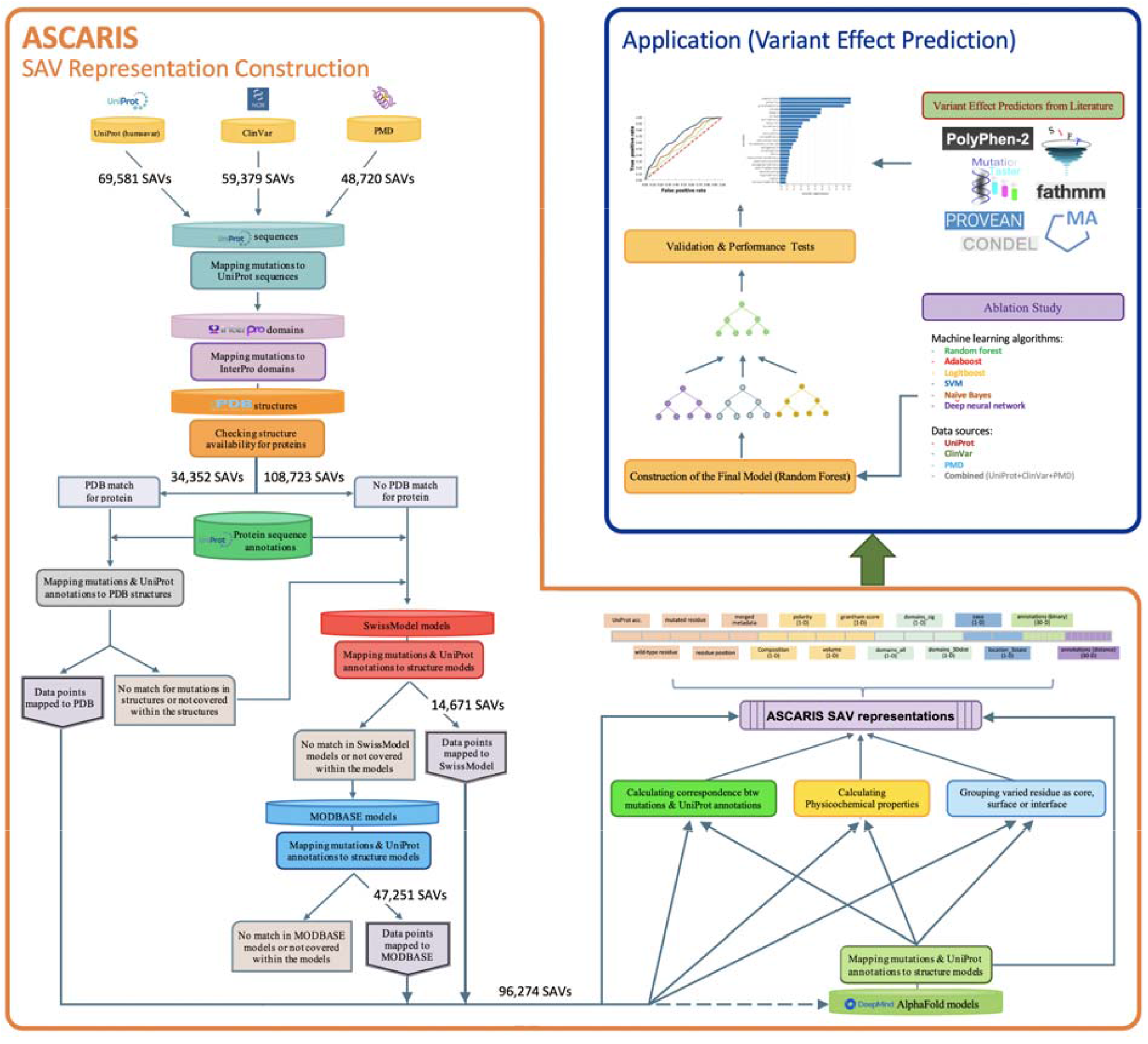
The overall workflow of the proposed variant featurization/representation method, ASCARIS, together with the application of it to the problem of variant effect prediction.

## 2. Methods

### 2.1. Data

The variation datasets were retrieved from three databases, namely UniProt, ClinVar, and Protein Mutant Database (PMD). To be able to use variation data points from different databases together, we grouped each variation into one of the two classes, namely “neutral” and “deleterious”. From UniProt’s (v2019_01) human variation (“humsavar”) dataset (current link: https://ftp.uniprot.org/pub/databases/uniprot/current_release/knowledgebase/complete/docs/humsavar), we obtained 40,02 polymorphisms and 29,553 disease associated SAVs for 12,519 human protein entries. Here, we labeled polymorphisms as neutral mutations, while variations associated with a disease condition are labeled as deleterious.

The second database, ClinVar, was used to retrieve clinically reported variants. Among the variation data points in the ClinVar database version 1.61 (downloaded from: https://ftp.ncbi.nlm.nih.gov/pub/clinvar/tab_delimited/), “pathogenic” and “likely pathogenic” variants were considered members of the deleterious class, whereas, “benign” and “likely benign” variants were considered neutral. The rest of the variation data points in ClinVar were discarded since their annotated effect was considered ambiguous. ClinVar variant data points were mapped to UniProt protein sequences using Ensembl transcript IDs and the bioDBnet database (Mudunuri et al., 2009). As a result of these filtering and mapping operations, 17,945 benign (i.e., neutral) and 41,434 pathological (i.e., deleterious) mutations were retrieved for 4,132 human proteins.

The last data source, Protein Mutant Database (PMD), includes manually curated mutations and their consequences regarding protein’s stability, interaction(s) or functional changes, in terms of the severity of the effect. In the PMD SAV dataset (downloaded on February, 2020 from http://pmd.ddbj.nig.ac.jp, which is not accessible as of 2022), [++] and [+] signs denote an increase in the activity/stability, whereas “[--]” and “[-]” denote a decrease in the activity/stability, “[0]” denote complete loss of function, and “[=]” denote no change. An increase or a decrease in the activity, no matter what the magnitude is, has a potential to impair the protein’s native state. For this reason, variations with increase and decrease in activity and/or stability, along with the ones with complete loss of function were recorded as deleterious, and no-effect cases were recorded for the neutral class. We collected 15,348 neutral and 33,372 deleterious variations for 3,401 distinct proteins (human and model organisms) from PMD.

To prepare the finalized dataset, we removed duplicate data points (i.e., data points that exist in multiple databases) (Figure S1a). Since different resources’ approaches to variant interpretation change, there were also a few conflicting cases, in terms of the annotated effect of these variations, i.e., labeled as neutral in one source, and deleterious in another (Figure S1b). Such conflicting data points were removed from our dataset. We also eliminated amino acid changes that resulted in a termination codon as they were highly biased to be deleterious. After these filtering operations, our dataset was composed of 144,075 SAVs (76,951 deleterious and 67,124 neutral) on 15,402 distinct proteins. Finally, we removed SAV data points for which 3-D structural information cannot be obtained (details of this elimination procedure is given below), making the finalized dataset of 96,274 SAVs, of which 52,512 are deleterious and 43,762 are neutral.

### 2.2. Incorporation of Structural Information

We obtained structure files for the proteins in our dataset from Protein Data Bank (PDB) (Berman et al., 2000). For the variation data points for which the corresponding protein’s structure has not been solved at all, or that the available structures do not span the region of the protein sequence where the variation is located, homology models from SwissModel (Waterhouse et al., 2018) and ModBase (Pieper et al., 2014) have been incorporated, in respective order. In the case of availability of multiple PDB structures or models that satisfy the above-mentioned conditions, the structure with the highest resolution or the model that possesses the highest quality score is retained. Variations without any corresponding structure or model were eliminated from the dataset. Figure 1 and Figure 2a show the workflow of the structure incorporation process, including the number of variation data points mapped at each step. At the end of this procedure, 96,274 mutations were associated with structural data, which constitutes our finalized variation dataset.

**Figure 2.**
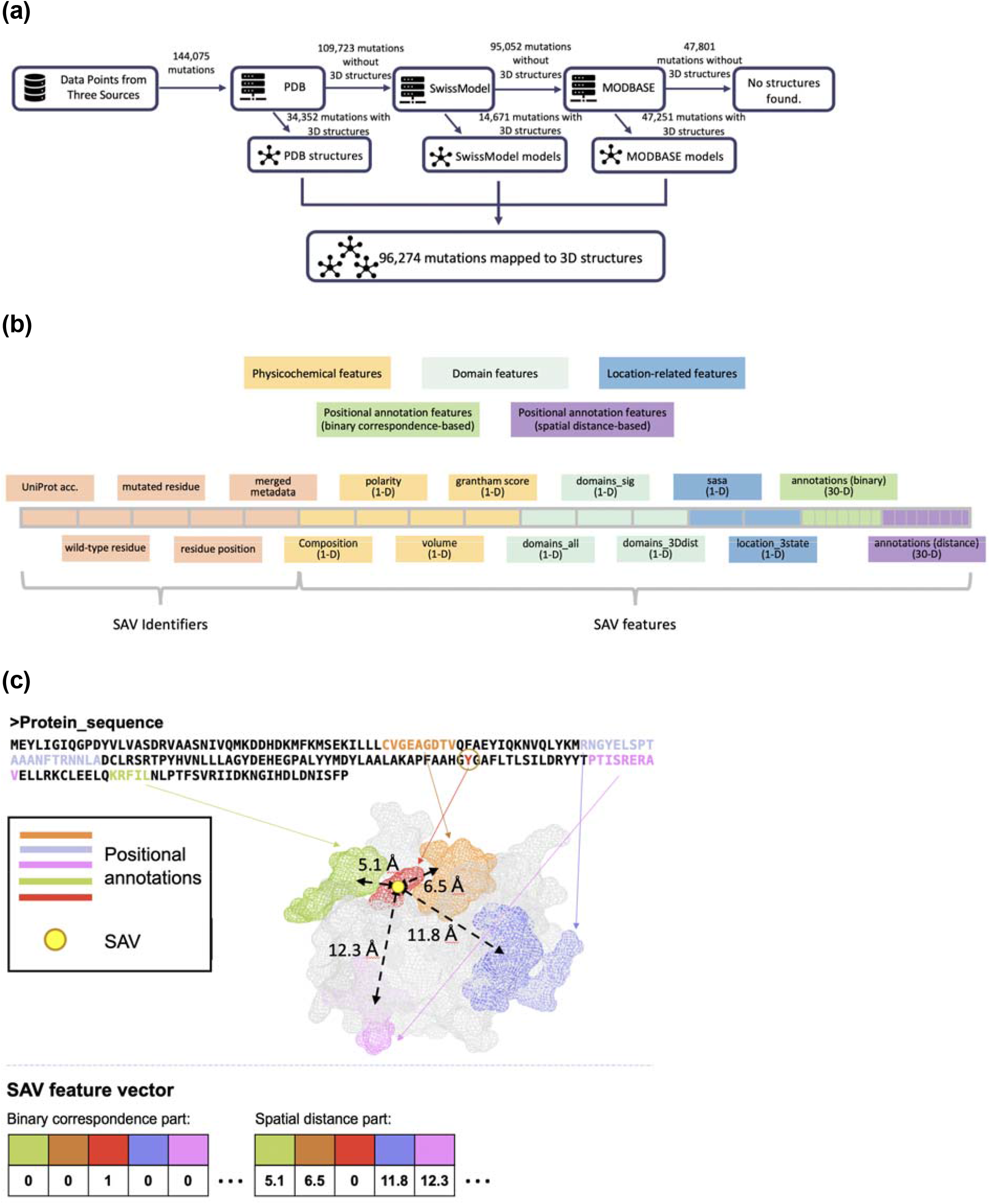
Dataset construction and featurization steps: **(a)** the sources of structural information and statistics about the data from each source, **(b)** the representation of different types of features on SAV representations, **(c)** mapping of positional sequence annotations and the SAV onto the 3-D protein structure, and part of the SAV feature vector that corresponds to these annotations.

As an alternative version of ASCARIS, we utilized the AlphaFold2 tool’s (Jumper et al., 2021) protein 3-D structure predictions instead of PDB and homology modeling. AlphaFold is a deep learning-based method developed by DeepMind that is capable of predicting protein structures from the primary amino acid sequences with extremely high accuracy (Jumper et al., 2021). We used the AlphaFold version of ASCARIS mainly to assess its performance and compare it against the original version. For this purpose, we downloaded structure models for the reference human proteome (release 2021_03) from AlphaFold-DB (at https://alphafold.ebi.ac.uk/).

### 2.3. Featurization

We featurized each SAV with a unique numerical representation to be later used for any predictive modeling task. Our representations contain information regarding multiple types of structural data including protein domains, physicochemical properties, structural location (i.e., core, interface or surface) and 30 different types of functional residue- or region-based annotations. This way, each SAV data point is represented by a 74-dimensional feature set including the meta data columns (e.g., accession of the protein, wild type and mutated residues, position, etc.). Table S1 and Figure 2b displays the names, descriptions, categories and number of dimensions that correspond to each type of feature on the representation. SAV features vectors that are used in ML-based variant effect prediction models are 68-dimensional and obtained by removing meta data columns from initial representations. We provided detailed information regarding each feature below.

#### 2.3.1. Protein Domains

Domain region annotations of proteins were retrieved from InterPro (Mitchell et al., 2019). In some cases, multiple domains from the same hierarchy (i.e., a group of domain entries that are roughly defining the same structure with different levels of specificity) are annotated to the same region of a protein. In such cases, the one at the highest level in the hierarchy (i.e., the most generic one) was considered and other domain annotations were discarded. In the case of multi-domain proteins, only the domain that spans the site of variation is retained. If none of the annotated domains spans the position of variation, the one closest to the variation, in terms of the number of amino acid positions in-between, was kept and the rest were discarded. This way, we retained domain information for 96,131 SAV data points out of 144,075 SAVs in the raw variation dataset. The remaining 47,944 SAV data points have not been associated with any InterPro domains. Out of the 96,131 SAVs, 31,893 of them have distantly located (i.e., out of region) domains, and 64,238 SAVs are found to be within the domain annotated regions. The unique number of retrieved domains was 2401.

Due to the high number of unique domains in our dataset, most of which were only encountered in one or a few SAV data points, we performed a statistical analysis using Fisher’s exact test to evaluate their significance considering the separation between neutral and deleterious SAVs. In other words, we observed the change in the frequency of observing deleterious mutations between the cases; (*1*) when the mutation is on the domain of interest, and (*2*) when the mutation is not on the domain of interest. Details and results of this analysis are provided in Supplementary Information S2. Based on the results of this analysis, we selected the most significant domains by takinf the reported p-values into account (Table S2). We incorporated the domain information into our SAV feature vectors using a categorical representation, where each domain is expressed by a unique numerical label assigned to it. As part of our ablation study, we compared the predictive performance of the variant effect prediction model that contained information on all domains found in our dataset, with the model that contained only the significant domains.

#### 2.3.2. Physicochemical Features

Physicochemical properties are evaluated at the individual amino acid level, considering their property value changes, as the difference between the wild-type amino acid to the mutated one. Differences in three different physicochemical features, i.e., polarity, volume, and composition, together with their consensus, Grantham Matrix Scores, are calculated for each SAV and incorporated into 4 dimensions of the corresponding feature vectors as real values. Amino acid-based volume and polarity values are taken from published data (Aboderin, 1971; Goldsack and Chalifoux, 1973; Grantham, 1974). The composition is calculated as the ratio between the atomic weight of non-carbon atoms and the total weight of carbon atoms in the side chain. These values indicate the magnitude of change in terms of physical constraints and provide a measure for the similarity/dissimilarity between the changed amino acid and the original one.

#### 2.3.3. Structural location of the variation

We incorporated the location of the mutated residue on the structure either as core, surface, or interface region since it may provide clues about the possible effect of the mutation (Capriotti et al., 2019; Engin et al., 2015; Guo et al., 2004; Nishi et al., 2013). We deduced this information from relative solvent accessible surface area (rASA) values. For this, we calculated solvent-accessible surface area (SASA) values for the residues of interest using FreeSASA (Mitternacht, 2016). In order to classify residues into one of the three groups (i.e., core, interface and surface), we applied a cut-off accessibility value of 5% to select between core and surface residues (Dincer, et al. 2019; Momen-Roknabadi et al., 2008). According to this, a residue with a rASA value less than 5% is considered as buried, while a residue with a rASA value greater than or equal to 5% is considered to be located at the surface. In order to differentiate between surface and interface, we directly employed protein-based interface residue information from InteractomeInsider (Meyer et al., 2018). Here, we selected validated interface residues along with high quality ÉCLAIR interface predictions. When a residue, that was previously labeled as surface, is listed as an interface residue in InteractomeInsider, it is removed from the surface group and placed into the interface group. When a previously core-labeled residue is found in the interface residue list, it is removed from the core residues group and labeled as conflicting. Structural location information is recorded using 2 dimensions of our variation feature vectors: a 1-D categorical variable using core/surface/interface grouping, and a 1-D real-valued variable containing the actual rASA values. 42,976 mutations in our dataset were found to be in the surface region, 12,900 of them in the core region and 5,810 are in the interface region. 724 mutations were labeled as conflicting. For the remaining data points, rASA values could not be calculated. Conflicting cases are later merged with those without any SASA values and treated as a 4th category.

#### 2.3.4. Mapping Positional Sequence Annotations

Sequence annotations were retrieved from the UniProt database version v2019_01 for each protein in our dataset. There are 34 different types of positional annotations in UniProt and we selected 30 of them. The types and descriptions of these positional annotations are given in Table 1. We included the positional annotation data via two different ways. First, we identified the annotated sites/regions which directly correspond to the SAV positions on the sequence. We incorporated this information into our feature vectors by reserving 30-dimensions, where each dimension belongs to a different type of positional annotation. We used a categorical variable to signify the correspondence between an annotation and the SAV of interest, i.e., ‘2’ if the SAV and annotation correspond to each other on the same sequence position, ‘1’ if that type of annotation exists on the protein of interest but the annotation and the SAV are not on the exact same position in the sequence, and ‘0’ if the annotation does not exist for that protein at all (Figure 2c).

Second, to account for the cases where there is no direct correspondence between the annotation and the SAV, we calculated the spatial distance in-between, using 3-D structural information. For this, we aligned full sequences of proteins in our dataset with the sequences of their corresponding PDB structures and identified the spatial location of both SAVs and the annotated residues, using the sequence-based position information. Then, we calculated the Euclidean distance between the C-alpha of both residues (i.e., SAV and the functionally annotated site/region) on the 3-D plane in the unit of Angstroms (Figure 2d). If the annotation is region-based instead of site based, the residue that is in the closest proximity with the SAV residue is taken into account. Some of the proteins have multiple sites/regions annotated with the same type of positional annotation. In such instances, the one closest to the site of variation is retained. The proximity information is incorporated into SAV representations via 30 additional dimensions, each of which corresponds to a different type of annotation, and the value inside is the real-valued spatial distance between the SAV and the corresponding annotated residue. If there is direct correspondence between a SAV and an annotation, distance value is recorded as 0. If the annotation type of interest does not exist for that protein or its position is out of the structurally solved regions of the protein, we impute the corresponding cells with the median spatial distance of the respective annotation type in the whole SAV dataset. These annotation type specific mean distance values are given in Table S3. Here, we did not use the value zero for the imputation in order to distinguish between a missing value and a true 0 distance (i.e., a case where the variation and the annotation correspond to the same position in the sequence).

### 2.4. The performance Analysis of AlphaFold Models

We created an alternative version of ASCARIS feature vectors using AlphaFold’s structure predictions as the source of 3D structural information. For this, we employed the same filtered SAV dataset, the preparation of which is described above. In AlphaFold-DB, the structure of a particular protein is separated into multiple models, if the sequence length is greater than 1400 residues. In our analysis, we evaluated all models given for a protein. If the variation is found in one model file and the annotation is found on the other model file, since these are two different spaces, direct distance calculation between these two residues is not possible. As a result, we only considered the comparisons if the variant and annotation of interest is mutually found in the same model. Also, some annotations and variations occur in multiple models. In such cases, we calculated the distance between the variation and the annotation within each model and considered the minimum of all distances for each annotation-variation pair.

### 2.5. Machine Learning-based Classification of Variants

In order to measure the biological relevance of our SAV representations, we trained classification models that use our representations as input and predict the effect of query SAVs as either neutral or deleterious. We evaluated the performance of our models, first, via 5-fold cross-validation, as reported under sections 3.2 and 3.3, and second, over independent hold-out test datasets for the comparison with state-of-the-art methods, which is reported under section 3.4.

In this study, we used the random forest (RF) algorithm for the binary classification of variation data points. The random forest algorithm is an extension of decision trees where multiple trees are built, an ensemble of which are used to make a decision (Breiman, 2001). A randomly selected subset of a given size is drawn with replacement from the original data and trees are built with each dataset separately (Dasgupta et al., 2011; Meyer et al., 2018). The RF algorithm randomly selects features to be used for splitting at each node, thus being less prone to overfitting.

In the analyses explained in section 3.2, we used the default hyper-parameters of random forest which can be listed as; the number of trees: 100, the maximum number of decision splits: n-1, and the number of predictors to select at random for each split: √p, where n and p represent the number of observations and the number of predictors, respectively. For the hyper-parameter optimization using grid-search, we tested the following values; the number of trees: 50, 150, 300, and 500; the maximum number of decision splits: 3, 81, 2187, 96273; and the number of predictors to select at random for each split: 2, 8, 24, 68. We also employed additional algorithms, such as Adaptive boosting (Adaboost) (Freund and Schapire, 1997), logistic regression (LogitBoost) (Friedman et al., 2000), support vector machine (SVM) (Cristianini and Shawe-Taylor, 2000), and the naive Bayes (NBayes) (Hastie et al., 2001), for algorithmic baseline model comparison.

In this study, Python (v3.7) was employed for data pre-processing and analysis, and feature vector construction jobs. MDS and t-SNE algorithms are implemented using Python’s (v3.7) sklearn (v0.21.1) library. MATLAB by MathWorks is employed for classification model development and performance evaluation. As far as we are aware, this is the only available and supported implementation to handle both categorical and real-valued variables in the same feature vector, without any limitation on the number of features.

### 2.6. Performance Metrics

8 different metrics are used for the evaluation of the performance of prediction models (i.e., accuracy, sensitivity/recall, specificity, negative predictive value - NPV, precision, F1-score, Matthew’s correlation coefficient - MCC, and the area under receiver operating characteristic curve - AUROC). Accuracy denotes the number of correct predictions to the total number of instances. Sensitivity/recall is a measure of the proportion of instances that are correctly predicted as positive with respect to all truly positive data points. Specificity denotes the proportion of negatives that are correctly predicted, to all cases of truly negative instances. Negative predictive value (NPV) is the proportion of negatives which are correctly predicted, to all cases of negative predictions. Precision is the proportion of correct positive predictions to the number of all positive predictions. In other words, it measures what proportion of positively predicted instances are actually correct. F1-score is the harmonic mean of precision and recall. Matthew’s correlation coefficient (MCC) is a balancing performance metric which takes true negatives into account along with the other constituents of the confusion matrix. MCC returns values between −1 and 1; (i.e., 1 representing the perfect correlation, 0: random prediction, and −1: a perfect negative correlation). Formulations of these performance metrics are given below:

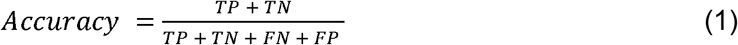

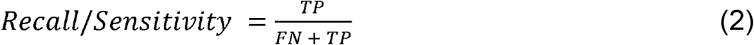

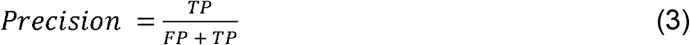

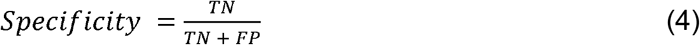

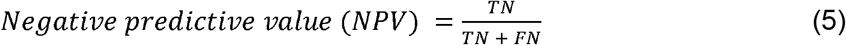

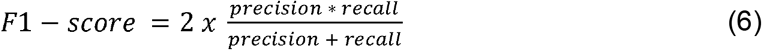

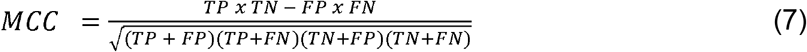

where, TP is the number of true positives, FP is the number of false positives, FN is the number of false negatives, and TN is the number of true negatives.

Some of the variant effect datasets used in this study are imbalanced with respect to the consequence of the mutation (i.e., deleterious/neutral). For this reason, MCC is taken as the principal performance metric since it provides reliable results in the case of class imbalance. In the benchmark comparison analyses, performance metrics employed in the respective studies are taken into account.

## 3. Results and discussion

### 3.1. Exploration of the Dataset and Features

In this section, we identified and discussed the relationship between the effects of variations and their structural and annotation-specific properties, to evaluate the biological relevance of incorporating these features. After that, we visualized our variation dataset on a 2-D plane via dimensionality reduction, based on our representation vectors, to observe the distribution of neutral and deleterious variations coming from different sources.

#### 3.1.1. Domain Annotation-Based Evaluation

We examined the relationship between a variant’s effect and its location with respect to annotated domain regions. We formed 3 groups for this purpose; “no domain” group signifies SAV data points where the corresponding proteins have no domain annotation in InterPro, whereas “within domain” and “out of domain” signify the variations that are located inside and outside the domain annotated regions on the sequence, respectively. As observed from Figure 3a, mutations are more likely to have a deleterious effect when they are located within domain annotated regions (59%), compared to the variations that remain out of domain regions (31%). We also statistically tested this observation using Fisher’s exact test and found the difference in deleteriousness to be statistically significant at a 99% confidence interval (p-value < 0.01). It is expected that a mutation that is within the region of a domain is more likely to cause a deleterious effect on the functionality of the protein, compared to a mutation on a non-domain (probably disordered) region, since domains are the main structural and functional building blocks of proteins (Doğan et al., 2016). It was also observed that the percentage of deleterious SAVs among no domain variations (61%) is similar to the one for the within domain variations (59%), which is plausible since it is highly probable that these so-called “no domain” proteins are understudied and have domains that are yet to be identified/documented in respective databases, and that many of these SAVs may actually reside in domain regions.

**Figure 3.**
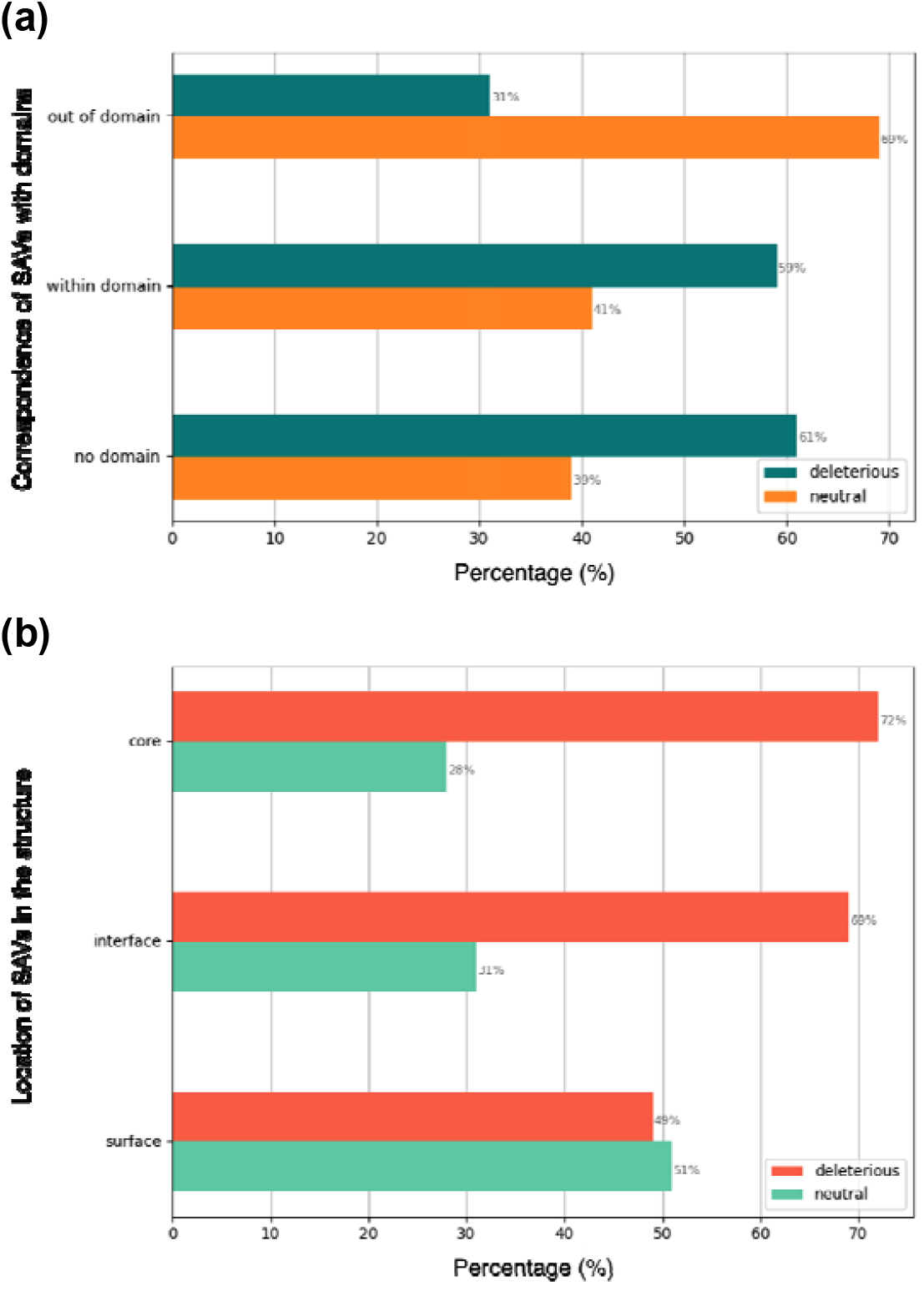
**(a)** Distribution of neutral and deleterious SAV data points according to their domain region correspondence. Mutations found within the domains tend to be more deleterious (59%) compared to the ones outside (31%), **(b)** distribution of neutral and deleterious SAV data points according to their location on the structure of the protein. Mutations found within the core and interface regions tend to be more deleterious (70% and 68%, respectively) compared to the ones on the surface (51%).

#### 3.1.2. Physicochemical Property-Based Evaluation

We analyzed changes in physicochemical descriptor values between the wild-type amino acid and the mutated one, in terms of polarity, volume, composition, and the Grantham score which represents the consensus of the former three. Since these physicochemical properties are given as relative values (i.e., changes occurred due to the variation, with respect to the wild-type amino acid in that position), their evaluation should be made accordingly. Here, SAVs with large physicochemical values indicate significant property changes and thus, expected to be deleterious, on the other hand, we expect to observe neutral variations with a higher ratio in the cases with insignificant property value changes. To test this, we applied statistical testing on each property, independently.

First, we drew histograms of the value distributions (Figure S4). Then, for each physicochemical property, we labeled each data point with conditions as “significant change” or “non-significant change” via thresholding using a cut-off value determined with respect to the whole value distribution. Thresholds were chosen to leave approximately the same number of data points per group (i.e., significant and non-significant change). Since the change in polarity, volume and composition can be negative, as well as positive, two thresholds were selected for each property type. Data points with polarity values higher than 1.6 or lower than −1.6 are considered for the “significant change” group, while data points with polarity values between −1.6 and 1.6 are considered for the ‘non-significant change’ group. For the volume property, the threshold values are set to −38 and 38. For the composition property, the thresholds are −0.52 and 0.52. Since the calculation of Grantham score comprises the summation of weighted and squared values of individual property differences, the minimum value is 0, as a result, only a positive threshold is set, which is 81 (i.e., the median of the distribution). Data points in each group are also divided into two conditions as deleterious and neutral SAVs. Counts for each group-condition combination (e.g., deleterious mutations in the significant polarity change group) are obtained and used in Fisher’s exact test to calculate p-values of associations between the magnitude of physicochemical change and the variant effect. The results considering all of the four physicochemical property types were found to be statistically significant at the confidence interval of 99%, with p-values 0, 0, 1.5×10^−146^ and 0 for polarity, volume, composition and the Grantham score, respectively. These results indicate physicochemical properties can be considered as good indicators of the functional effect of SAVs.

#### 3.1.3. Structural Location-Based Evaluation

Here, we aimed to observe if the location of the mutation on the protein structure contains information regarding its effect. For this, we grouped variation’s position on the sequence as core, interface or surface, according to the structural information and relative solvent accessible surface area measures (please refer to section 2.3.3 for more information). As observed in Figure 3b, mutations found in the core and interface regions have a higher deleteriousness rate (i.e., 72% and 69%, respectively) compared to the ones in the surface regions (i.e., 49%). We also tested this observation using Fisher’s exact test (taking into account that the number of deleterious mutations are higher than neutrals in the overall source dataset) and found the relationship between a mutation’s structural location as core, interface or surface is significantly related to being deleterious at 99% confidence interval with p-values of 0, 2.9×10^−141^ and 4.6×10^−127^, respectively. These results were expected since core regions are critical in terms of the stability of the protein and mutations may have a destabilizing effect leading to structural changes (Engin et al., 2015; Guo et al., 2004), whereas interface regions are important because they play roles in protein-protein interactions and a mutation at these regions may prevent the formation of a protein complex or a transient interaction, thus causing a deleterious effect (Engin et al., 2015; Nishi et al., 2013). Thus, our results are in correlation with the literature.

#### 3.1.4. Positional Sequence Annotation-Based Evaluation

Mutations in critical sites/regions in proteins (e.g., active sites, DNA binding regions, etc.) are generally more disrupting compared to the ones found in other regions. Information related to these important functional sites/regions can be incorporated into models using protein sequence annotations such as the positional annotations provided by UniProt (McGarvey et al., 2019). However, only a small number of proteins are associated with each type of positional annotation (Figure 4a). One of the possible reasons is that some of these annotation categories are protein family/class-specific, such as the active sites of enzymes, thus expected to be annotated to enzymes only. The second reason is the fact that annotations of proteins are incomplete, and further experimental and computational analyses are required to increase coverage. Nevertheless, proteins of only 201 SAV data points (out of 96,274) have no positional annotation at all in UniProtKB. On average, an annotation category is associated with the 24% of our dataset. Individual rates are shown in Figure 4a for each annotation type.

**Figure 4.**
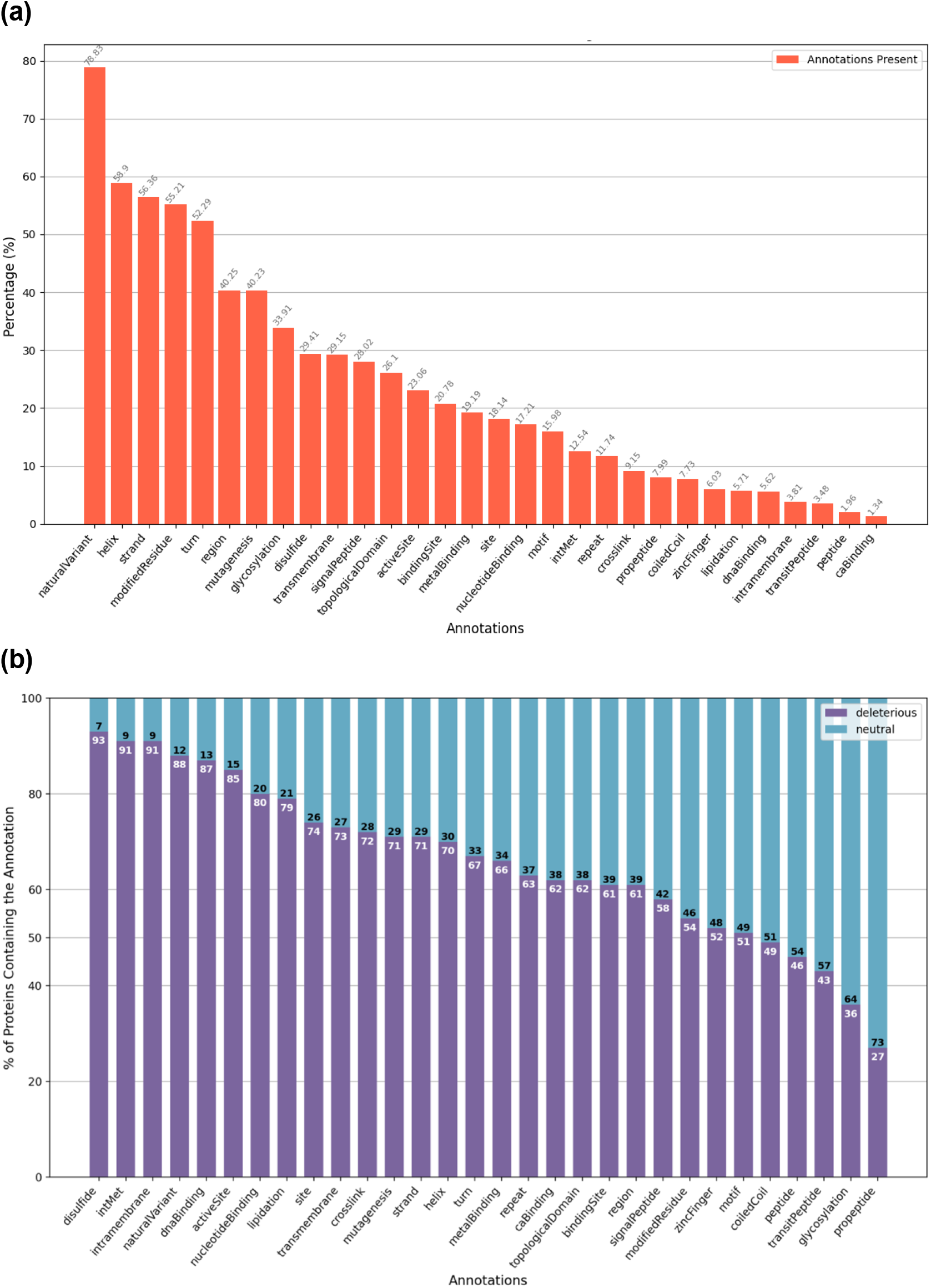
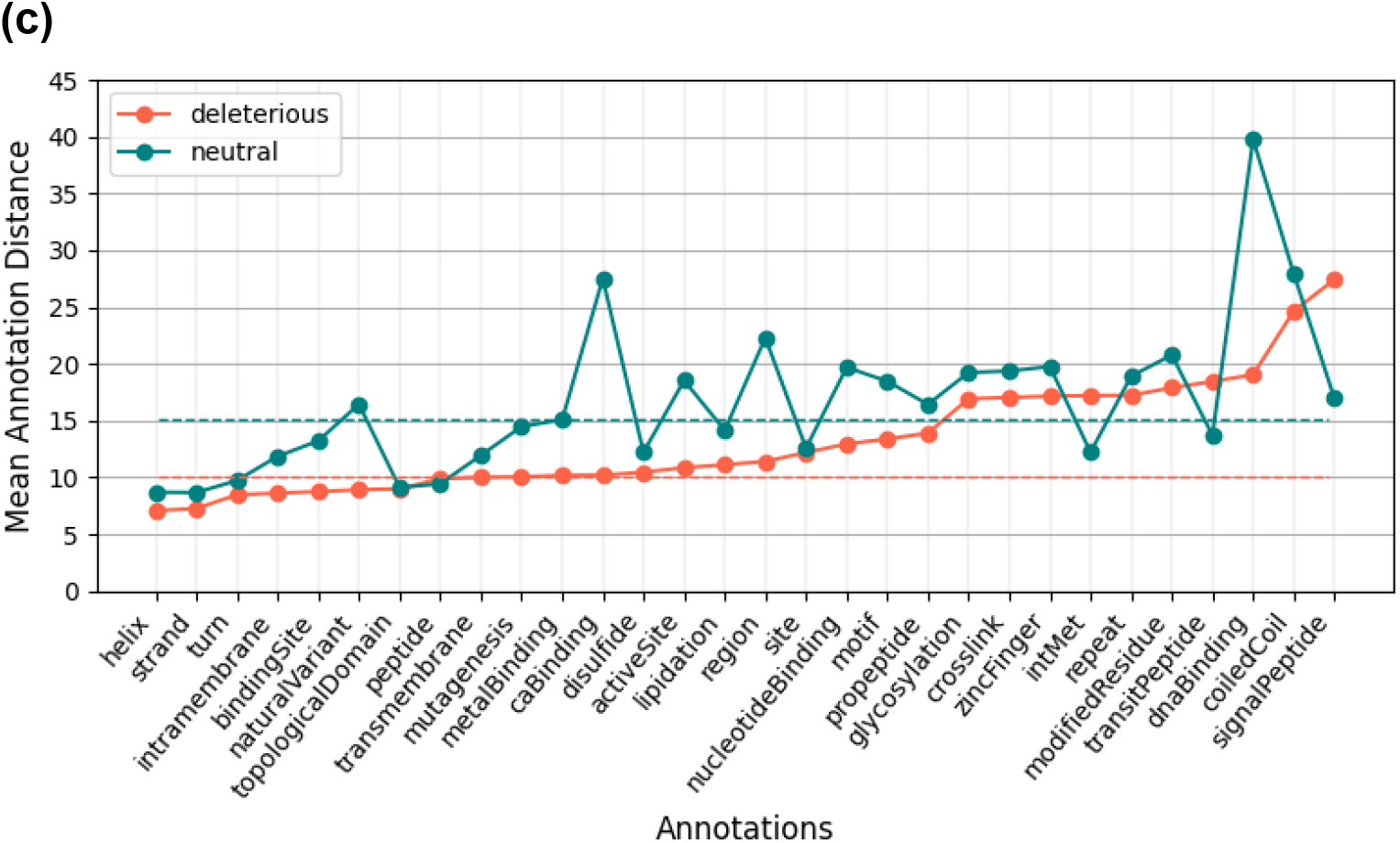
**(a)** The coverage of each annotation category on the proteins in our SAV dataset (e.g., nearly 23% of proteins in our dataset have at least one active site annotation in UniProtKB), **(b)** the rates of deleterious vs neutral variations for each annotation category, which are calculated considering the SAV data points in our dataset that coincide with a positional annotation on the same residue, **(c)** Mean spatial distances between annotated residues and mutated residues in the structure, calculated independently for each annotation category, considering the cases in which the SAV and annotated residue do not coincide on the same residue in the sequence. In the cases of the existence of multiple annotations on a protein, only the annotation that is located spatially closest to the mutation of interest is taken into account. Dashed lines indicate overall averages calculated by taking the mean of all distances. For the calculation of Euclidean distances between residues, the coordinates of C11 of the respective amino acids are extracted from PDB models.

With the aim of evaluating the effects of mutations in functional sites/regions, we extracted the percentage of deleterious and neutral mutations in our dataset that correspond to each of the 30 different positional annotation categories explained in Table 1. The results are displayed in Figure 4b, which indicates a prevalence of either being deleterious or neutral for most of the categories. Here, variations that coincide with 6 annotation categories (i.e., peptide, glycosylation, coiled-coil, propeptide, signal peptide, and transit peptide) have higher rates of neutral mutations; whereas, rates of deleterious mutations are higher for 23 categories. For the category called “natural variation sites”, the rates for neutral and deleterious SAVs are equal to each, since UniProt lists both disrupting and benign variations together under this category.

The results are as expected for the 23 categories that have higher rates of deleterious mutations since, for example, a mutation in the active site of an enzyme is highly likely to disrupt the enzymatic function, and thus, have an overall deleterious effect. Considering those 6 categories where the rate of neutrals is higher, we can infer that these sites/regions are not strongly related to the function of the mature protein. For example, coiled coils are dynamic and flexible regions, and they differ in terms of both length and variability, from being almost invariant to being hypervariable (Truebestein and Leonard, 2016). The flexible nature of coiled coils may explain the percentage of neutrality being higher in variations coinciding with these regions. Other such categories, e.g., propeptides, signal peptides, and transit peptides, function as recognition sites and for targeting proteins, which are cleaved during the maturation of the protein (Wu et al., 1990). SAVs usually cause a partial decrease in the efficiency of the recognition and targeting processes since the patterns themselves are variable. Thus, severe impacts are rarely observed on the overall protein function (Holbrook et al., 2016).

In our dataset, 67% of the SAV positions coincide with at least one positional annotation (excluding the “natural variant” category annotations), which means that we cannot utilize this information for 33% of the data points, to model and predict the effect. On the other hand, even if a variation does not directly correspond to an important site/region, ones that are located proximally to critical positions still tend to disrupt the intended function. For example, a mutation in the same pocket with the active site of an enzyme, where the mutated amino acid has significantly different properties from the wild-type, may have a deleterious effect on the enzymatic function. In order to take these cases into account, we decided to utilize spatial distances between the SAV and the positionally annotated sites/regions on the 3-D structure of the protein. To observe whether the distance data contain information relevant to the effects of variations, we calculated the average spatial distances between variations and each of the 30 positional annotation categories, considering the cases where SAVs and annotations are on the same protein, but do not coincide with each other on the same residue. We plotted the curves for both neutral and deleterious SAV data points in Figure 4c. Here, it is observed that, for most of the annotation categories, average distances are lower for deleterious variations compared to neutral ones, which indicate that the spatial distance data carry information that can be utilized for modeling and predicting the effects of SAVs.

#### 3.1.5. Visualization of the Variation Space

Visualizing high dimensional data in reduced dimensions provides a means for exploring the distribution of different properties of the samples in the dataset. One obvious visualization in our case would be to compare the neutral and deleterious SAVs. Another one would be a comparison between SAVs from different source databases. As mentioned previously, in this study, variation data has been gathered from three individual sources; ClinVar, UniProt and PMD. Each database differs in the way they evaluate and label SAVs. For instance, ClinVar and UniProt report the variants’ associations with diseases. On the other hand, PMD categorizes variations in terms of whether they impair the structure of the protein. With the aim of examining the distribution of data coming from these three databases, and between neutral and deleterious classes, we conducted a dimensionality reduction analysis using both Multidimensional Scaling (MDS) (Cox and Cox, 2008) and t-distributed stochastic neighbor embedding (t-SNE) (van der and Geoffrey Hinton, 2008) on our SAV dataset, and visualized the results on a 2-dimensional plane.

For both MDS and t-SNE embeddings, pairwise distances between pairs of variation data points were required. We could not directly calculate a vectorial distance on our feature vectors since they contain both categorical and real-valued dimensions. To address this issue, we calculated mean distance values using a simple heuristic, the details of which can be found in Supplementary Information S3. Figure 5 displays the output of MDS (panels a, b and c) and t-SNE (panels d, e and f) embeddings for randomly selected 9000 SAV data points from our dataset (4500 neutral and 4500 deleterious variations, and 3000 data points are coming from each source database, i.e., UniProt, ClinVar and PMD). It is possible to observe from Figure 5b and e that there is no clear separation between ClinVar and UniProt; however, PMD formed distinct clusters, probably due to the technical differences in categorizing variations in these databases. Also, data points from ClinVar are scattered throughout the 2-D plane, indicating a high heterogeneity in ClinVar data. Considering the consequence of variations, there is no clear distinction between neutral and deleterious conditions (Figure 5a and d). These results generally indicated that although there are cluster formations to a certain degree, it is not possible to separate neutral and deleterious variations from each other at reduced dimensions, and further analyses are required. A possible reason contributing to the heterogeneity would be the heuristic we employed to calculate pairwise distances between SAVs.

**Figure 5.**
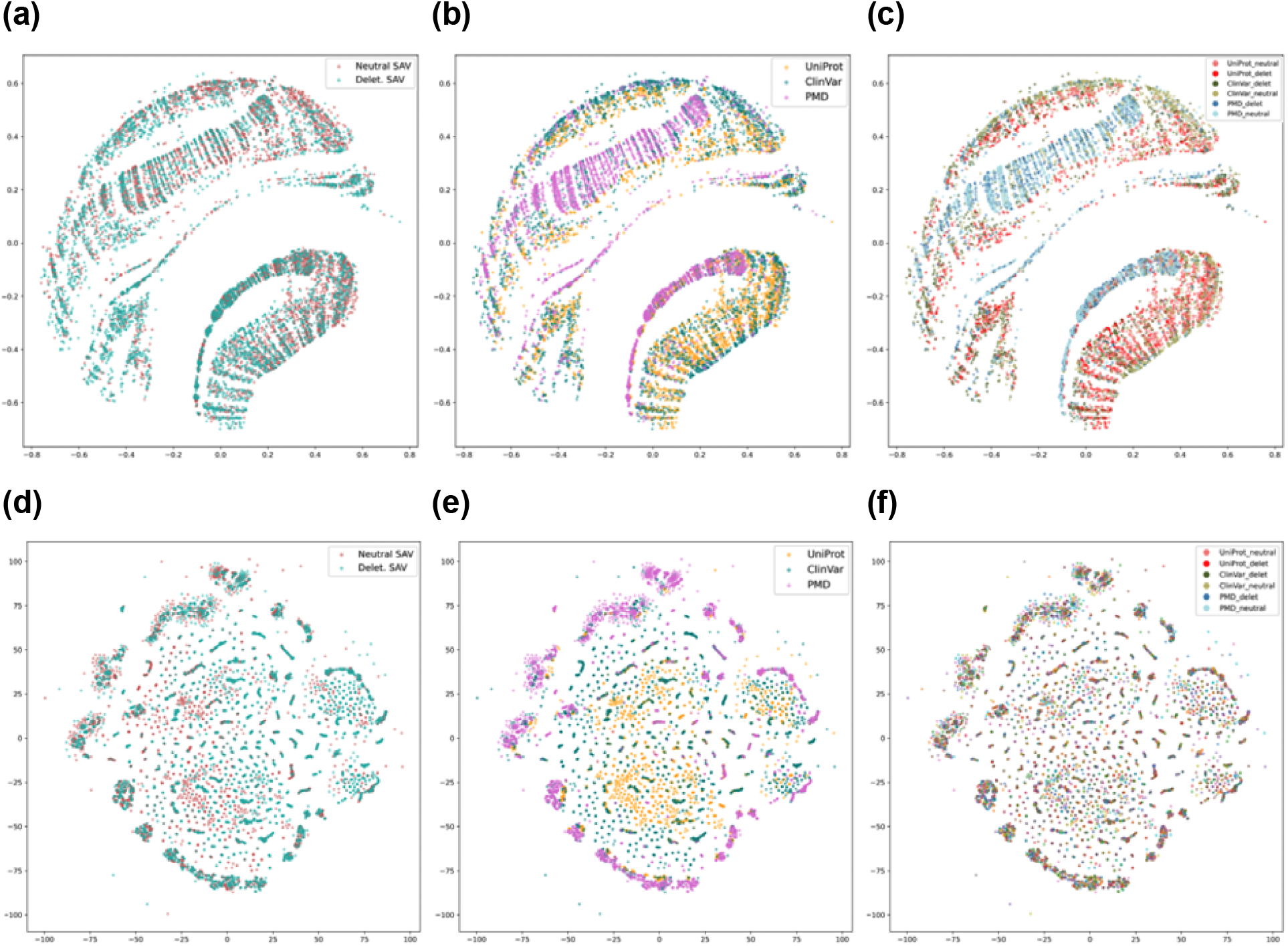
2-dimensional visualization of the variation dataset via MDS (a, b and c) and t-SNE (d, e and f) embeddings; **(a & d)** neutral vs. deleterious SAVs, **(b & e)** source dataset comparison, **(c & f)** data represented in 6 groups: ClinVar-neutral, ClinVar-deleterious, UniProt-neutral, UniProt-deleterious, PMD-neutral and PMD-deleterious.

### 3.2. Predicting the Effects of Variations via ASCARIS

In this section, we evaluate the discriminative power of our SAV feature vectors in terms of separating deleterious mutations from neutral ones, in the framework of machine learning-based modeling, considering the PDB and AlphaFold versions of ASCARIS independently. For this, we first carried out an investigative (ablation) analysis to observe the effect of features, datasets and algorithms. Afterwards, we trained and optimized a predictive model using our multi-source variation dataset. We then compared the performance of our model against the state-of-the-art variant effect predictors on different benchmark datasets. Finally, we conducted a use-case analysis on 2 SAVs, the effects of which are correctly predicted by ASCARIS.

#### 3.2.1. Ablation Study to Investigate Features, Data Sources and Classifiers

We evaluated the predictive power of different feature types in our ASCARIS-PDB and ASCARIS-AlphaFold SAV representations by generating feature vectors with different feature combinations, and training/validating a random forest classification prediction model with each version (using our combined SAV dataset of 96,274 data points). We then compared the performance results and calculated feature rankings to evaluate the importance of features. We measured the performance of all models via a 5-fold cross validation analysis by keeping the data points on each validation split the same between models, and using the metrics: AUROC, accuracy, precision, recall, F1-score, and finally MCC which is considered as the main metric due to imbalance between neutral and deleterious SAVs in the dataset.

Our first set of models (i.e., p1 and a1) only contained domain annotation information, which is incorporated in the form of identifiers of InterPro domains where the SAV of interest resides in the protein. This model incorporates domain annotations considering 307 domains that were found to be statistically significant in terms of separating neutral and deleterious SAVs from each other in Fisher’s exact test analysis (please see Methods subsection 2.3.1 and Supplementary Information S2). Feature vectors of this model are composed of a single dimensional variable that contains 308 categories (i.e., 307 significant domains displayed in Table S2 and an additional category to accommodate the rest of the domains and the “no domain hit” cases). Our second set of models (i.e., p2 and a2) incorporates physicochemical properties by generating 4-dimensional feature vectors containing real-valued polarity, composition, volume and Grantham scores. Our third set of models (i.e., p3 and a3) is composed of 2 dimensions related to the location of the SAV on the protein sequence, (1) the solvent accessible surface area value and (2) its categorization as “core”, “surface” or “interface”.

Performance comparison between the first 3 PDB-based models (Table 2a) indicated that physicochemical features are notable indicators for variant effect prediction together with domains (MCC: 0.26 and 0.24, respectively). These findings are in accordance with the literature as previous studies also highlighted the importance of physicochemical features for variant effect prediction (Stone and Sidow, 2005) and usefulness of domain annotations in modeling functions (Doğan et al., 2016) and ligand interactions (Doğan et al., 2021-I) of proteins. As our fourth set of models, we integrated features of the first 3 models and trained a new one with these integrated features, which resulted in a significant performance increase (MCC: 0.42 and 0.48 for p4 and a4, respectively), indicating their complementarity. In the fifth set of models, we utilized positional feature/sequence annotations in terms of one-to-one correspondence between the annotated positions and SAVs on the sequence. This binary 30-dimensional feature vector resulted in a high performance (MCC: 0.56 and 0.54 for p5 and a5, respectively) which points out to the effectiveness of this approach, which also happens to be the main contribution of our study to the literature. Furthermore, adding 30 more dimensions corresponding to the spatial distances between the annotated residues and the SAV of interest (as our 6th set of models) further increased the performance (MCC:0.59 and 0.61 for p6 and a6, respectively). In our 8th and final set of models, we incorporated all features in 68 dimensions, which displayed the best performance in terms of all metrics (MCC: 0.61 and 0.63 for p8 and a8, respectively). Based on the fact that the maximum predictive performance has been achieved by the model that incorporates all types of features, we decided to construct our finalized variant effect prediction model using all features.

**Table 2.**
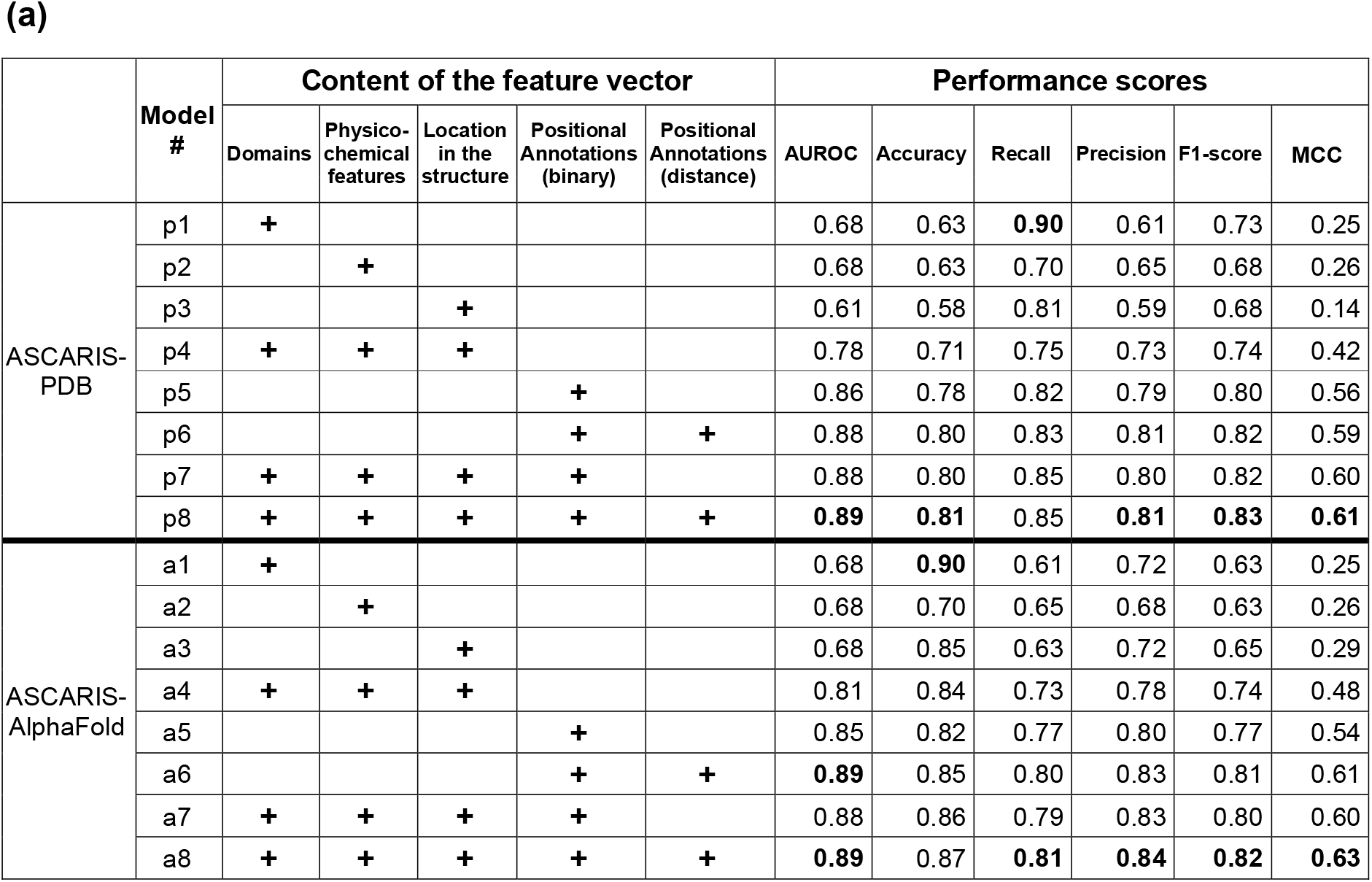
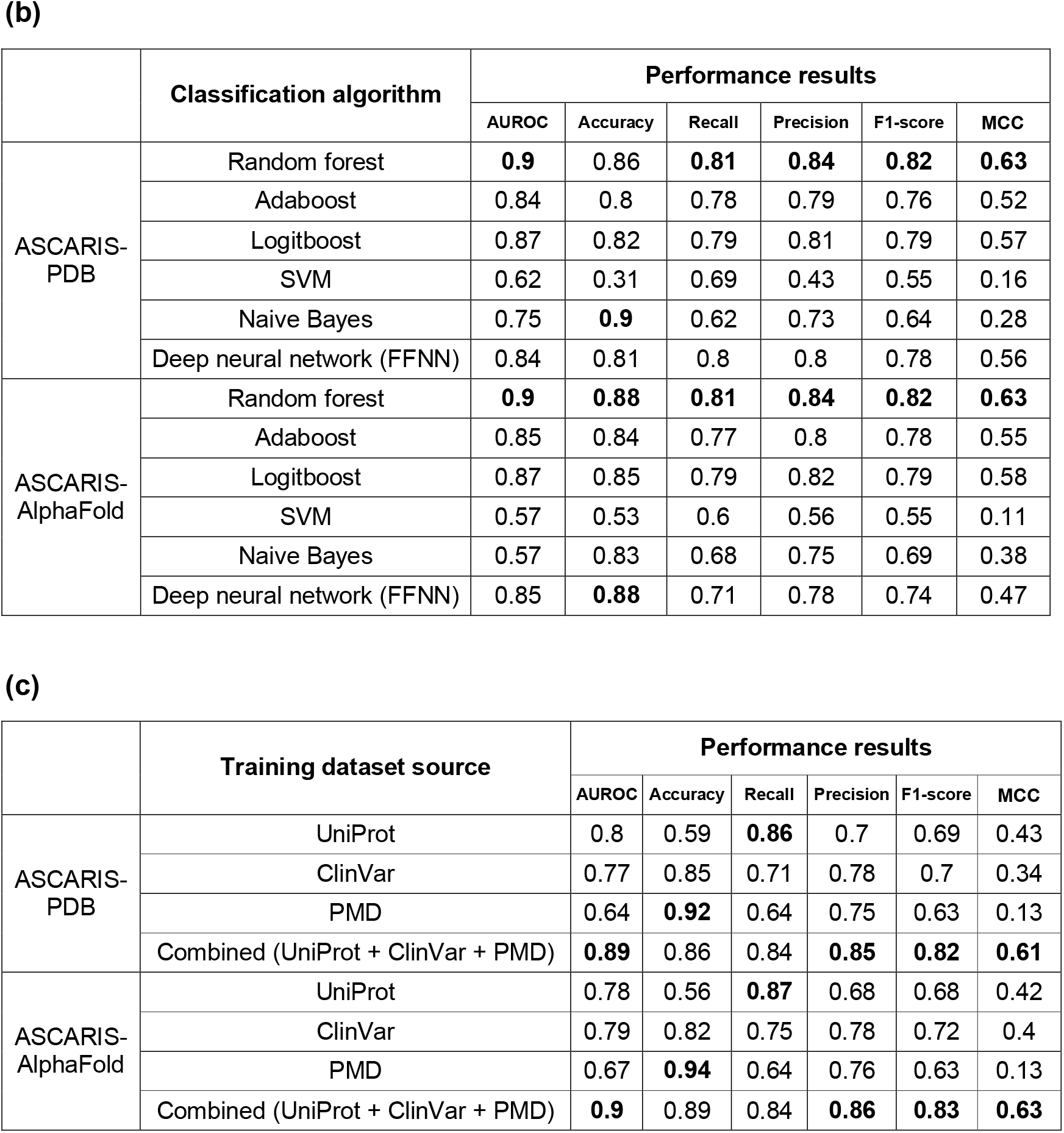
Performance results of the ablation study for PDB and AlphaFold versions of ASCARIS, where; **(a)** different versions of the feature vector are tested (i.e., different combinations of physicochemical features, significant domains, location on the structure and positional sequence annotations) via Random Forest binary classifier and the combined training data, and **(b)** different classification algorithms are evaluated (i.e., random forest, adaboost, logitboost, SVM and naive Bayes) using the full-sized feature vector that contains all types of features and the combined training data; **(c)** different training data resources are analyzed (i.e. UniProt, ClinVar, PMD and the full dataset which is the combination of three) by benchmarking on the same hold-out test dataset.

With the aim of selecting the classification algorithm, we performed a cross-validation based analysis to compare the performance of random forest classifier (RF), Adaptive boosting (Adaboost), adaptive logistic regression (LogitBoost), support vector machine (SVM), and the naive Bayes (NBayes). Widely accepted and default hyperparameter values were used for all classifiers, such as the maximum number of decision splits: n-1, and the number of predictors to select at random for each split: √n (n represents the number of predictors/features), the number of ensemble learning cycles: 100, minimum number of leaf node observations: 1 for all ensemble-based methods, Learning rate: 1 for Adaboost and LogitBoost, kernel function: linear and kernel scale: 1 for SVM, and distribution type for categorical and real values predictors: multivariate multinomial and Gaussian, respectively. The results of the 5-fold cross-validation test is given in Table 2b, which indicated that RF is the most successful classifier. With the observation of these results, we decided to base our predictor on the RF algorithm. In this analysis, we did not prefer artificial (deep) neural networks due to the requirement of further feature encodings for our categorical variables (e.g., domains, structural location of variation, direct positional correspondence with sequence annotations), and possible inconsistencies when those encodings are used together with our real-valued variables (e.g., spatial distances between the variation and different types of positional annotations).

Finally, we performed an analysis to observe the performance of the models trained on SAV datasets from individual data sources; i.e., ClinVar, UniProt and PMD, to observe which source yields a higher generalization power to the model, and evaluate whether combining data from 3 different SAV resources under one model, or using only one of the data sources, is the better approach for training our final predictive model. We prepared one test dataset composed of 7,694 SAVs (8% of the whole combined dataset) by taking a nearly equal number of data points from each data source, and used it as a hold-out test dataset to calculate the performance of all models. Three training datasets, each composed of SAV data points from an individual data resource and containing 20,876 SAVs (the size of the smallest individual SAV dataset), have been prepared. SAVs in the test dataset have already been excluded from these training datasets. Three predictive models were trained with these datasets. For all of these models, the finalized full set of features were incorporated into SAV representation vectors. Table 2c shows the performance results of these three individual data resource models, along with the model that utilized the combined training dataset. According to MCC scores, the combined dataset model provided the best performances with 0.61 and 0.63 for PDB and AlphaFold versions of ASCARIS, respectively (Table 2c). In terms of individual-dataset models, the UniProt dataset led to the highest performances (MCC: 0.43 and 0.42) followed by ClinVar (MCC: 0.34 and 0.40) and PMD (MCC: 0.13 and 0.13). This could be due to the way SAVs are classified in these databases (i.e., UniProt and ClinVar focus on the reported effect in terms of associations with diseases, whereas PMD focuses on effects related to protein’s structural stability). Since the performance of the model using the combined dataset is the best, we based our predictive method on this training dataset and used it further in this study.

An interesting observation here is that ASCARIS-AlphaFold generally performs slightly better compared to the PDB version. We believe the reason behind is not related to the quality of the models but their coverage, since AlphaFold provides full structural coverage over the entire protein sequences, whereas in the PDB version, less than half of the SAVs and positional annotations could be mapped to regions covered by a PDB model, and the rest of them were resolved using SwissModel or MODBASE models (Figure 2a).

#### 3.2.2. Training, Validation and Evaluation of the Finalized Model

We built our finalized variant effect predictor model using all types of variables in feature vectors (with significant domains), the merged training dataset from all three sources, and the RF algorithm. We employed a grid-search-based hyper-parameter optimization test via 5-fold cross-validation. The evaluated hyper-parameter types and their respective values are explained in section 2.4. The selected values at the end of the optimization process are; number of trees: 300, maximum number of decision splits: 96273 (size of the whole dataset - 1), and number of predictors to select at random for each split: 8. The detailed results of hyper-parameter optimization tests can be found in Table S4. We measured the final performance of our model on a hold-out test dataset, which corresponds to 10% of the data points in our original dataset. According to the results of this analysis, our ASCARIS-PDB and ASCARIS-AlphaFold models perform with; AUROC: 0.89 and 0.90, accuracy: 0.81 and 0.82, recall: 0.85 and 0.88, precision: 0.81 and 0.81, F1-score: 0.83 and 0.84, and MCC: 0.62 and 0.64, respectively.

The feature importance ranking of the optimized models shows that “significant domains”, solvent accessible surface area and physicochemical features are the most critically important determinants of the decision process (Figure S3). Positional annotations vary in their importance ranking; however, it is important to note that these annotations are scarce, thus, the missing information may cause some of the annotation categories to rank lower. Among them, spatial distance to previously reported mutagenesis sites is the most critical. It is also important to note that variations recorded in natural variant and mutagenesis variables in our feature vectors do not correspond to the SAV data points in our variant effect prediction model training/validation/test dataset. As a result, there is no data/information leak from training to test.

Mainly due to the simplicity of our feature vectors, the proposed model had convenient run times, such that, training of the full model with the optimal parameters took 91 seconds, running the whole hyper-parameter optimization analysis with 5-fold cross-validation and grid search (64 hyperparameter values sets * 5 folds = 320 training/validation runs) required 7 hours, and predicting the effects of 10,000 SAVs in the hold-out test dataset (with the pre-trained model) took 3 seconds on an 8-core 2.3 MHz Intel i9 CPU with 16 GB memory.

#### 3.2.3 Performance Comparison with the State-of-the-art

In order to compare our model to the state-of-the-art methods, we performed four different benchmark analyses against widely-used variant effect prediction tools.

In the study by Schwarz et al., authors compared the performance of their method, MutationTaster2 (Schwarz et al., 2014), to that of SIFT (Ng and Henikoff, 2003), PROVEAN (Choi and Chan, 2015) and two different versions of PolyPhen-2 (Adzhubei et al., 2013) on multiple benchmark datasets. Here, we used the main benchmark test dataset from Schwarz et al., that contains 2600 variation data points from ClinVar and the 1000 Genomes project (1000 Genomes Project Consortium, 2015). We created feature vectors for the variation data points in the *MutationTaster* dataset. To yield a fair comparison, we filtered our training dataset by first, removing the hold-out test data points, and second, by removing all SAV data points that entered our source databases (i.e., UniProt, ClinVar and PMD) at and after the year 2014, so that our training data would be temporally consistent with the training datasets used in Schwarz et al., We re-trained our RF model using default hyper-parameter values. The results are displayed in Table 3a, where two versions of ASCARIS (i.e., PDB and AlphaFold) are shown together with methods from the literature. ASCARIS-AlphaFold was among the top three in terms of accuracy and F1-score (after MutationTaster2) and the best in terms of precision and specificity. The predictive performance of all competing methods on this test dataset are considerably high, which decreases the capacity for discriminating competing methods. To address this issue, we constructed a challenging sub-set by selecting SAV data points that are correctly predicted by half or less of the competing methods (i.e., < 4 methods), including ours (for fair comparison). Then we used this challenging sub-set, which is composed of 167 SAVs (74 neutral and 93 deleterious), as our hold-out test set and calculated the performance metrics of all methods. These challenging SAVs are provided in Table S5. According to performance results in Table 3b, our method (ASCARIS-AlphaFold) was the best performer, followed by MutationTester2, which indicates that our approach performs well on challenging/difficult cases. The reason behind observing low performance values here is deliberately selecting mostly inaccurately predicted data points in this analysis. To observe whether our method produces complementary results to others, we analyzed prediction similarities/intersections among methods. Figure S2 displays the number of intersecting predictions on this challenging dataset, in terms of neutral SAVs and deleterious SAVs via Venn diagrams (in panels a and b, respectively) (Heberle et al., 2015). As shown, our method has the highest number of distinct predictions in this benchmark, especially for neutral SAVs. Intersections among the state-of-the-art methods are much higher compared to intersections between our method and the state-of-the-art methods, indicating the value of the proposed approach in terms of complementing the widely-used alignment and/or structure-based methods. This can be attributed to our annotation-based featurization approach, as it is marginally different from the widely-used state-of-the-art methods included in this analysis. Due to the elevated performance of the AlphaFold version of ASCARIS compared to the PDB version, we continued the remaining benchmarking analysis only with ASCARIS-AlphaFold.

**Table 3.**
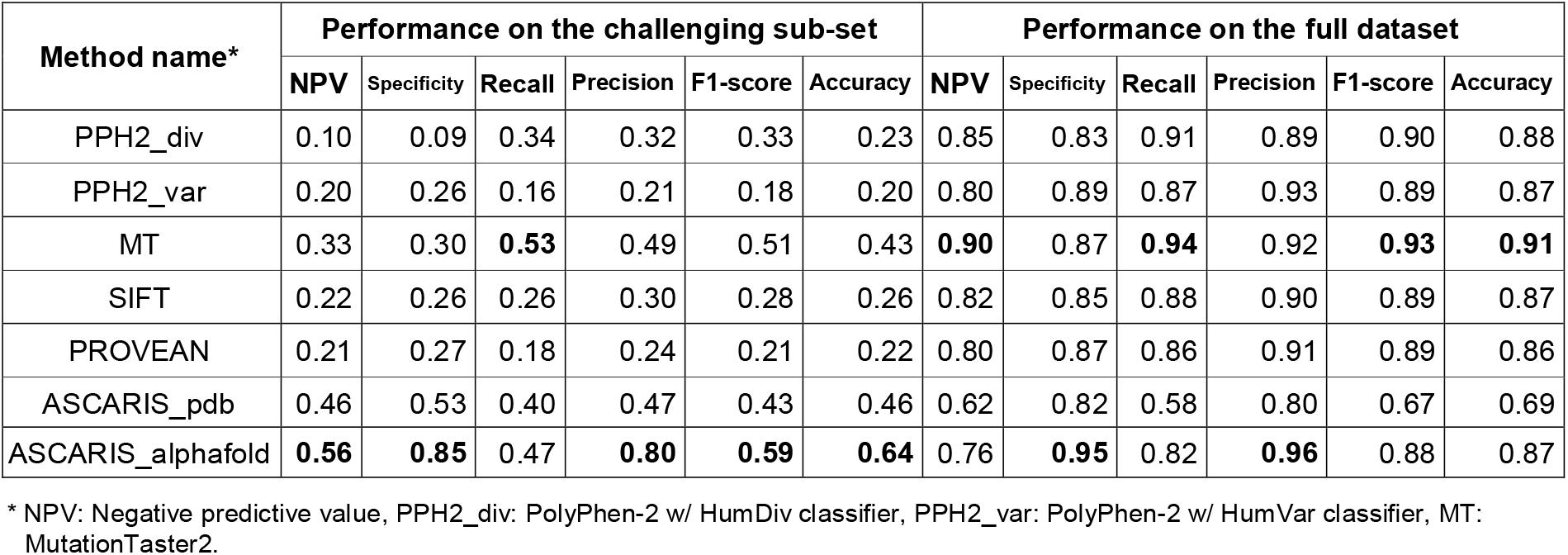
Variant effect prediction performance comparison on the MutationTaster dataset on the full dataset (benchmark 1) and its challenging sub-set. Best performances are shown in bold font for each metric.

| Method name* | Performance on the challenging sub-set |  |  |  |  |  | Performance on the full dataset |  |  |  |  |  |
| --- | --- | --- | --- | --- | --- | --- | --- | --- | --- | --- | --- | --- |
|  | NPV | Specificity | Recall | Precision | F1-score | Accuracy | NPV | Specificity | Recall | Precision | F1-score | Accuracy |
| PPH2_div | 0.10 | 0.09 | 0.34 | 0.32 | 0.33 | 0.23 | 0.85 | 0.83 | 0.91 | 0.89 | 0.90 | 0.88 |
| PPH2_var | 0.20 | 0.26 | 0.16 | 0.21 | 0.18 | 0.20 | 0.80 | 0.89 | 0.87 | 0.93 | 0.89 | 0.87 |
| MT | 0.33 | 0.30 | **0.53** | 0.49 | 0.51 | 0.43 | **0.90** | 0.87 | **0.94** | 0.92 | **0.93** | **0.91** |
| SIFT | 0.22 | 0.26 | 0.26 | 0.30 | 0.28 | 0.26 | 0.82 | 0.85 | 0.88 | 0.90 | 0.89 | 0.87 |
| PROVEAN | 0.21 | 0.27 | 0.18 | 0.24 | 0.21 | 0.22 | 0.80 | 0.87 | 0.86 | 0.91 | 0.89 | 0.86 |
| ASCARIS_pdb | 0.46 | 0.53 | 0.40 | 0.47 | 0.43 | 0.46 | 0.62 | 0.82 | 0.58 | 0.80 | 0.67 | 0.69 |
| ASCARIS_alphafold | **0.56** | **0.85** | 0.47 | **0.80** | **0.59** | **0.64** | 0.76 | **0.95** | 0.82 | **0.96** | 0.88 | 0.87 |

\* NPV: Negative predictive value, PPH2\_div: PolyPhen-2 w/ HumDiv classifier, PPH2\_var: PolyPhen-2 w/ HumVar classifier, MT: MutationTaster2.

The remaining benchmark datasets were obtained from the study by Grimm et al., (Grimm et al., 2015) where authors conducted a comparative study to assess various tools. We used 3 datasets from this benchmark (i.e., *predictSNP, VariBench* and *SwissVar*) to compare our method against predictors from the literature. More information regarding these datasets can be found in Grimm et al. We again removed both the entire test dataset and SAVs that entered our source databases after the year 2014 to yield a fair comparison. The performance metrics that were used in the original study have also been incorporated here. Performances are calculated using a sub-set of the datasets composed of SAVs where all competing methods provided a prediction, to yield a fair comparison.

The performance results on *predictSNP* dataset (Bendl et al., 2014) (Table 4a) displays that our method is the best performer in terms of precision, accuracy, F1-score and MCC, indicating the effectiveness of our featurization approach. According to results in Table 4b, ASCARIS-AlphaFold was the best performer together with Condel+ (González-Pérez and López-Bigas, 2011) on the *VariBench* dataset (Sasidharan Nair and Vihinen, 2013; Thusberg et al., 2011) in terms of MCC. Condel+ is an ensemble predictor combining the tools: PP2 (Adzhubei et al., 2010), SIFT (Ng and Henikoff, 2003), MASS (Reva et al., 2011) and FatHMM-W (Shihab et al., 2013). Finally, the results on the last dataset, which consists of a selection of SAVs from the SwissVar database (Mottaz et al., 2010), showed that our method was the fourth best performer in terms of MCC, after Logit, Logit+ and Condel+ (Table 4c). In all benchmark analyses, our method displayed considerably high performance and competed well with the widely used methods in the literature, even though variant effect prediction is not the main purpose of ASCARIS. This indicates that our approach has the ability to represent SAVs regarding their functional consequences.

**Table 4.** Variant effect prediction performance comparison on the challenging sub-set of: **(a)** the *predictSNPSelected* dataset (benchmark 2), **(b)** *VariBenchSelected* dataset (benchmark 3), **(c)** *SwissVarSelected* dataset (benchmark 4). Best performances are shown in bold font for each metric.

**(a)**
| Method name | NPV | Specificity | Recall | Precision | F1-score | Accuracy | MCC |
| --- | --- | --- | --- | --- | --- | --- | --- |
| Logit | 0.81 | 0.75 | 0.81 | 0.75 | 0.78 | 0.78 | 0.56 |
| Logit+ | 0.96 | **0.86** | 0.92 | 0.71 | 0.80 | 0.87 | 0.72 |
| Condel | 0.90 | 0.48 | **0.95** | 0.64 | 0.77 | 0.71 | 0.48 |
| Condel+ | 0.92 | 0.72 | 0.93 | 0.76 | 0.84 | 0.82 | 0.67 |
| FatHMM-W | 0.89 | 0.82 | 0.89 | **0.82** | 0.85 | 0.85 | 0.71 |
| FatHMM-U | 0.72 | 0.75 | 0.68 | 0.71 | 0.70 | 0.72 | 0.43 |
| LRT | 0.77 | 0.57 | 0.83 | 0.66 | 0.74 | 0.70 | 0.42 |
| SIFT | 0.81 | 0.68 | 0.84 | 0.72 | 0.77 | 0.76 | 0.52 |
| MutationTaster2 | **0.98** | 0.60 | 0.79 | 0.12 | 0.21 | 0.61 | 0.20 |
| MutationAssessor | 0.79 | 0.73 | 0.80 | 0.73 | 0.76 | 0.76 | 0.52 |
| PolyPhen2 | 0.84 | 0.66 | 0.87 | 0.70 | 0.78 | 0.76 | 0.54 |
| ASCARIS_alphafold | 0.91 | 0.81 | 0.91 | **0.82** | **0.86** | **0.86** | **0.73** |

(b)
| Method name | NPV | Specificity | Recall | Precision | F1-score | Accuracy | MCC |
| --- | --- | --- | --- | --- | --- | --- | --- |
| Logit | 0.61 | 0.73 | 0.77 | 0.85 | 0.81 | 0.76 | 0.48 |
| Logit+ | 0.75 | 0.68 | 0.74 | 0.67 | 0.70 | 0.71 | 0.42 |
| Condel | 0.74 | 0.42 | 0.93 | 0.79 | 0.85 | 0.78 | 0.43 |
| Condel+ | **0.76** | 0.58 | 0.92 | 0.83 | **0.87** | **0.81** | **0.54** |
| FatHMM-W | 0.67 | 0.63 | 0.87 | 0.85 | 0.86 | 0.80 | 0.51 |
| FatHMM-U | 0.53 | **0.76** | 0.72 | **0.88** | 0.79 | 0.73 | 0.44 |
| LRT | 0.59 | 0.63 | 0.82 | 0.84 | 0.83 | 0.77 | 0.44 |
| SIFT | 0.61 | 0.62 | 0.83 | 0.83 | 0.83 | 0.76 | 0.44 |
| MutationTaster2 | 0.71 | 0.55 | **0.90** | 0.82 | 0.86 | 0.79 | 0.49 |
| MutationAssessor | 0.53 | 0.69 | 0.74 | 0.85 | 0.79 | 0.73 | 0.41 |
| PolyPhen2 | 0.67 | 0.66 | 0.86 | 0.85 | 0.86 | 0.80 | 0.52 |
| ASCARIS_alphaFold | 0.66 | 0.71 | 0.84 | 0.87 | 0.86 | 0.80 | **0.54** |

(c)
| Method name | NPV | Specificity | Recall | Precision | F1-score | Accuracy | MCC |
| --- | --- | --- | --- | --- | --- | --- | --- |
| Logit | 0.90 | 0.80 | 0.74 | 0.57 | 0.64 | 0.78 | 0.50 |
| Logit+ | 0.91 | **0.86** | 0.78 | **0.66** | **0.71** | **0.84** | **0.60** |
| Condel | 0.93 | 0.45 | **0.90** | 0.38 | 0.54 | 0.57 | 0.33 |
| Condel+ | 0.91 | 0.71 | 0.80 | 0.50 | 0.61 | 0.73 | 0.46 |
| FatHMM-W | 0.84 | 0.80 | 0.58 | 0.51 | 0.54 | 0.74 | 0.37 |
| FatHMM-U | 0.85 | 0.75 | 0.63 | 0.48 | 0.54 | 0.72 | 0.35 |
| LRT | 0.90 | 0.59 | 0.82 | 0.44 | 0.57 | 0.66 | 0.37 |
| SIFT | 0.87 | 0.66 | 0.74 | 0.44 | 0.55 | 0.68 | 0.35 |
| MutationTaster2 | **0.94** | 0.56 | **0.90** | 0.43 | 0.58 | 0.65 | 0.41 |
| MutationAssessor | 0.87 | 0.69 | 0.72 | 0.46 | 0.56 | 0.70 | 0.36 |
| PolyPhen2 | 0.92 | 0.67 | 0.83 | 0.47 | 0.60 | 0.71 | 0.44 |
| ASCARIS_alphaFold | 0.88 | 0.77 | 0.70 | 0.53 | 0.60 | 0.75 | 0.44 |

#### 3.2.4 Use-case analyses

To evaluate a few examples where the proposed method could successfully detect the variant effect where the other predictors failed, we selected 3 example SAV data points (2 deleterious/disrupting and 1 neutral), from benchmark 1 and examined the corresponding SAV representation vectors together with relevant information from literature.

A SAV of the human Arylsulfatase B protein (ARSB_HUMAN) is selected as the first example (gene name: ASB or ARSB, protein UniProt accession: P15848, variation: C405Y, consequence: deleterious). ASB (or ARSB) is an enzyme (*N*-acetylgalactosamine-4-sulfatase) that removes the 4-sulfate group from chondroitin-4-sulfate (C4S) and regulates its degradation (Sharma et al., 2013). Mutations in the N-acetylgalactosamine-4-sulfatase gene cause reduced enzyme activity, and ASB deficiency is reported to be the cause of Mucopolysaccharidosis type VI (MPS VI; Maroteaux-Lamy syndrome) which is a lysosomal storage disorder (Bhattacharyya and Tobacman, 2009). Disease onset and rate of progression is variable for this disorder depending on the mutation, thus assessing mutational characteristics is critical. C405Y mutation changes cysteine in the 405^th^ position to tyrosine and disrupts the disulfide bond between the residues 405 and 447.

Theoretically, such inability to form a disulfide bridge that was present in the native form will cause the destabilization of the protein. According to a study by Karageorgos et al., (2007), where the authors analyze mutations from 105 patients with Mucopolysaccharidosis Type VI, and they reported that the C405Y mutation causes a slowly progressing disease with a late onset (Karageorgos et al., 2007). They suggest that this effect might be observed due to the destabilization introduced by the breaking of the disulfide bond. ASB’s active site involves at least 10 residues, and mutations around this region are expected to be highly critical. It is possible to observe from the visualization of the structure of the ASB protein that the mutation is located in a relatively close proximity with the active site (i.e., minimum spatial distance: 11.44 A), which may explain the effect suggested by the authors (Figure S5a). It is also reported that this mutation is within the annotated region of the superfamily “Alkaline-phosphatase-like, core domain superfamily” [InterPro ID: IPR017850], which is found to be heavily associated with deleterious/disrupting mutations (deleterious and neutral occurrences in our dataset for this domain are 722 and 112, respectively). To sum up, it is possible to state that the information included in our mutation feature vector including; (i) the direct correspondence of this mutation with a disulfide bond, (ii) close proximity to the active site of the protein, and (iii) the domain/family region in which the mutation occurred, may contribute to the correct classification of this mutation as deleterious/disrupting by our method.

Another example of a deleterious/disrupting case involves galactose-1-phosphate uridylyltransferase (gene name: GALT, protein UniProt accession: P07902, variation: Q38P, consequence: deleterious). This enzyme plays an important role in galactose metabolism. Enzymes, including GALT, in the Leloir pathway process galactose by converting it into glucose for glycolysis. Additionally, these enzymes are involved in inter-conversion of uridine diphosphate (UDP) hexoses that are used in the formation of glycogen and glycoconjugates, and the metabolism of UDP-*N*-acetyl-hexose-amines, and they are important structural elements of glycosaminoglycans (Tang et al., 2012). Mutations in these enzymes may result in autosomal recessive, classic galactosemia. Although early treatment with a strict diet can treat most symptoms, older patients develop neurocognitive disabilities and ovarian failure (McCorvie et al., 2016). Although it is only found to hit a turn region in terms of secondary structural elements, the Q38P mutation falls close to the metal binding site of the protein (i.e., Zn binding residues at positions 319 and 321) with a distance of 5.72 A (Figure S5b). It has been suggested that zinc binding influences the stability and aggregation tendency of the human GALT protein (McCorvie et al., 2016). It is also reported that changes in metal binding are among major factors causing destabilization and misfolding in this protein. Given that the mutation is in close proximity with the mentioned regions, it may contribute to the destabilization of the protein. Additionally, this mutation is reported to fall into the boundary of the structural HIT-like domain / superfamily of the GALT protein [InterPro IDs of the related domain and superfamily: IPR011146 and IPR036265] (out of 200 reported mutations that fall into this domain, 192 are implicated with a deleterious/disrupting label). Based on these properties, our method correctly predicted the effect of this mutation. Similar to the previous case, this example indicates the potential and importance of residue level annotation-centric evaluation of the effect of mutations.

A SAV of the human olfactory receptor 2W1 (OR2W1_HUMAN) is selected as the third example (protein UniProt accession: Q9Y3N9, variation: D296N, consequence: neutral). Olfactory receptors (ORs) are members of the G proteinccoupled receptor (GPCR) family and made up of seven transmembrane regions (Oh, 2021). This SAV is presented in UniProt’s humsavar dataset and reported as a polymorphism (a neutral variation) with dbSNP ID rs35771565. Upon taking a closer look at the protein sequence, it can be observed that the mutation is found in the extracellular portion of the protein, within the region annotated as topological domain. Also, the variation is found to be at a considerable distance from annotated sites/regions such as disulfide bond, transmembrane regions, and glycosylation sites. Due to this, these annotated sites/regions are probably not disrupted by the variation, and thus the function of the protein is not significantly impaired. Also, the change is between two hydrophilic amino acids, and this probably helps the protein to retain its original hydrophilic affinity on the surface. The position of the SAV also remains out of the annotated domain (InterPro id: IPR017452, domain name: GPCR, rhodopsin-like, 7TM) region of this protein. Based on the consequence of SAVs that have similar structural and annotation-centric characteristics, our method successfully predicted a correct neutral outcome for this SAV.

### 3.3. SAV Representation Construction Tool

We developed an open access command line tool for ASCARIS that generates the 74-dimensional SAV representations proposed in this study. The input to the tool is a file composed of one or more SAV data points (one in each line), composed of UniProt accession of the protein containing the SAV of interest, one letter notation of the wild type residue of interest, position of the mutated residue, and one letter notation of the mutated residue of interest in different columns, in tab delimited format. The output is again a tab delimited file containing the representation of each input SAV on a different row. Each row contains meta-data related to the SAV along with the actual 68-dimensions of the numerical features. The detailed explanation of the output file is provided in Table S1. The tool is available at https://github.com/HUBioDataLab/ASCARIS together with all datasets, instructions, and dependencies.

## 4. Conclusion

In this study, we developed a methodology, ASCARIS, to quantitatively represent single amino acid variations in proteins in terms of their spatial organization with positional sequence features, to reflect their functional characteristics. Our representations also include structure-derived information regarding physicochemical changes caused by the amino acid change, its location on the protein structure and its domain correspondence, constituting a 74-dimensional representation that can be utilized in any statistical data model to represent SAVs in a function-centric way. Possible applications can be predicting the effect of variations, omics-based modeling of cells or patients for precision medicine, designing new proteins, and many more.

As an application of our method, we trained ML models to predict consequences of single amino acid variations on protein functionality and compared their performance against well-known predictors from the literature. During this application, our initial idea was that, when used in the modeling alone, these representations could not compete with alignment-based variant effect predictors as it quantitatively describes SAVs from a limited perspective. However, we expected that it could produce complementary results as its point of view is different from existing methods. Results indicated that our method actually produces complementary results to conventional variant effect predictors. Moreover, it performs quite well on challenging cases where these methods mostly fail. Another interesting observation was that, the AlphaFold version of ASCARIS scored a higher performance compared to the PDB version of the method, indicating the benefit of having high quality structure predictions with almost complete coverage on sequences.

One of the advantages of our method is being practical as it utilizes documented annotations instead of trying to detect conservation via sequence alignments. Also, since the annotations are curated, the noise in data is expected to be low. Another advantage of our method is the interpretability of results when it is used in ML-based modeling, given that each dimension on our feature vectors have a known molecular/structural/functional correspondence. One limitation of ASCARIS is that the curated feature annotations of proteins are far from being complete, which means that we are dealing with missing information during modeling. Our method’s representation power will further increase with the addition of new protein feature annotations in databases such as UniProt. As future work, we plan to construct ensemble-based variant representations by integrating successful structure and alignment-based approaches with our method using multi-modal deep learning. We also plan to incorporate ASCARIS representations in large-scale biomedical knowledge graphs (Doğan et al., 2021-II) as variant feature vectors for integrative modeling of heterogeneous biomedical data via deep graph learning. We hope that these comprehensive SAV representations will be effectively utilized for data-centric modeling in various areas of biomedicine and biotechnology.

## Acknowledgement

Authors thank Dr. Nurcan Tuncbag (faculty member, Koc University, Turkey) for valuable discussions and insight regarding protein structure analysis in the context of evaluating single amino acid variations.

## Data availability

All of the datasets, results and the source code of this project are available at: https://github.com/HUBioDataLab/ASCARIS.

## References

Aboderin, A.A. An empirical hydrophobicity scale for α-amino-acids and some of its applications. International Journal of Biochemistry 1971;2(11):537–544.

Adzhubei, I., Jordan, D.M. and Sunyaev, S.R. Predicting functional effect of human missense mutations using PolyPhen-2. Curr. Protoc. Hum. Genet. 2013;76(1):7–20.

Adzhubei, I.A. et al., A method and server for predicting damaging missense mutations. Nat. Methods 2010;7(4):248–249.

Bendl, J. et al., PredictSNP: Robust and Accurate Consensus Classifier for Prediction of Disease-Related Mutations. PLoS Computational Biology 2014;10(1):e1003440.

Berman, H.M. et al., The Protein Data Bank. Nucleic Acids Res. 2000;28(1):235–242.

Bhattacharyya, S. and Tobacman, J.K. Arylsulfatase B regulates colonic epithelial cell migration by effects on MMP9 expression and RhoA activation. Clin Exp Metastasis 2009;26(6):535–545.

Breiman, L. Machine Learning. 2001;45(3):261–277.

Bromberg, Y. and Rost, B. SNAP: predict effect of non-synonymous polymorphisms on function. Nucleic Acids Res. 2007;35(11):3823–3835.

Bromberg, Y., Yachdav, G. and Rost, B. SNAP predicts effect of mutations on protein function. Bioinformatics 2008;24(20):2397–2398.

Calabrese, R. et al., Functional annotations improve the predictive score of human disease-related mutations in proteins. Hum. Mutat. 2009;30(8):1237–1244.

Capriotti E, et al., WS-SNPs&GO: a web server for predicting the deleterious effect of human protein variants using functional annotation. BMC genomics 2013;14(3):1–7.

Capriotti, E., Fariselli, P. and Casadio, R. I-Mutant2.0: predicting stability changes upon mutation from the protein sequence or structure. Nucleic Acids Research 2005;33(Web Server):W306–W310.

Capriotti, E., Ozturk, K. and Carter, H. Integrating molecular networks with genetic variant interpretation for precision medicine. Wiley Interdiscip. Rev. Syst. Biol. Med. 2019;11(3):e1443.

Carter, H. et al., Cancer-specific high-throughput annotation of somatic mutations: computational prediction of driver missense mutations. Cancer Res. 2009;69(16):6660–6667.

Chennen, K. et al., MISTIC: A prediction tool to reveal disease-relevant deleterious missense variants. PLoS One 2020;15(7):e0236962.

Choi, Y. and Chan, A.P. PROVEAN web server: a tool to predict the functional effect of amino acid substitutions and indels. Bioinformatics 2015;31(16):2745–2747.

Clifford, R.J. et al., Large-scale analysis of non-synonymous coding region single nucleotide polymorphisms. Bioinformatics 2004;20(7):1006–1014.

1000 Genomes Project Consortium. A global reference for human genetic variation. Nature 2015;526(7571):68–74.

The UniProt Consortium. UniProt: a worldwide hub of protein knowledge. Nucleic Acids Research 2019;47(D1):D506–D515.

Cox, M.A.A. and Cox, T.F. Multidimensional Scaling. Handbook of Data Visualization 2008:315–347.

Cristianini, N. and Shawe-Taylor, J. An Introduction to Support Vector Machines and Other Kernel-based Learning Methods. 2000.

Dasgupta, A. et al., Brief review of regression-based and machine learning methods in genetic epidemiology: the Genetic Analysis Workshop 17 experience. Genet. Epidemiol. 2011;35 Suppl 1:S5–11.

Datta, A. et al., Functional and Structural Consequences of Damaging Single Nucleotide Polymorphisms in Human Prostate Cancer Predisposition Gene RNASEL. Biomed Res Int 2015;2015:271458.

Dincer C, Kaya T, Keskin O, Gursoy A, Tuncbag N. 3D spatial organization and network-guided comparison of mutation profiles in Glioblastoma reveals similarities across patients. PLoS computational biology 2019;15(9):e1006789.

Doğan T. HPO2GO: prediction of human phenotype ontology term associations for proteins using cross ontology annotation co-occurrences. PeerJ 2018;6:e5298.

Doğan, T. et al., UniProt-DAAC: domain architecture alignment and classification, a new method for automatic functional annotation in UniProtKB. Bioinformatics 2016;32(15):2264–2271.

Doğan T, et al., Protein domain-based prediction of drug/compound–target interactions and experimental validation on LIM kinases. PLoS Computational Biology 2021;17(11):e1009171.

Doğan T, et al., CROssBAR: comprehensive resource of biomedical relations with knowledge graph representations. Nucleic Acids Research 2021;49(16):e96–e96.

Engin, H.B., Hofree, M. and Carter, H. Identifying mutation specific cancer pathways using a structurally resolved protein interaction network. Pac. Symp. Biocomput. 2015:84–95.

Farh, K.K. et al., Genetic and epigenetic fine mapping of causal autoimmune disease variants. Nature 2015;518(7539):337–343.

Fariselli P, Martelli PL, Savojardo C, Casadio R. INPS: predicting the impact of non-synonymous variations on protein stability from sequence. Bioinformatics 2015;31(17):2816–21.

Freund, Y. and Schapire, R.E. A Decision-Theoretic Generalization of On-Line Learning and an Application to Boosting. Journal of Computer and System Sciences 1997;55(1):119–139.

Friedman, J., Hastie, T. and Tibshirani, R. Additive logistic regression: a statistical view of boosting (With discussion and a rejoinder by the authors). The Annals of Statistics 2000;28(2).

Goldman, S.A. and Warmuth, M.K. Learning binary relations using weighted majority voting. Machine Learning 1995;20(3):245–271.

Goldsack, D.E. and Chalifoux, R.C. Contribution of the free energy of mixing of hydrophobic side chains to the stability of the tertiary structure of proteins. J. Theor. Biol. 1973;39(3):645–651.

González-Pérez, A. and López-Bigas, N. Improving the Assessment of the Outcome of Nonsynonymous SNVs with a Consensus Deleteriousness Score, Condel. The American Journal of Human Genetics 2011;88(4):440–449.

Grantham, R. Amino acid difference formula to help explain protein evolution. Science 1974;185(4154):862–864.

Grimm, D.G. et al., The evaluation of tools used to predict the impact of missense variants is hindered by two types of circularity. Hum. Mutat. 2015;36(5):513–523.

Guo, H.H., Choe, J. and Loeb, L.A. Protein tolerance to random amino acid change. Proc. Natl. Acad. Sci. U. S. A. 2004;101(25):9205–9210.

Halushka, M.K. et al., Patterns of single-nucleotide polymorphisms in candidate genes for blood-pressure homeostasis. Nat Genet 1999;22(3):239–247.

Hastie, T., Friedman, J. and Tibshirani, R. The Elements of Statistical Learning. Springer Series in Statistics 2001.

Heberle, H. et al., InteractiVenn: a web-based tool for the analysis of sets through Venn diagrams. BMC Bioinformatics 2015;16:169.

Hindorff, L.A. et al., Potential etiologic and functional implications of genome-wide association loci for human diseases and traits. Proc Natl Acad Sci U S A 2009;106(23):9362–9367.

Holbrook, K. et al., Functional Analysis of Semi-conserved Transit Peptide Motifs and Mechanistic Implications in Precursor Targeting and Recognition. Mol. Plant 2016;9(9):1286–1301.

Jumper, J. et al., Highly accurate protein structure prediction with AlphaFold. Nature 2021;596(7873):583–589.

Kaminker, J.S. et al., CanPredict: a computational tool for predicting cancer-associated missense mutations. Nucleic Acids Res. 2007;35(Web Server issue):W595–598.

Karageorgos, L. et al., Mutational analysis of 105 mucopolysaccharidosis type VI patients. Hum Mutat 2007;28(9):897–903.

Kawabata, T., Ota, M. and Nishikawa, K. The Protein Mutant Database. Nucleic Acids Research 1999;27(1):355–357.

Khurana, E. et al., Role of non-coding sequence variants in cancer. Nat Rev Genet 2016;17(2):93–108.

Kulmanov M, Hoehndorf R. DeepGOPlus: improved protein function prediction from sequence. Bioinformatics 2020;36(2):422–9.

Landrum, M.J. et al., ClinVar: improving access to variant interpretations and supporting evidence. Nucleic Acids Res. 2018;46(D1):D1062–D1067.

Manolio, T.A., Brooks, L.D. and Collins, F.S. A HapMap harvest of insights into the genetics of common disease. J Clin Invest 2008;118(5):1590–1605.

McCorvie, T.J. et al., Molecular basis of classic galactosemia from the structure of human galactose 1-phosphate uridylyltransferase. Hum Mol Genet 2016;25(11):2234–2244.

McGarvey, P.B. et al., UniProt genomic mapping for deciphering functional effects of missense variants. Hum. Mutat. 2019;40(6):694–705.

Meyer, M.J. et al., Interactome INSIDER: a structural interactome browser for genomic studies. Nat. Methods 2018;15(2):107–114.

Mitchell, A.L. et al., InterPro in 2019: improving coverage, classification and access to protein sequence annotations. Nucleic Acids Res. 2019;47(D1):D351–D360.

Mitternacht, S. FreeSASA: An open source C library for solvent accessible surface area calculations. F1000Res. 2016;5:189.

Momen-Roknabadi, A. et al., Impact of residue accessible surface area on the prediction of protein secondary structures. BMC Bioinformatics 2008;9:357.

Mottaz, A. et al., Easy retrieval of single amino-acid polymorphisms and phenotype information using SwissVar. Bioinformatics 2010;26(6):851–852.

Mudunuri, U. et al., bioDBnet: the biological database network. Bioinformatics 2009;25(4):555–556.

Ng, P.C. and Henikoff, S. SIFT: Predicting amino acid changes that affect protein function. Nucleic Acids Res 2003;31(13):3812–3814.

Nishi, H. et al., Cancer missense mutations alter binding properties of proteins and their interaction networks. PLoS One 2013;8(6):e66273.

Oh, S.J. Computational evaluation of interactions between olfactory receptor OR2W1 and its ligands. Genomics Inform. 2021;19(1):e9.

Pandurangan, A.P. and Blundell, T.L. Prediction of impacts of mutations on protein structure and interactions: SDM, a statistical approach, and mCSM, using machine learning. Protein Sci. 2020;29(1):247–257.

Pieper, U. et al., ModBase, a database of annotated comparative protein structure models and associated resources. Nucleic Acids Res. 2014;42(Database issue):D336–346.

Pires, D.E.V., Ascher, D.B. and Blundell, T.L. mCSM: predicting the effects of mutations in proteins using graph-based signatures. Bioinformatics 2014;30(3):335–342.

Presnyak, V. et al., Codon optimality is a major determinant of mRNA stability. Cell 2015;160(6):1111–1124.

Quan, L., Lv, Q. and Zhang, Y. STRUM: structure-based prediction of protein stability changes upon single-point mutation. Bioinformatics 2016;32(19):2936–2946.

Rentzsch, P. et al., CADD: predicting the deleteriousness of variants throughout the human genome. Nucleic Acids Res. 2019;47(D1):D886–D894.

Reva, B., Antipin, Y. and Sander, C. Determinants of protein function revealed by combinatorial entropy optimization. Genome Biol. 2007;8(11):R232.

Reva, B., Antipin, Y. and Sander, C. Predicting the functional impact of protein mutations: application to cancer genomics. Nucleic Acids Res. 2011;39(17):e118.

Rifaioglu AS, et al., Large-scale automated function prediction of protein sequences and an experimental case study validation on PTEN transcript variants. Proteins: Structure, Function, and Bioinformatics 2018;86(2):135–51.

Sasidharan Nair, P. and Vihinen, M. VariBench: a benchmark database for variations. Hum Mutat 2013;34(1):42–49.

Sauna, Z.E. and Kimchi-Sarfaty, C. Understanding the contribution of synonymous mutations to human disease. Nat Rev Genet 2011;12(10):683–691.

Savojardo C, Fariselli P, Martelli PL, Casadio R. INPS-MD: a web server to predict stability of protein variants from sequence and structure. Bioinformatics 2016;32(16):2542–4.

Schwarz, J.M. et al., MutationTaster2: mutation prediction for the deep-sequencing age. Nat. Methods 2014;11(4):361–362.

Schymkowitz, J. et al., The FoldX web server: an online force field. Nucleic Acids Res. 2005;33(Web Server issue):W382–388.

Sharma, G. et al., Reduced Arylsulfatase B activity in leukocytes from cystic fibrosis patients. Pediatr Pulmonol 2013;48(3):236–244.

Shihab, H.A. et al., Predicting the functional, molecular, and phenotypic consequences of amino acid substitutions using hidden Markov models. Hum. Mutat. 2013;34(1):57–65.

Stone, E.A. and Sidow, A. Physicochemical constraint violation by missense substitutions mediates impairment of protein function and disease severity. Genome Res. 2005;15(7):978–986.

Supek, F. et al., Synonymous mutations frequently act as driver mutations in human cancers. Cell 2014;156(6):1324–1335.

Tang, M. et al., Correlation assessment among clinical phenotypes, expression analysis and molecular modeling of 14 novel variations in the human galactose-1-phosphate uridylyltransferase gene. Hum Mutat 2012;33(7):1107–1115.

Tavtigian, S.V. et al., Classification of rare missense substitutions, using risk surfaces, with genetic- and molecular-epidemiology applications. Hum. Mutat. 2008;29(11):1342–1354.

Tavtigian, S.V. et al., Comprehensive statistical study of 452 BRCA1 missense substitutions with classification of eight recurrent substitutions as neutral. J. Med. Genet. 2006;43(4):295–305.

Thusberg, J., Olatubosun, A. and Vihinen, M. Performance of mutation pathogenicity prediction methods on missense variants. Hum Mutat 2011;32(4):358–368.

Topham, C.M., Srinivasan, N. and Blundell, T.L. Prediction of the stability of protein mutants based on structural environment-dependent amino acid substitution and propensity tables. Protein Eng. 1997;10(1):7–21.

Truebestein, L. and Leonard, T.A. Coiled-coils: The long and short of it. BioEssays 2016;38(9):903–916.

Unsal S, Atas H, Albayrak M, Turhan K, Acar AC, Doğan T. Learning functional properties of proteins with language models. Nature Machine Intelligence 2022;4(3):227–45.

van der, M. and Geoffrey Hinton, L. Visualizing Data using t-SNE. J. Mach. Learn. Res. 2008;9(86):2579–2605.

Waterhouse, A. et al., SWISS-MODEL: homology modelling of protein structures and complexes. Nucleic Acids Res. 2018;46(W1):W296–W303.

Worth, C.L., Preissner, R. and Blundell, T.L. SDM--a server for predicting effects of mutations on protein stability and malfunction. Nucleic Acids Res. 2011;39(Web Server issue):W215–222.

Wu, S.M. et al., In vitro gamma-carboxylation of a 59-residue recombinant peptide including the propeptide and the gamma-carboxyglutamic acid domain of coagulation factor IX. Effect of mutations near the propeptide cleavage site. J. Biol. Chem. 1990;265(22):13124–13129.

Yang, Y. et al., Structure-based prediction of the effects of a missense variant on protein stability. Amino Acids 2013;44(3):847–855.

Yue, P., Li, Z. and Moult, J. Loss of protein structure stability as a major causative factor in monogenic disease. J. Mol. Biol. 2005;353(2):459–473.

Yue, P. and Moult, J. Identification and Analysis of Deleterious Human SNPs. Journal of Molecular Biology 2006;356(5):1263–1274.

Zwart, M.P. et al., Unraveling the causes of adaptive benefits of synonymous mutations in TEM-1 beta-lactamase. Heredity (Edinb) 2018;121(5):406–421.

