## Supplementary Information for "ASCARIS: Positional Feature Annotation and Protein Structure-Based Representation of Single Amino Acid Variations"

#### S1. A literature review on variant effect predictors

Computational variant effect predictors can be classified according to their input features. One of the most widely used information sources for identifying the characteristics of a variation is evolutionary conservation derived from the biomolecular sequence. The methods that utilize conservation are based on the alignment of the sequences. For example, SIFT (Ng and Henikoff, 2003) identifies conservation among similar sequences to decide whether a given substitution will be tolerated in that particular position. PROVEAN (Choi and Chan, 2015), MAPP (Stone and Sidow, 2005), Align-GVGD (Tavtigian et al., 2006; Tavtigian et al., 2008), and MutationAssessor (Reva et al., 2007) are other examples to tools with evolutionary conservation-based models. MutationTaster (Schwarz et al., 2014) and CADD (Rentzsch et al., 2019) utilize sequence-derived information such as splice sites, Kozak consensus sequences, etc. to determine the degree of conservation of a given nucleotide.

The 3-D structure of proteins is another source of information utilized by variant effect predictors. These methods mostly aim to identify the changes in structural characteristics of proteins upon mutation, such as stability or contact energies (Yang et al., 2013; Yue et al., 2005). The method, DUET (Pandurangan and Blundell, 2020), combines two other structure-based approaches to predict the effect of mutations on protein stability. There are also a few methods that try to predict changes in stability. Since a protein's stability is critical for its functionality, these predictions ultimately guide the effort to predict the functional effects of variations. Energy calculations evaluating (Quan et al., 2016; Schymkowitz et al., 2005) hydrophobic area, overpacking, backbone strain, loss/gain of electrostatic interactions (Capriotti et al., 2005; Yue et al., 2005) are some of the approaches used to predict stability changes upon single amino acid variations.

An alternative approach is to combine structure and sequence-related features with the aim of increasing the predictive performance. A well-known tool, PolyPhen-2, incorporates structure-derived information, such as accessible surface area values and hydrophobic propensities, together with sequence conservation through alignment, to distinguish between the wild type and varied amino acids (Adzhubei et al., 2013; Adzhubei et al., 2010). SNAP (Bromberg et al., 2008) and SNPs&GO (Calabrese et al., 2009) are two other predictors that employed both sequence and structure-derived features. From sequence alignment, SNAP calculates the likelihood of observing certain mutations in a position and the degree to which a residue is conserved in a family of related proteins. It also adds features like hydrophobicity, charge, size, and the presence of a buried charge. SNPs&GO utilizes conservation profiles of the amino acids as well as structural features such as the count of residues in a radius of 6 Å around the Cα atom and relative accessible surface area of the mutated residue in 3D. MISTIC (Chennen et al., 2020) evaluates evolutionary conservation, biochemical features and multi-allele frequencies on genomic variations. A few methods integrate scores from other predictors within a meta prediction approach. A cancer-specific tool CanPredict (Kaminker et al., 2007) combines scores from SIFT, LogR.E-value (Clifford et al., 2004) and GOSS metrics (Clifford et al., 2004; Kaminker et al., 2007) to distinguish cancer-associated mutations.

Variant effect prediction methods can also be grouped in terms of the algorithmic approach they use. SIFT incorporates a statistical approach by calculating the probability of amino acid substitutions being tolerated at different positions in the sequence. MAPP uses a statistical framework in which it estimates the physicochemical constraints on each position in the alignment column that corresponds to the variant and computes a single score measuring the violation of these constraints. MutationTaster, which employs a Bayesian classifier, can be counted as another example of probabilistic classifiers. PROVEAN, on the other hand, introduces an alignment-based score to measure the change in sequence similarities caused by a variation, to predict its damaging effect. Another method Align-GVGD calculates two scores; the Grantham Variation (GV) for positions in protein multiple sequence alignments and the Grantham Deviation (GD) for missense substitutions at those positions. Both GD and GV are extensions of the original Grantham difference (Grantham, 1974).

Machine learning (ML) techniques are utilized in variant effect prediction due to their ability to identify hidden patterns in complex and high-dimensional data. SNPs3D (Carter et al., 2009; Yue and Moult, 2006) and SNPs&GO (Calabrese et al., 2009) use support vector machines (SVM) to partition the sample space as deleterious or neutral (Pandurangan and Blundell, 2020; Pires et al., 2014; Rentzsch et al., 2019; Topham et al., 1997; Worth et al., 2011). A widely used tool, PolyPhen-2, employs a Naive Bayes classifier to calculate the posterior probability of mutations. MISTIC does its classification by integrating a soft voting system (Goldman and Warmuth, 1995). CHASM (Carter et al., 2009) and Can-Predict (Kaminker et al., 2007) are methods that employ random forest (RF) to predict cancer-associated mutations. Finally, neural networks are utilized in variant effect predictors (Bromberg and Rost, 2007; Bromberg et al., 2008; Calabrese et al., 2009).

### S2. Significance analysis of domains

We applied a statistical test to retrieve domains that are statistically significant considering SAVs (i.e., variation data points that are located in the domain region in the protein sequence) being either predominantly neutral or deleterious. For each domain, the number of occurrences for deleterious and neutral SAVs is recorded. We employed Fisher's exact test for the statistical analysis, where the contingency table is composed of the numbers of SAVs within the domain region as neutral and deleterious, and the numbers of SAVs that are located out of that domain's region as neutral and deleterious. We identified significant domains at the 99% confidence interval. **Table S2** shows these 314 significant domains and their statistics along with the significance values. Following the filtering of SAVs whose proteins do not have available structure models, 307 of these domains are found in our dataset, and the InterPro identifiers of these domains are included in the "significant domains" (categorical) variable of our SAV feature vectors. Domains that are not found to be significant are merged and included in the "significant domains" variable under one generic/dummy label, "DomainX", as the 308th category.

### S3. Distance calculation for the dimensionality reduction analysis

It was required to calculate pairwise similarities between SAV data points for the t-SNE embeddings and the MDS analysis. This is not a straightforward process since our feature vectors are composed of both real valued and categorical variables. To address this issue, we came up with a simple heuristic to merge these variables under 4-dimensional distance vectors with directly comparable dimensions (each have values between 0 and 1), and took the average of these 4 distance values to calculate the mean pairwise distance between two SAVs. For each SAV pair:

a) Physicochemical distance:

- The actual difference in terms of Grantham scores between two SAV data points are normalized using the ‘min-max normalization’, and these normalized values are recorded as the distance value between these two data points.

b) Domain distance:

- If both SAV data points are missing domain annotation, the distance value is left blank.
- If one data point is missing a domain annotation and the other one has a domain annotation, the value is recorded as 1 to indicate maximum distance.
- If both data points are annotated with different domains, the value is again recorded as 1; and if both are annotated with the same domain the distance value is recorded as 0.

c) Variation location distance:

- If both SAV data points are annotated to different regions (i.e., core, interface, surface), the distance value is recorded as 1, otherwise the distance value is recorded as 0.
- If one or both of them are missing location annotation; the value is left blank.

d) Positional sequence annotation distance (for each positional annotation type):

- If both SAV data points correspond to the same type of positional annotation, a count that indicates “commonality” is increased by 1.
- If one SAV is corresponding and the other SAV is not, a count that indicates “difference” is increased by 1.
- If both are non-corresponding, there is no count.
- This procedure is repeated for 30 different positional annotation types, and finally, the distance score is calculated by: 1-(commonality/(commonality+difference)).

As the final pairwise distance score, the mean of these 4 distance values is recorded for each SAV data point pair.

### Supplementary References

Adzhubei, I., Jordan, D.M. and Sunyaev, S.R. Predicting functional effect of human missense mutations using PolyPhen-2. Curr. Protoc. Hum. Genet. 2013;Chapter 7:Unit7.20.

Adzhubei, I.A. et al., A method and server for predicting damaging missense mutations. Nat. Methods 2010;7(4):248-249.

Bromberg, Y. and Rost, B. SNAP: predict effect of non-synonymous polymorphisms on function. Nucleic Acids Res. 2007;35(11):3823-3835.

Bromberg, Y., Yachdav, G. and Rost, B. SNAP predicts effect of mutations on protein function. Bioinformatics 2008;24(20):2397-2398.

Calabrese, R. et al., Functional annotations improve the predictive score of human disease-related mutations in proteins. Hum. Mutat. 2009;30(8):1237-1244.

Carter, H. et al., Cancer-specific high-throughput annotation of somatic mutations: computational prediction of driver missense mutations. Cancer Res. 2009;69(16):6660-6667.

Chennen, K. et al., MISTIC: A prediction tool to reveal disease-relevant deleterious missense variants. PLoS One 2020;15(7):e0236962.

Choi, Y. and Chan, A.P. PROVEAN web server: a tool to predict the functional effect of amino acid substitutions and indels. Bioinformatics 2015;31(16):2745-2747.

Clifford, R.J. et al., Large-scale analysis of non-synonymous coding region single nucleotide polymorphisms. Bioinformatics 2004;20(7):1006-1014.

Goldman, S.A. and Warmuth, M.K. Learning binary relations using weighted majority voting. Machine Learning 1995;20(3):245-271.

Grantham, R. Amino acid difference formula to help explain protein evolution. Science 1974;185(4154):862-864.

Kaminker, J.S. et al., CanPredict: a computational tool for predicting cancer-associated missense mutations. Nucleic Acids Res. 2007;35(Web Server issue):W595-598.

Ng, P.C. and Henikoff, S. SIFT: Predicting amino acid changes that affect protein function. Nucleic Acids Res 2003;31(13):3812-3814.

Pandurangan, A.P. and Blundell, T.L. Prediction of impacts of mutations on protein structure and interactions: SDM, a statistical approach, and mCSM, using machine learning. Protein Sci. 2020;29(1):247-257.

Pires, D.E.V., Ascher, D.B. and Blundell, T.L. mCSM: predicting the effects of mutations in proteins using graph-based signatures. Bioinformatics 2014;30(3):335-342.

Quan, L., Lv, Q. and Zhang, Y. STRUM: structure-based prediction of protein stability changes upon single-point mutation. Bioinformatics 2016;32(19):2936-2946.

Rentzsch, P. et al., CADD: predicting the deleteriousness of variants throughout the human genome. Nucleic Acids Res. 2019;47(D1):D886-D894.

Reva, B., Antipin, Y. and Sander, C. Determinants of protein function revealed by combinatorial entropy optimization. Genome Biol. 2007;8(11):R232.

Schwarz, J.M. et al., MutationTaster2: mutation prediction for the deep-sequencing age. Nat. Methods 2014;11(4):361-362.

Schymkowitz, J. et al., The FoldX web server: an online force field. Nucleic Acids Res. 2005;33(Web Server issue):W382-388.

Stone, E.A. and Sidow, A. Physicochemical constraint violation by missense substitutions mediates impairment of protein function and disease severity. Genome Res. 2005;15(7):978-986.

Tavtigian, S.V. et al., Classification of rare missense substitutions, using risk surfaces, with genetic- and molecular-epidemiology applications. Hum. Mutat. 2008;29(11):1342-1354.

Tavtigian, S.V. et al., Comprehensive statistical study of 452 BRCA1 missense substitutions with classification of eight recurrent substitutions as neutral. J. Med. Genet. 2006;43(4):295-305.

Topham, C.M., Srinivasan, N. and Blundell, T.L. Prediction of the stability of protein mutants based on structural environment-dependent amino acid substitution and propensity tables. Protein Eng. 1997;10(1):7-21.

Worth, C.L., Preissner, R. and Blundell, T.L. SDM--a server for predicting effects of mutations on protein stability and malfunction. Nucleic Acids Res. 2011;39(Web Server issue):W215-222.

Yang, Y. et al., Structure-based prediction of the effects of a missense variant on protein stability. Amino Acids 2013;44(3):847-855.

Yue, P., Li, Z. and Moult, J. Loss of protein structure stability as a major causative factor in monogenic disease. J. Mol. Biol. 2005;353(2):459-473.

Yue, P. and Moult, J. Identification and Analysis of Deleterious Human SNPs. Journal of Molecular Biology 2006;356(5):1263-1274.

### Supplementary Figures

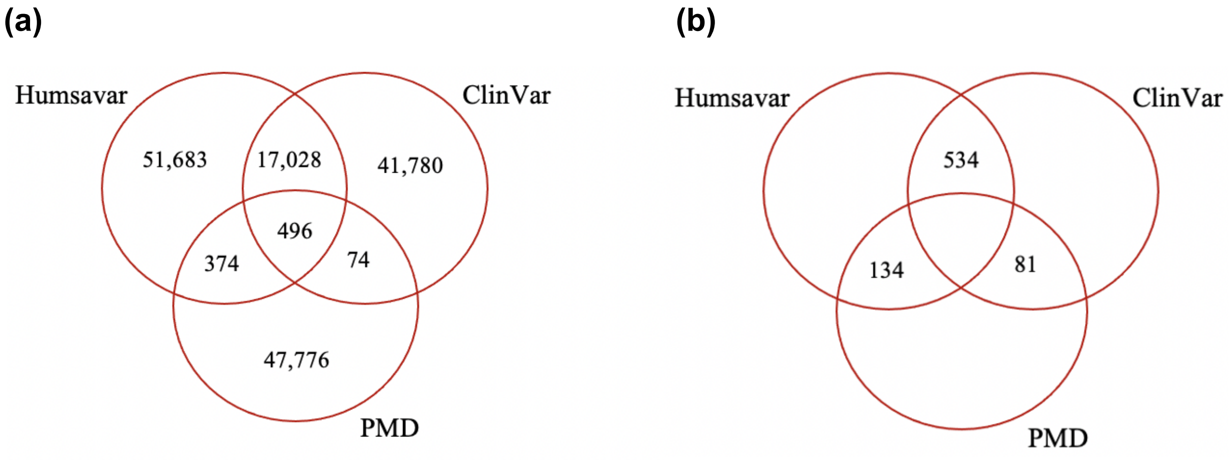


**Figure S1. (a)** Number of common (i.e., the SAV of interest is found in multiple resources with a consistent/agreeable label, either as neutral or deleterious on both resources), and **(b)** conflicting (i.e., the SAV of interest is labelled as neutral in one resource, and as deleterious in the other) data points for each database.

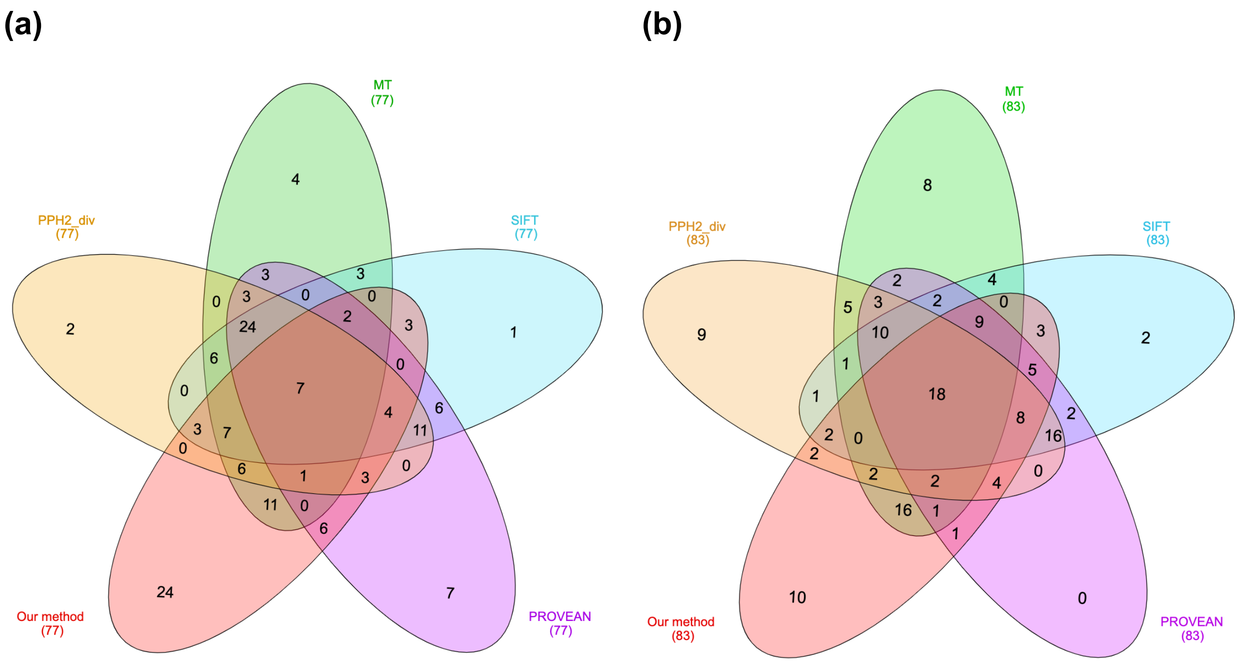


**Figure S2.** Intersecting predictions between our method and the state-of-the-art methods on **(a)** neutral, and **(b)** deleterious SAVs in benchmark 1 (the MutationTaster dataset) shown via Venn diagrams.

**(a)**

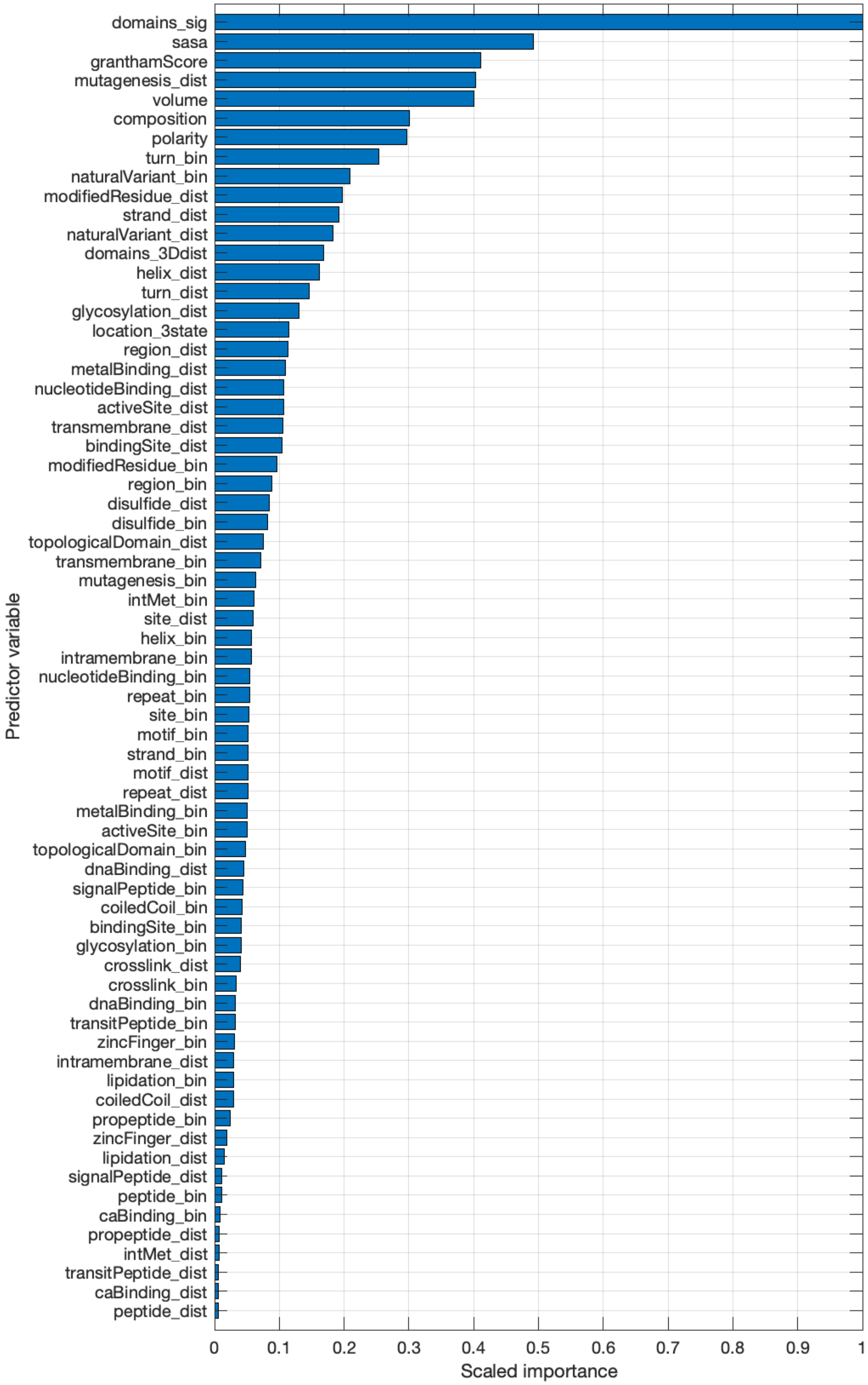
domains_sig
sasa
granthamScore
mutagenesis_dist
volume
composition
polarity
turn_bin
naturalVariant_bin
modifiedResidue_dist
strand_dist
naturalVariant_dist
domains_3Ddist
helix_dist
turn_dist
glycosylation_dist
location_3state
region_dist
metalBinding_dist
nucleotideBinding_dist
activeSite_dist
transmembrane_dist
bindingSite_dist
modifiedResidue_bin
region_bin
disulfide_dist
disulfide_bin
topologicalDomain_dist
transmembrane_bin
mutagenesis_bin
intMet_bin
site_dist
helix_bin
intramembrane_bin
nucleotideBinding_bin
repeat_bin
site_bin
motif_bin
strand_bin
motif_dist
repeat_dist
metalBinding_bin
activeSite_bin
topologicalDomain_bin
dnaBinding_dist
signalPeptide_bin
coiledCoil_bin
bindingSite_bin
glycosylation_bin
crosslink_dist
crosslink_bin
dnaBinding_bin
transitPeptide_bin
zincFinger_bin
intramembrane_dist
lipidation_bin
coiledCoil_dist
propeptide_bin
zincFinger_dist
lipidation_dist
signalPeptide_dist
peptide_bin
caBinding_bin
propeptide_dist
intMet_dist
transitPeptide_dist
caBinding_dist
peptide_dist
Predictor variable
0
0.1
0.2
0.3
0.4
0.5
0.6
0.7
0.8
0.9
1
Scaled importance

(b)

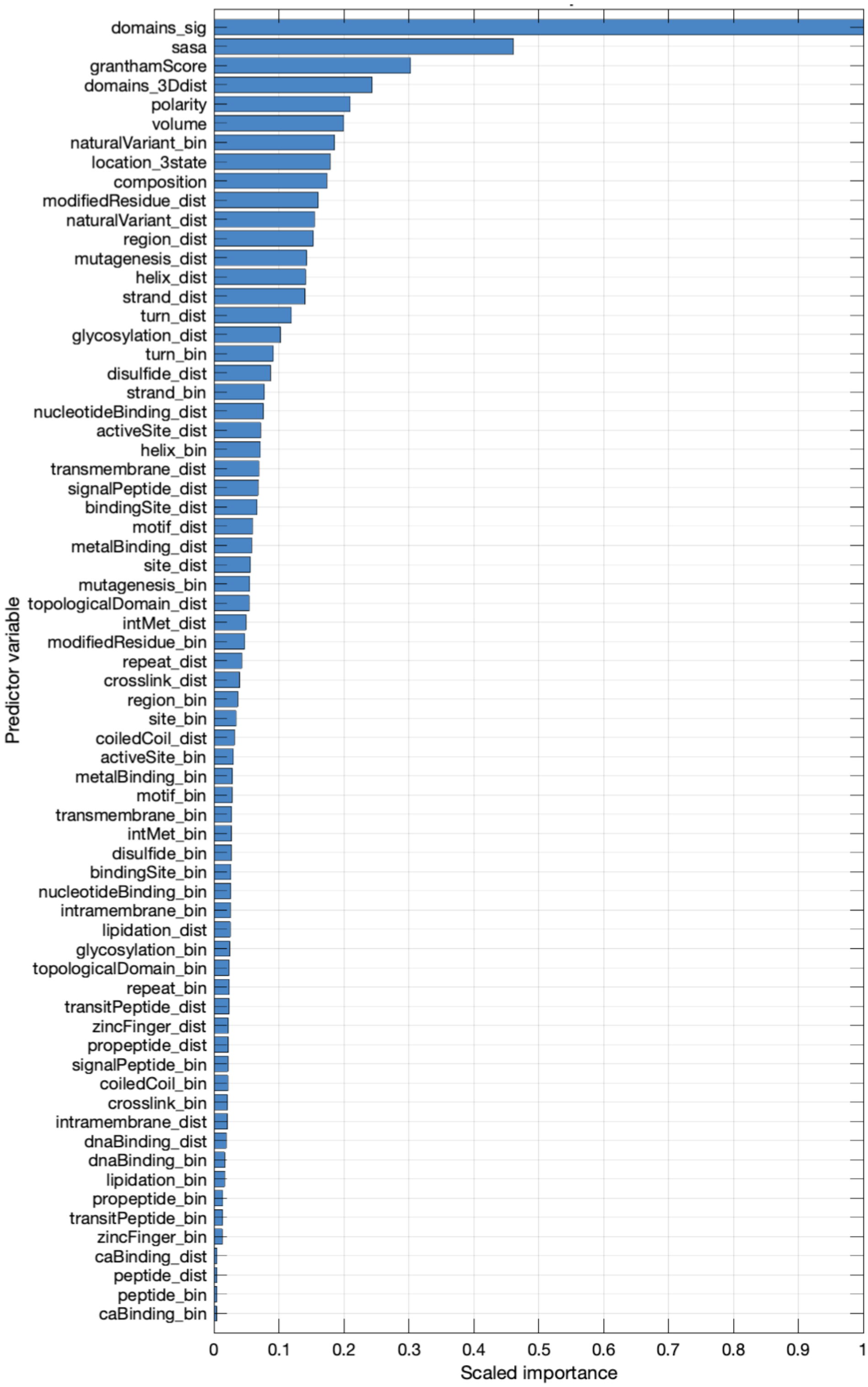


**Figure S3.** Scaled feature importance measurements of the random forest variant effect classification model that utilize all types of features (i.e., 68-dimensional vectors) with optimal hyper-parameters for; **(a)** the main PDB and modeling-based analysis, and **(b)** AlphaFold2 tracks.

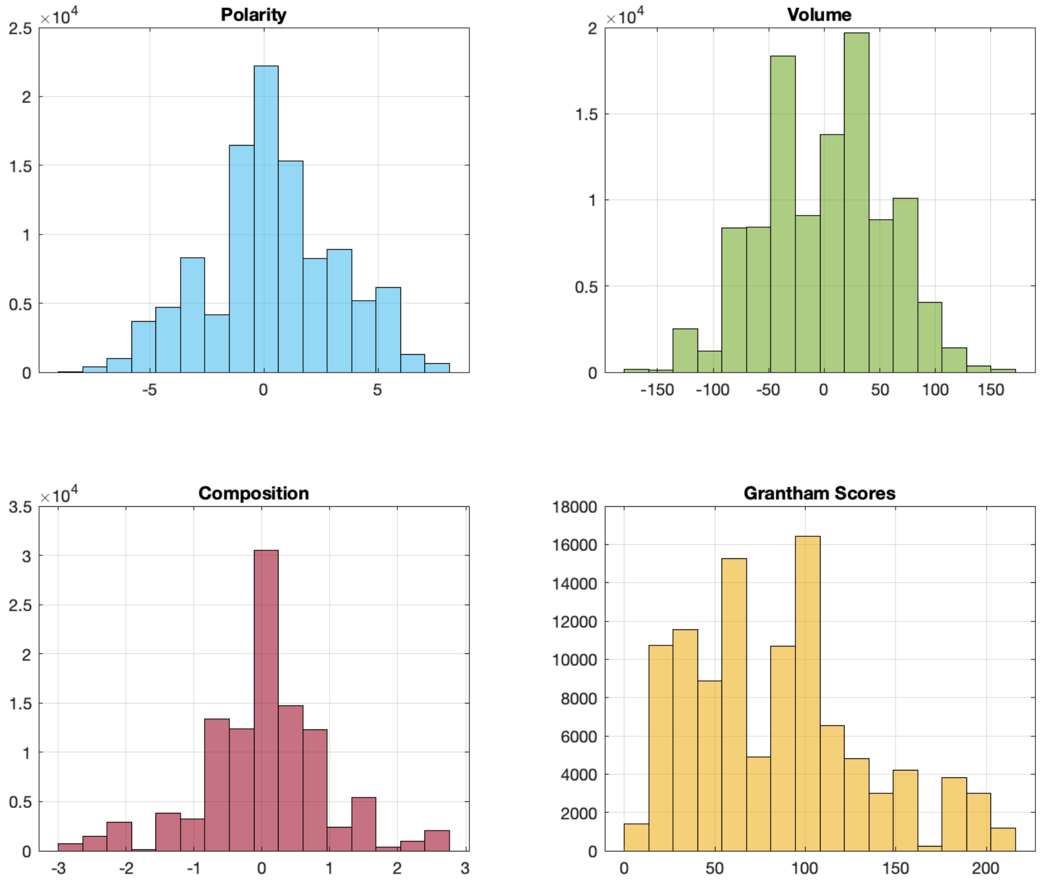


**Figure S4.** The distributions of residue-level physicochemical value changes in single amino acid variations in our dataset.

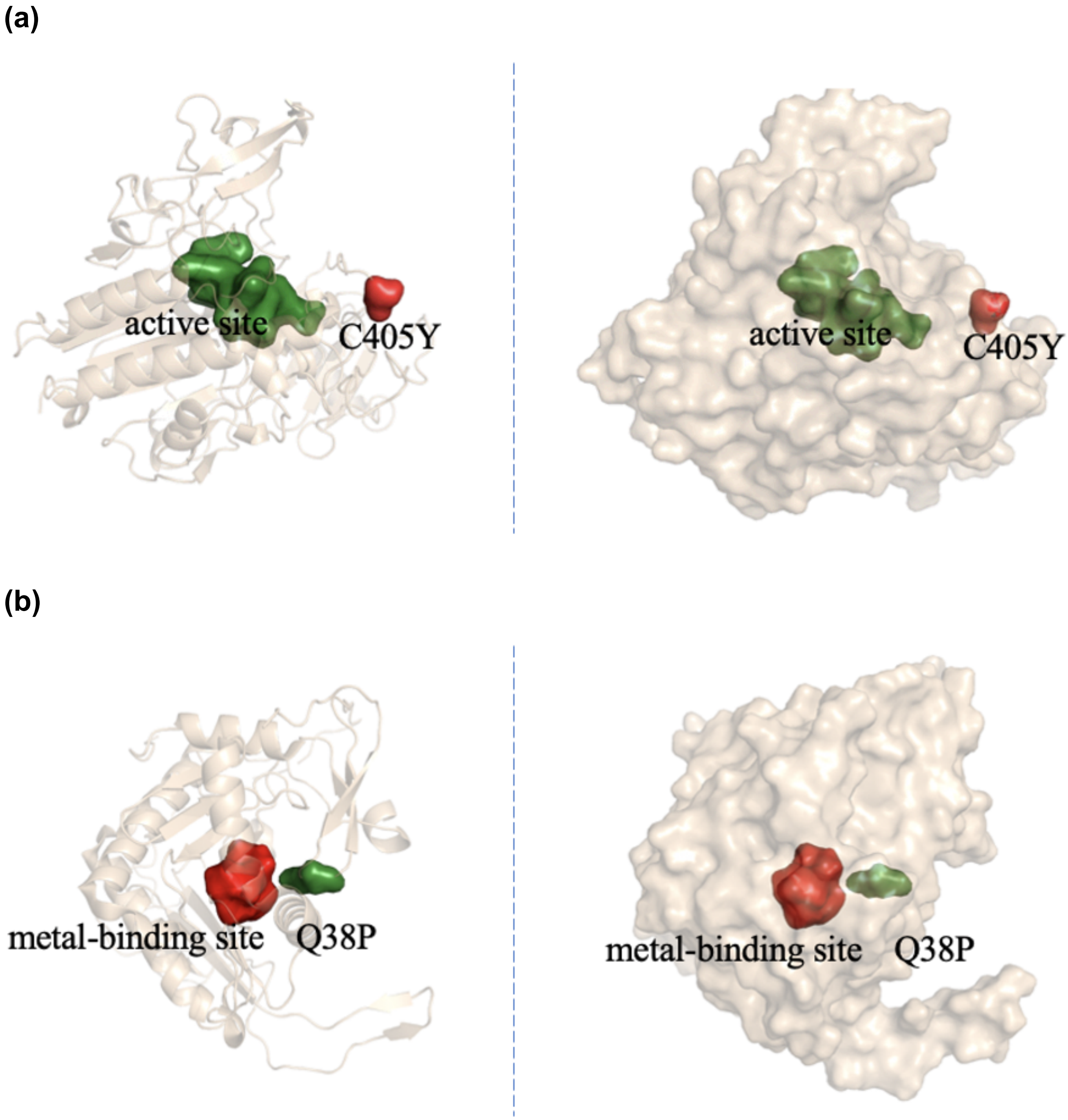


**Figure S5.** Structural representations (cartoon and surface from left to right) of; **(a)** ASB, and **(b)** GALT proteins, on which the mutations of interest (red) and reported active sites (green) are highlighted.

### Supplementary Tables

**Table S1.** List of variables in the output delimited file of the method, which contain the 74-dimensional SAV feature set (containing the metadata of variations in 5 of these dimensions).

| Column title/name | Order of the variable | Description | Data type |
|---|---|---|---|
| prot_uniprotAcc | 1 | Information about the mutation: UniProt accession of the protein, wild type residue, mutated residue, and the position of the variation on the sequence (metadata, not part of the feature vector) | - |
| wt_residue | 2 | | |
| mut_residue | 3 | | |
| position | 4 | | |
| meta_merged | 5 | Merged meta data: prot_uniprotAcc+wt_residue+ mut_residue+position (metadata, not part of the SAV feature vector) | - <br> - |
| composition | 6 | The change in physicochemical feature values upon the occurrence of variation | Real values |
| polarity | 7 | | |
| volume | 8 | | |
| granthamScore | 9 | | |
| domains_all | 10 | Domains (all) that the mutation corresponds to (not suggested to be used in the SAV feature vector) | Categorical (2204 domains, given with their InterPro identifiers) |
| domains_sig | 11 | Domains (only statistically significant ones) that the mutation corresponds to (please use this variable in the SAV feature vector instead of dimension #10) | Categorical (307 domains, given with their InterPro identifiers, an additional category called “domainX” is used to represent the rest of the domains and no domain cases) |
| domains_3Ddist | 12 | Spatial distance to nearest domain for the cases where the mutation does not fall inside a domain region | Real values (if the residue of SAV is already in the domain region, the corresponding dimension is imputed with 0) |
| sasa | 13 | Solvent-accessible surface area (SASA) values | Real values |
| location_3state | 14 | Location of the mutation on the structure (calculated from SASA values) | Categorical (core, interface, surface or unknown) |
| disulfide_bin | 15 | Positional sequence annotations, binary correspondence-based (30 different types of annotations, each one on a different dimension) | Categorical (0: not annotated to the protein, 1: annotation is presented on the protein but the mutated position does not correspond to the annotation, 2: mutation corresponds to the annotation on the same position) |
| intMet_bin | 16 | | |
| intramembrane_bin | 17 | | |
| naturalVariant_bin | 18 | | |
| dnaBinding_bin | 19 | | |
| activeSite_bin | 20 | | |
| nucleotideBinding_bin | 21 | | |
| lipidation_bin | 22 | | |
| site_bin | 23 | | |
| transmembrane_bin | 24 | | |

| | | | |
|---|---|---|---|
| crosslink_bin | 25 | | |
| mutagenesis_bin | 26 | | |
| strand_bin | 27 | | |
| helix_bin | 28 | | |
| turn_bin | 29 | | |
| metalBinding_bin | 30 | | |
| repeat_bin | 31 | | |
| caBinding_bin | 32 | | |
| topologicalDomain_bin | 33 | | |
| bindingSite_bin | 34 | | |
| region_bin | 35 | | |
| signalPeptide_bin | 36 | | |
| modifiedResidue_bin | 37 | | |
| zincFinger_bin | 38 | | |
| motif_bin | 39 | | |
| coiledCoil_bin | 40 | | |
| peptide_bin | 41 | | |
| transitPeptide_bin | 42 | | |
| glycosylation_bin | 43 | | |
| propeptide_bin | 44 | | |
| disulfide_dist | 45 | Positional sequence annotations, distance-based (the spatial distance between the annotated residue and the mutated residue, in the protein structure, for 30 different types of annotations, each one on a different dimension) | Real values (if the annotation does not exist on the respective protein, the corresponding dimension is imputed with the distance value at the mid-point of neutral and deleterious mutations' distance distributions) |
| intMet_dist | 46 | | |
| intramembrane_dist | 47 | | |
| naturalVariant_dist | 48 | | |
| dnaBinding_dist | 49 | | |
| activeSite_dist | 50 | | |
| nucleotideBinding_dist | 51 | | |
| lipidation_dist | 52 | | |
| site_dist | 53 | | |
| transmembrane_dist | 54 | | |
| crosslink_dist | 55 | | |
| mutagenesis_dist | 56 | | |
| strand_dist | 57 | | |
| helix_dist | 58 | | |
| turn_dist | 59 | | |
| metalBinding_dist | 60 | | |
| repeat_dist | 61 | | |
| caBinding_dist | 62 | | |
| topologicalDomain_dist | 63 | | |
| bindingSite_dist | 64 | | |
| region_dist | 65 | | |
| signalPeptide_dist | 66 | | |
| modifiedResidue_dist | 67 | | |
| zincFinger_dist | 68 | | |
| motif_dist | 69 | | |
| coiledCoil_dist | 70 | | |
| peptide_dist | 71 | | |
| transitPeptide_dist | 72 | | |
| glycosylation_dist | 73 | | |
| propeptide_dist | 74 | | |

**Table S2.** The list of significant domains in terms of discerning deleterious variations from neutral ones.

| Domain InterPro ID | Domain name | Total # of occurrence | Deleterious cases | Neutral cases | p-value of the difference |
|---|---|---|---|---|---|
| IPR011162 | MHC classes I/II-like antigen recognition protein | 1161 | 92 | 1069 | 3.90E-299 |
| IPR005821 | Ion transport domain | 2026 | 1829 | 197 | 4.56E-226 |
| IPR017452 | GPCR | 1975 | 557 | 1418 | 1.43E-170 |
| IPR013783 | Immunoglobulin-like fold | 3258 | 1271 | 1987 | 7.24E-119 |
| IPR009050 | Globin-like superfamily | 588 | 82 | 506 | 6.79E-113 |
| IPR017853 | Glycoside hydrolase superfamily | 882 | 761 | 121 | 4.51E-72 |
| IPR017850 | Alkaline-phosphatase-like | 834 | 722 | 112 | 7.96E-70 |
| IPR008972 | Cupredoxin | 477 | 438 | 39 | 9.09E-60 |
| IPR013087 | Zinc finger C2H2-type | 575 | 150 | 425 | 7.61E-57 |
| IPR019774 | Aromatic amino acid hydroxylase | 270 | 262 | 8 | 1.13E-49 |
| IPR013092 | Connexin | 450 | 404 | 46 | 1.39E-48 |
| IPR015919 | Cadherin-like superfamily | 596 | 176 | 420 | 3.13E-47 |
| IPR008967 | p53-like transcription factor | 482 | 426 | 56 | 7.28E-47 |
| IPR000536 | Nuclear hormone receptor | 523 | 442 | 81 | 2.03E-37 |
| IPR011146 | HIT-like domain | 200 | 192 | 8 | 2.50E-34 |
| IPR001909 | Krueppel-associated box | 87 | 0 | 87 | 3.62E-34 |
| IPR000742 | EGF-like domain | 1525 | 1117 | 408 | 5.72E-33 |
| IPR013088 | Zinc finger | 191 | 183 | 8 | 2.09E-32 |
| IPR029021 | Protein-tyrosine phosphatase-like | 468 | 392 | 76 | 1.37E-31 |
| IPR027417 | P-loop containing nucleoside triphosphate hydrolase | 3339 | 2273 | 1066 | 2.14E-30 |
| IPR022772 | von Hippel-Lindau disease tumour suppressor | 150 | 146 | 4 | 1.65E-28 |
| IPR016024 | Armadillo-type fold | 806 | 318 | 488 | 2.37E-28 |
| IPR008948 | L-Aspartase-like | 158 | 152 | 6 | 7.28E-28 |
| IPR013320 | Concanavalin A-like lectin/glucanase domain superfamily | 570 | 207 | 363 | 4.28E-27 |
| IPR011991 | ArsR-like helix-turn-helix domain | 570 | 447 | 123 | 2.03E-23 |
| IPR009057 | Homeobox-like domain superfamily | 379 | 312 | 67 | 7.50E-23 |
| IPR020858 | Serum albumin-like | 91 | 9 | 82 | 2.15E-22 |
| IPR006132 | Aspartate/ornithine carbamoyltransferase | 108 | 106 | 2 | 3.49E-22 |
| IPR029057 | Phosphoribosyltransferase-like | 113 | 110 | 3 | 6.95E-22 |
| IPR014710 | RmlC-like jelly roll fold | 204 | 181 | 23 | 5.26E-21 |
| IPR016176 | Cobalamin (vitamin B12)-dependent enzyme | 108 | 105 | 3 | 1.22E-20 |
| IPR003644 | Na-Ca exchanger/integrin-beta4 | 70 | 5 | 65 | 9.40E-20 |
| IPR006131 | Aspartate/ornithine carbamoyltransferase | 101 | 98 | 3 | 3.47E-19 |
| IPR009100 | Acyl-CoA dehydrogenase/oxidase | 159 | 144 | 15 | 4.58E-19 |
| IPR017943 | Bactericidal permeability-increasing protein | 129 | 27 | 102 | 3.36E-18 |
| IPR029063 | S-adenosyl-L-methionine-dependent methyltransferase | 251 | 80 | 171 | 1.13E-17 |
| IPR023214 | HAD superfamily | 355 | 284 | 71 | 1.59E-17 |
| IPR001478 | PDZ domain | 96 | 16 | 80 | 3.07E-17 |
| IPR008280 | Tubulin/FtsZ | 137 | 124 | 13 | 1.60E-16 |
| IPR008922 | Uncharacterised domain | 121 | 111 | 10 | 7.73E-16 |
| IPR004273 | Dynein heavy chain region D6 P-loop domain | 86 | 14 | 72 | 1.11E-15 |
| IPR022417 | Porphobilinogen deaminase | 61 | 61 | 0 | 9.31E-15 |

| | | | | | |
|---|---|---|---|---|---|
| IPR015943 | WD40/YVTN repeat-like-containing domain superfamily | 605 | 261 | 344 | 1.41E-14 |
| IPR015813 | Pyruvate/Phosphoenolpyruvate kinase-like domain superfamily | 77 | 74 | 3 | 4.74E-14 |
| IPR029061 | Thiamin diphosphate-binding fold | 164 | 141 | 23 | 5.09E-14 |
| IPR011990 | Tetratricopeptide-like helical domain superfamily | 277 | 101 | 176 | 9.17E-14 |
| IPR013821 | Potassium channel | 74 | 71 | 3 | 2.40E-13 |
| IPR004841 | Amino acid permease/ SLC12A domain | 89 | 83 | 6 | 3.25E-13 |
| IPR014743 | Chloride channel | 169 | 143 | 26 | 4.87E-13 |
| IPR011042 | Six-bladed beta-propeller | 429 | 323 | 106 | 5.99E-13 |
| IPR002181 | Fibrinogen | 76 | 14 | 62 | 7.75E-13 |
| IPR009075 | Acyl-CoA dehydrogenase/oxidase C-terminal | 165 | 139 | 26 | 1.76E-12 |
| IPR008979 | Galactose-binding-like domain superfamily | 286 | 224 | 62 | 3.40E-12 |
| IPR001926 | Pyridoxal-phosphate dependent enzyme | 97 | 88 | 9 | 4.97E-12 |
| IPR022673 | Hexokinase | 137 | 118 | 19 | 5.36E-12 |
| IPR015424 | Pyridoxal phosphate-dependent transferase | 358 | 272 | 86 | 7.28E-12 |
| IPR024074 | Argininosuccinate synthetase | 55 | 54 | 1 | 9.28E-12 |
| IPR022675 | Glucose-6-phosphate dehydrogenase | 76 | 71 | 5 | 1.32E-11 |
| IPR022636 | S-adenosylmethionine synthetase superfamily | 54 | 53 | 1 | 1.64E-11 |
| IPR001811 | Chemokine interleukin-8-like domain | 28 | 0 | 28 | 1.76E-11 |
| IPR017878 | TB domain | 154 | 129 | 25 | 3.29E-11 |
| IPR001007 | VWFC domain | 60 | 10 | 50 | 3.80E-11 |
| IPR028992 | Hedgehog/Intein (Hint) domain | 46 | 46 | 0 | 4.09E-11 |
| IPR008983 | Tumour necrosis factor-like domain superfamily | 345 | 261 | 84 | 4.43E-11 |
| IPR023298 | P-type ATPase | 78 | 72 | 6 | 6.39E-11 |
| IPR000859 | CUB domain | 106 | 29 | 77 | 7.26E-11 |
| IPR018884 | Glutamate [NMDA] receptor | 50 | 7 | 43 | 8.44E-11 |
| IPR001846 | von Willebrand factor | 153 | 50 | 103 | 9.85E-11 |
| IPR031688 | Voltage-gated calcium channel subunit alpha | 26 | 0 | 26 | 1.03E-10 |
| IPR022672 | Hexokinase | 105 | 92 | 13 | 1.16E-10 |
| IPR001424 | Superoxide dismutase | 127 | 108 | 19 | 1.74E-10 |
| IPR009071 | High mobility group box domain | 92 | 82 | 10 | 1.83E-10 |
| IPR032695 | Integrin domain superfamily | 77 | 18 | 59 | 3.53E-10 |
| IPR008930 | Terpenoid cyclases/protein prenyltransferase alpha-alpha toroid | 70 | 15 | 55 | 3.68E-10 |
| IPR024041 | Ammonium transporter AmtB-like domain | 51 | 8 | 43 | 3.69E-10 |
| IPR013816 | ATP-grasp fold | 85 | 76 | 9 | 6.36E-10 |
| IPR017448 | SRCR-like domain | 53 | 9 | 44 | 9.20E-10 |
| IPR003533 | Doublecortin domain | 118 | 100 | 18 | 1.45E-09 |
| IPR015425 | Formin | 23 | 0 | 23 | 1.47E-09 |
| IPR020683 | Ankyrin repeat-containing domain | 344 | 146 | 198 | 1.57E-09 |
| IPR027387 | Cytochrome b/b6-like domain superfamily | 34 | 3 | 31 | 1.60E-09 |
| IPR000980 | SH2 domain | 218 | 170 | 48 | 2.04E-09 |
| IPR009045 | Hedgehog signalling/DD-peptidase zinc-binding domain superfamily | 50 | 48 | 2 | 2.35E-09 |
| IPR000034 | Laminin IV | 22 | 0 | 22 | 3.55E-09 |
| IPR006207 | Cystine knot | 68 | 62 | 6 | 3.58E-09 |
| IPR002035 | von Willebrand factor | 429 | 191 | 238 | 3.80E-09 |
| IPR010982 | Lambda repressor-like | 63 | 58 | 5 | 4.35E-09 |
| IPR016185 | Pre-ATP-grasp domain superfamily | 64 | 59 | 5 | 4.38E-09 |

| | | | | | |
|---|---|---|---|---|---|
| IPR001452 | SH3 domain | 134 | 45 | 89 | 5.64E-09 |
| IPR013780 | Glycosyl hydrolase | 76 | 68 | 8 | 5.95E-09 |
| IPR028889 | Ubiquitin specific protease domain | 41 | 6 | 35 | 7.80E-09 |
| IPR016137 | RGS domain | 32 | 3 | 29 | 7.83E-09 |
| IPR008250 | P-type ATPase | 162 | 130 | 32 | 7.95E-09 |
| IPR023416 | Transthyretin/hydroxyisourate hydrolase domain | 103 | 88 | 15 | 8.01E-09 |
| IPR002912 | ACT domain | 33 | 33 | 0 | 2.65E-08 |
| IPR003008 | Tubulin/FtsZ | 95 | 81 | 14 | 2.78E-08 |
| IPR013785 | Aldolase-type TIM barrel | 186 | 145 | 41 | 3.58E-08 |
| IPR032675 | Leucine-rich repeat domain superfamily | 653 | 314 | 339 | 4.97E-08 |
| IPR000585 | Hemopexin-like domain | 44 | 8 | 36 | 7.62E-08 |
| IPR029332 | PEHE domain | 18 | 0 | 18 | 1.22E-07 |
| IPR029041 | FAD-linked oxidoreductase-like | 51 | 47 | 4 | 1.80E-07 |
| IPR020602 | GTP cyclohydrolase I domain | 41 | 39 | 2 | 1.95E-07 |
| IPR001098 | DNA-directed DNA polymerase | 58 | 52 | 6 | 3.12E-07 |
| IPR002870 | Peptidase M12B | 24 | 2 | 22 | 3.59E-07 |
| IPR024732 | Alpha-N-acetylglucosaminidase | 28 | 28 | 0 | 5.10E-07 |
| IPR013847 | POU domain | 28 | 28 | 0 | 5.10E-07 |
| IPR000472 | Activin types I and II receptor domain | 49 | 45 | 4 | 5.29E-07 |
| IPR001320 | Ionotropic glutamate receptor | 158 | 123 | 35 | 5.75E-07 |
| IPR013680 | Voltage-dependent calcium channel | 16 | 0 | 16 | 7.17E-07 |
| IPR010579 | MHC class I | 16 | 0 | 16 | 7.17E-07 |
| IPR032455 | Cadherin | 16 | 0 | 16 | 7.17E-07 |
| IPR002350 | Kazal domain | 50 | 12 | 38 | 8.23E-07 |
| IPR003010 | Carbon-nitrogen hydrolase | 103 | 84 | 19 | 1.00E-06 |
| IPR016179 | Insulin-like | 96 | 79 | 17 | 1.13E-06 |
| IPR029052 | Metallo-dependent phosphatase-like | 161 | 124 | 37 | 1.19E-06 |
| IPR006158 | Cobalamin (vitamin B12)-binding domain | 41 | 38 | 3 | 1.68E-06 |
| IPR029058 | Alpha/Beta hydrolase fold | 554 | 380 | 174 | 1.73E-06 |
| IPR029047 | Heat shock protein 70kD | 15 | 0 | 15 | 1.74E-06 |
| IPR009254 | Laminin alpha | 15 | 0 | 15 | 1.74E-06 |
| IPR000477 | Reverse transcriptase domain | 26 | 26 | 0 | 1.80E-06 |
| IPR006782 | Platelet-derived growth factor | 46 | 11 | 35 | 2.76E-06 |
| IPR009011 | Mannose-6-phosphate receptor binding domain superfamily | 33 | 6 | 27 | 2.82E-06 |
| IPR003331 | UDP-N-acetylglucosamine 2-epimerase domain | 30 | 29 | 1 | 2.89E-06 |
| IPR006612 | THAP-type zinc finger | 31 | 30 | 1 | 2.92E-06 |
| IPR001322 | Lamin tail domain | 56 | 49 | 7 | 3.97E-06 |
| IPR015798 | Copper amine oxidase | 14 | 0 | 14 | 4.20E-06 |
| IPR006134 | DNA-directed DNA polymerase | 14 | 0 | 14 | 4.20E-06 |
| IPR008916 | Retrovirus capsid | 14 | 0 | 14 | 4.20E-06 |
| IPR008519 | Tandem-repeating region of mucin | 14 | 0 | 14 | 4.20E-06 |
| IPR022418 | Porphobilinogen deaminase | 29 | 28 | 1 | 4.94E-06 |
| IPR000326 | Phosphatidic acid phosphatase type 2/haloperoxidase | 48 | 43 | 5 | 4.94E-06 |
| IPR027309 | P2X purinoreceptor extracellular domain superfamily | 32 | 6 | 26 | 5.43E-06 |
| IPR011992 | EF-hand domain pair | 248 | 110 | 138 | 5.48E-06 |
| IPR014010 | REJ domain | 40 | 9 | 31 | 5.60E-06 |
| IPR000834 | Peptidase M14 | 37 | 8 | 29 | 5.81E-06 |
| IPR008984 | SMAD/FHA domain superfamily | 84 | 69 | 15 | 6.25E-06 |

| | | | | | |
|---|---|---|---|---|---|
| IPR031162 | CBP/p300-type histone acetyltransferase domain | 43 | 39 | 4 | 6.37E-06 |
| IPR002049 | Laminin EGF domain | 58 | 17 | 41 | 6.87E-06 |
| IPR027936 | Ephrin receptor | 17 | 1 | 16 | 7.45E-06 |
| IPR009048 | Alpha-macroglobulin | 17 | 1 | 16 | 7.45E-06 |
| IPR000294 | Gamma-carboxyglutamic acid-rich (GLA) domain | 211 | 155 | 56 | 9.00E-06 |
| IPR000873 | AMP-dependent synthetase/ligase | 53 | 15 | 38 | 9.10E-06 |
| IPR011764 | Biotin carboxylation domain | 22 | 22 | 0 | 9.86E-06 |
| IPR016039 | Thiolase-like | 49 | 43 | 6 | 1.45E-05 |
| IPR015794 | Pyruvate kinase | 27 | 26 | 1 | 1.47E-05 |
| IPR003112 | Olfactomedin-like domain | 71 | 59 | 12 | 1.64E-05 |
| IPR020843 | Polyketide synthase | 16 | 1 | 15 | 1.70E-05 |
| IPR003993 | Treacle protein domain | 16 | 1 | 15 | 1.70E-05 |
| IPR001747 | Lipid transport protein | 33 | 7 | 26 | 1.94E-05 |
| IPR009080 | Aminoacyl-tRNA synthetase | 30 | 6 | 24 | 2.01E-05 |
| IPR000257 | Uroporphyrinogen decarboxylase (URO-D) | 48 | 42 | 6 | 2.33E-05 |
| IPR021072 | Melanoma associated antigen | 12 | 0 | 12 | 2.46E-05 |
| IPR008274 | Aldehyde oxidase/xanthine dehydrogenase | 12 | 0 | 12 | 2.46E-05 |
| IPR001180 | Citron homology (CNH) domain | 12 | 0 | 12 | 2.46E-05 |
| IPR012674 | Calycin | 67 | 22 | 45 | 2.84E-05 |
| IPR000312 | Glycosyl transferase | 39 | 35 | 4 | 2.89E-05 |
| IPR001024 | PLAT/LH2 domain | 90 | 33 | 57 | 3.56E-05 |
| IPR017981 | GPCR | 124 | 50 | 74 | 4.92E-05 |
| IPR003594 | Histidine kinase/HSP90-like ATPase | 126 | 96 | 30 | 5.66E-05 |
| IPR016040 | NAD(P)-binding domain | 534 | 359 | 175 | 5.69E-05 |
| IPR012932 | Vitamin K epoxide reductase | 19 | 19 | 0 | 5.89E-05 |
| IPR024810 | Mab-21 domain | 11 | 0 | 11 | 5.97E-05 |
| IPR018484 | Carbohydrate kinase | 11 | 0 | 11 | 5.97E-05 |
| IPR010526 | Sodium ion transport-associated | 84 | 31 | 53 | 8.16E-05 |
| IPR003599 | Immunoglobulin subtype | 17 | 2 | 15 | 8.88E-05 |
| IPR002219 | Protein kinase C-like | 95 | 74 | 21 | 9.83E-05 |
| IPR002190 | MAGE homology domain | 20 | 3 | 17 | 0.000100355 |
| IPR023210 | NADP-dependent oxidoreductase domain | 79 | 29 | 50 | 0.000127732 |
| IPR001296 | Glycosyl transferase | 28 | 26 | 2 | 0.000135215 |
| IPR001107 | Band 7 domain | 40 | 35 | 5 | 0.000140989 |
| IPR009051 | Alpha-helical ferredoxin | 23 | 22 | 1 | 0.000142935 |
| IPR028074 | PTHB1 | 10 | 0 | 10 | 0.00014449 |
| IPR024571 | ERAP1-like C-terminal domain | 10 | 0 | 10 | 0.00014449 |
| IPR029020 | Ammonium/urea transporter | 10 | 0 | 10 | 0.00014449 |
| IPR011657 | Concentrative nucleoside transporter C-terminal domain | 10 | 0 | 10 | 0.00014449 |
| IPR014044 | CAP domain | 10 | 0 | 10 | 0.00014449 |
| IPR000569 | HECT domain | 51 | 43 | 8 | 0.000153496 |
| IPR001879 | GPCR | 34 | 9 | 25 | 0.000174019 |
| IPR032200 | Transcription factor COE | 17 | 17 | 0 | 0.000205007 |
| IPR014729 | Rossmann-like alpha/beta/alpha sandwich fold | 200 | 143 | 57 | 0.000226438 |
| IPR012677 | Nucleotide-binding alpha-beta plait domain superfamily | 54 | 18 | 36 | 0.000244879 |
| IPR033118 | EXPERA domain | 21 | 20 | 1 | 0.000254358 |
| IPR001104 | 3-oxo-5-alpha-steroid 4-dehydrogenase | 30 | 27 | 3 | 0.00027754 |

| | | | | | |
|---|---|---|---|---|---|
| IPR013158 | APOBEC-like | 31 | 8 | 23 | 0.000332108 |
| IPR011547 | SLC26A/SulP transporter domain | 106 | 80 | 26 | 0.000349665 |
| IPR028565 | Mu homology domain | 9 | 0 | 9 | 0.000349863 |
| IPR018292 | A-kinase anchor 110kDa | 9 | 0 | 9 | 0.000349863 |
| IPR024779 | 2OGFeDO | 9 | 0 | 9 | 0.000349863 |
| IPR028235 | Dynein assembly factor 3 | 9 | 0 | 9 | 0.000349863 |
| IPR003137 | PA domain | 9 | 0 | 9 | 0.000349863 |
| IPR002791 | Damage-control phosphatase ARMT1-like | 9 | 0 | 9 | 0.000349863 |
| IPR002502 | N-acetylmuramoyl-L-alanine amidase domain | 9 | 0 | 9 | 0.000349863 |
| IPR001932 | PPM-type phosphatase domain | 9 | 0 | 9 | 0.000349863 |
| IPR010630 | Olduvai domain | 9 | 0 | 9 | 0.000349863 |
| IPR010991 | p53 | 41 | 35 | 6 | 0.000360155 |
| IPR031160 | F-BAR domain | 26 | 6 | 20 | 0.000416028 |
| IPR029006 | ADF-H/Gelsolin-like domain superfamily | 63 | 23 | 40 | 0.000464065 |
| IPR023346 | Lysozyme-like domain superfamily | 181 | 129 | 52 | 0.000488914 |
| IPR011598 | Myc-type | 83 | 64 | 19 | 0.000499682 |
| IPR012340 | Nucleic acid-binding | 97 | 40 | 57 | 0.000602652 |
| IPR025766 | ADD domain | 14 | 14 | 0 | 0.000662938 |
| IPR032466 | Metal-dependent hydrolase | 107 | 80 | 27 | 0.000755224 |
| IPR011989 | Armadillo-like helical | 20 | 4 | 16 | 0.000843474 |
| IPR012315 | KASH domain | 8 | 0 | 8 | 0.000847125 |
| IPR012308 | DNA ligase | 8 | 0 | 8 | 0.000847125 |
| IPR032471 | GAIN domain | 8 | 0 | 8 | 0.000847125 |
| IPR013035 | Phosphoenolpyruvate carboxykinase | 8 | 0 | 8 | 0.000847125 |
| IPR027007 | C2 DOCK-type domain | 8 | 0 | 8 | 0.000847125 |
| IPR027841 | Interleukin-17 receptor C/E | 8 | 0 | 8 | 0.000847125 |
| IPR029155 | SIPAR domain | 8 | 0 | 8 | 0.000847125 |
| IPR032431 | Immunoglobulin C1-set, C-terminal extension | 8 | 0 | 8 | 0.000847125 |
| IPR031320 | GAGE | 8 | 0 | 8 | 0.000847125 |
| IPR012319 | Formamidopyrimidine-DNA glycosylase | 8 | 0 | 8 | 0.000847125 |
| IPR002999 | Tudor domain | 8 | 0 | 8 | 0.000847125 |
| IPR000375 | Dynamin stalk domain | 35 | 30 | 5 | 0.000868227 |
| IPR000716 | Thyroglobulin type-1 | 43 | 14 | 29 | 0.000892652 |
| IPR011029 | Death-like domain superfamily | 168 | 77 | 91 | 0.000929213 |
| IPR000699 | RIH domain | 56 | 45 | 11 | 0.000940169 |
| IPR016093 | MIR motif | 56 | 45 | 11 | 0.000940169 |
| IPR013057 | Amino acid transporter | 29 | 8 | 21 | 0.000978988 |
| IPR001750 | NADH:quinone oxidoreductase/Mrp antiporter | 36 | 11 | 25 | 0.000982521 |
| IPR000885 | Fibrillar collagen | 66 | 25 | 41 | 0.000984656 |
| IPR021887 | Domain of unknown function DUF3498 | 11 | 1 | 10 | 0.000992674 |
| IPR008253 | Marvel domain | 11 | 1 | 10 | 0.000992674 |
| IPR016187 | C-type lectin fold | 291 | 143 | 148 | 0.001013899 |
| IPR016177 | DNA-binding domain superfamily | 52 | 42 | 10 | 0.001024879 |
| IPR013121 | Ferric reductase | 23 | 21 | 2 | 0.00102493 |
| IPR033644 | Ferrochelatase | 23 | 21 | 2 | 0.00102493 |
| IPR006545 | EYA domain | 13 | 13 | 0 | 0.001178223 |
| IPR008121 | Transcription factor AP-2 alpha | 13 | 13 | 0 | 0.001178223 |
| IPR029067 | CDC48 domain 2-like superfamily | 13 | 13 | 0 | 0.001178223 |
| IPR011527 | ABC transporter type 1 | 812 | 431 | 381 | 0.001234769 |

| | | | | | |
|---|---|---|---|---|---|
| IPR004865 | HSR domain | 18 | 17 | 1 | 0.001260097 |
| IPR032630 | P-type ATPase | 14 | 2 | 12 | 0.001435721 |
| IPR008952 | Tetraspanin | 61 | 48 | 13 | 0.001566978 |
| IPR001214 | SET domain | 88 | 66 | 22 | 0.001607143 |
| IPR012336 | Thioredoxin-like fold | 201 | 96 | 105 | 0.001970698 |
| IPR002937 | Amine oxidase | 25 | 22 | 3 | 0.002043348 |
| IPR017854 | Metallothionein domain superfamily | 7 | 0 | 7 | 0.002051105 |
| IPR031474 | Protein phosphatase 1 regulatory subunit 26 | 7 | 0 | 7 | 0.002051105 |
| IPR011008 | Dimeric alpha-beta barrel | 7 | 0 | 7 | 0.002051105 |
| IPR025136 | MAP3K | 7 | 0 | 7 | 0.002051105 |
| IPR000922 | D-galactoside/L-rhamnose binding SUEL lectin domain | 7 | 0 | 7 | 0.002051105 |
| IPR031907 | Germinal-centre associated nuclear protein | 7 | 0 | 7 | 0.002051105 |
| IPR010926 | Class I myosin tail homology domain | 7 | 0 | 7 | 0.002051105 |
| IPR027357 | DOCKER domain | 7 | 0 | 7 | 0.002051105 |
| IPR024240 | Alpha-N-acetylglucosaminidase | 12 | 12 | 0 | 0.002114899 |
| IPR006594 | LIS1 homology motif | 12 | 12 | 0 | 0.002114899 |
| IPR013803 | Amyloidogenic glycoprotein | 12 | 12 | 0 | 0.002114899 |
| IPR003032 | Ryanodine receptor Ryr | 10 | 1 | 9 | 0.002198217 |
| IPR001763 | Rhodanese-like domain | 30 | 9 | 21 | 0.002356617 |
| IPR012351 | Four-helical cytokine | 676 | 358 | 318 | 0.002488008 |
| IPR003191 | Guanylate-binding protein/Atlastin | 34 | 11 | 23 | 0.002561672 |
| IPR023578 | Ras guanine nucleotide exchange factor domain superfamily | 61 | 24 | 37 | 0.002593501 |
| IPR001487 | Bromodomain | 38 | 13 | 25 | 0.002712016 |
| IPR000731 | Sterol-sensing domain | 42 | 34 | 8 | 0.002741691 |
| IPR027397 | Catenin binding domain superfamily | 13 | 2 | 11 | 0.002782118 |
| IPR014001 | Helicase superfamily 1/2 | 13 | 2 | 11 | 0.002782118 |
| IPR000082 | SEA domain | 18 | 4 | 14 | 0.002787353 |
| IPR025837 | CFTR regulator domain | 21 | 19 | 2 | 0.002916361 |
| IPR031701 | Homeobox protein SIX1 | 21 | 19 | 2 | 0.002916361 |
| IPR000772 | Ricin B | 15 | 3 | 12 | 0.003012679 |
| IPR005480 | Carbamoyl-phosphate synthetase | 16 | 15 | 1 | 0.003772801 |
| IPR002100 | Transcription factor | 11 | 11 | 0 | 0.003838191 |
| IPR010536 | Repulsive guidance molecule | 11 | 11 | 0 | 0.003838191 |
| IPR034154 | Archaeal primase DnaG/twinkle | 11 | 11 | 0 | 0.003838191 |
| IPR008928 | Six-hairpin glycosidase superfamily | 72 | 30 | 42 | 0.00389772 |
| IPR010994 | RuvA domain 2-like | 9 | 1 | 8 | 0.004825225 |
| IPR027295 | Quinonprotein alcohol dehydrogenase-like-domain | 9 | 1 | 8 | 0.004825225 |
| IPR032803 | PLD-like domain | 9 | 1 | 8 | 0.004825225 |
| IPR029045 | ClpP/crotonase-like domain superfamily | 189 | 130 | 59 | 0.004841277 |
| IPR001357 | BRCT domain | 111 | 50 | 61 | 0.00491063 |
| IPR021040 | LRRC8 | 6 | 0 | 6 | 0.004966133 |
| IPR003726 | Homocysteine-binding domain | 6 | 0 | 6 | 0.004966133 |
| IPR009017 | Green fluorescent protein | 6 | 0 | 6 | 0.004966133 |
| IPR010979 | Ribosomal protein S13-like | 6 | 0 | 6 | 0.004966133 |
| IPR004102 | Poly(ADP-ribose) polymerase | 6 | 0 | 6 | 0.004966133 |
| IPR007725 | Timeless | 6 | 0 | 6 | 0.004966133 |
| IPR001194 | cDENN domain | 6 | 0 | 6 | 0.004966133 |

| | | | | | |
|---|---|---|---|---|---|
| IPR024309 | Nuclear Testis protein | 6 | 0 | 6 | 0.004966133 |
| IPR001599 | Alpha-2-macroglobulin | 6 | 0 | 6 | 0.004966133 |
| IPR032680 | SUN domain-containing protein 1 | 6 | 0 | 6 | 0.004966133 |
| IPR022409 | PKD/Chitinase domain | 6 | 0 | 6 | 0.004966133 |
| IPR016182 | Copper amine oxidase | 6 | 0 | 6 | 0.004966133 |
| IPR000674 | Aldehyde oxidase/xanthine dehydrogenase | 6 | 0 | 6 | 0.004966133 |
| IPR010600 | Inter-alpha-trypsin inhibitor heavy chain | 6 | 0 | 6 | 0.004966133 |
| IPR013992 | Adenylate cyclase-associated CAP | 6 | 0 | 6 | 0.004966133 |
| IPR013697 | DNA polymerase epsilon | 6 | 0 | 6 | 0.004966133 |
| IPR000959 | POLO box domain | 6 | 0 | 6 | 0.004966133 |
| IPR008942 | ENTH/VHS | 6 | 0 | 6 | 0.004966133 |
| IPR018491 | SLC12A transporter | 53 | 41 | 12 | 0.0051256 |
| IPR025994 | BRCA1 | 12 | 2 | 10 | 0.005400998 |
| IPR017978 | GPCR family 3 | 78 | 58 | 20 | 0.005423893 |
| IPR020630 | Tetrahydrofolate dehydrogenase/cyclohydrolase | 15 | 14 | 1 | 0.006576816 |
| IPR001048 | Aspartate/glutamate/uridylate kinase | 15 | 14 | 1 | 0.006576816 |
| IPR000157 | Toll/interleukin-1 receptor homology (TIR) domain | 63 | 26 | 37 | 0.0067286 |
| IPR000219 | Dbl homology (DH) domain | 63 | 26 | 37 | 0.0067286 |
| IPR029030 | Caspase-like domain superfamily | 47 | 18 | 29 | 0.00695587 |
| IPR013234 | PIGA | 10 | 10 | 0 | 0.007046911 |
| IPR024986 | Sister chromatid cohesion C-terminal domain | 10 | 10 | 0 | 0.007046911 |
| IPR009061 | Putative DNA-binding domain superfamily | 10 | 10 | 0 | 0.007046911 |
| IPR032419 | NF-kappa-B essential modulator NEMO | 10 | 10 | 0 | 0.007046911 |
| IPR000008 | C2 domain | 304 | 155 | 149 | 0.007155847 |
| IPR011009 | Protein kinase-like domain superfamily | 2593 | 1588 | 1005 | 0.007200292 |
| IPR013819 | Lipoxygenase | 91 | 66 | 25 | 0.007499111 |
| IPR032444 | Keratin type II head | 42 | 16 | 26 | 0.00759367 |
| IPR005135 | Endonuclease/exonuclease/phosphatase | 159 | 110 | 49 | 0.007622323 |
| IPR015988 | STAT transcription factor | 29 | 24 | 5 | 0.007821712 |
| IPR011500 | GPCR | 18 | 16 | 2 | 0.007960129 |
| IPR001111 | TGF-beta | 72 | 31 | 41 | 0.008197701 |
| IPR000315 | B-box-type zinc finger | 16 | 4 | 12 | 0.009098711 |
| IPR000203 | GPS motif | 23 | 7 | 16 | 0.009403224 |
| IPR002931 | Transglutaminase-like | 121 | 85 | 36 | 0.009584465 |
| IPR009078 | Ferritin-like superfamily | 35 | 28 | 7 | 0.00977277 |

**Table S3.** Median spatial distance values for 30 types of position-based sequence annotations and 2 structural features, which are used to impute the missing values in SAV feature vectors. The differences between PDB and AlphaFold models mainly due to higher coverage of AlphaFold models which allowed the calculation of spatial distances between variations and annotated positions that are very far away from each other.

| Annotation type | Imputed value (Angstroms) | |
|---|---|---|
| | PDB models | AlphaFold models |
| disulfide_dist | 17.84 | 20.71 |
| intMet_dist | 30.80 | 46.67 |
| intramembrane_dist | 24.96 | 28.13 |
| naturalVariant_dist | 13.12 | 15.50 |
| dnaBinding_dist | 23.62 | 35.94 |
| activeSite_dist | 18.97 | 21.84 |
| nucleotideBinding_dist | 20.87 | 25.15 |
| lipidation_dist | 29.59 | 45.15 |
| site_dist | 20.70 | 29.81 |
| transmembrane_dist | 12.70 | 29.91 |
| crosslink_dist | 22.85 | 34.67 |
| mutagenesis_dist | 17.21 | 24.72 |
| strand_dist | 9.80 | 10.66 |
| helix_dist | 9.00 | 11.55 |
| turn_dist | 15.99 | 13.02 |
| metalBinding_dist | 16.82 | 21.54 |
| repeat_dist | 20.46 | 27.42 |
| caBinding_dist | 24.58 | 38.39 |
| topologicalDomain_dist | 9.99 | 30.44 |
| bindingSite_dist | 17.43 | 20.90 |
| region_dist | 20.08 | 25.82 |
| signalPeptide_dist | 30.91 | 46.12 |
| modifiedResidue_dist | 20.86 | 32.10 |
| zincFinger_dist | 22.14 | 35.96 |
| motif_dist | 21.91 | 35.86 |
| coiledCoil_dist | 28.45 | 37.88 |
| peptide_dist | 17.81 | 19.09 |
| transitPeptide_dist | 25.12 | 35.20 |
| glycosylation_dist | 20.33 | 26.95 |
| propeptide_dist | 22.36 | 37.48 |
| sasa (structural feature) | 29.50 | 35.60 |
| domains_3Ddist (structural feature) | 24.50 | 29.78 |

**Table S4.** Hyper-parameter optimization test results of ASCARIS-PDB (via random forest classification). Maximum performance values for each metric are shown in bold.

| Ntrees* | MaxDecSpt* | PredSelRan* | AUROC | Accuracy | Recall | Precision | F1-score | MCC | TP | FN | FP | TN |
|---|---|---|---|---|---|---|---|---|---|---|---|---|
| 50 | 3 | 2 | 0.64 | 0.68 | 0.65 | 0.66 | 0.62 | 0.23 | 35470 | 17042 | 19368 | 24394 |
| 50 | 3 | 8 | 0.64 | 0.68 | 0.65 | 0.66 | 0.62 | 0.23 | 35505 | 17007 | 19393 | 24369 |
| 50 | 3 | 24 | 0.68 | 0.66 | 0.70 | 0.68 | 0.66 | 0.32 | 34708 | 17804 | 14786 | 28976 |
| 50 | 3 | 68 | 0.73 | 0.69 | 0.70 | 0.70 | 0.67 | 0.34 | 36267 | 16245 | 15391 | 28371 |
| 50 | 81 | 2 | 0.76 | 0.74 | 0.71 | 0.73 | 0.70 | 0.39 | 39090 | 13422 | 15818 | 27944 |
| 50 | 81 | 8 | 0.79 | 0.77 | 0.73 | 0.75 | 0.72 | 0.43 | 40662 | 11850 | 15054 | 28708 |
| 50 | 81 | 24 | 0.82 | 0.82 | 0.74 | 0.78 | 0.74 | 0.48 | 43196 | 9316 | 15461 | 28301 |
| 50 | 81 | 68 | 0.82 | 0.81 | 0.75 | 0.77 | 0.74 | 0.48 | 42279 | 10233 | 14393 | 29369 |
| 50 | 2187 | 2 | 0.84 | 0.81 | 0.77 | 0.79 | 0.77 | 0.53 | 42426 | 10086 | 12510 | 31252 |
| 50 | 2187 | 8 | 0.86 | 0.83 | 0.78 | 0.80 | 0.78 | 0.56 | 43434 | 9078 | 12079 | 31683 |
| 50 | 2187 | 24 | 0.87 | 0.84 | 0.78 | 0.81 | 0.79 | 0.57 | 44277 | 8235 | 12395 | 31367 |
| 50 | 2187 | 68 | 0.87 | 0.84 | 0.78 | 0.81 | 0.79 | 0.57 | 44163 | 8349 | 12233 | 31529 |
| 50 | 96273 | 2 | 0.86 | 0.83 | 0.79 | 0.81 | 0.79 | 0.57 | 43755 | 8757 | 11906 | 31856 |
| 50 | 96273 | 8 | 0.88 | 0.85 | 0.80 | **0.83** | **0.81** | 0.61 | 44623 | 7889 | 10838 | 32924 |
| 50 | 96273 | 24 | 0.88 | 0.84 | **0.81** | 0.82 | 0.80 | 0.60 | 44322 | 8190 | 10620 | 33142 |
| 50 | 96273 | 68 | 0.88 | 0.84 | 0.80 | 0.82 | 0.80 | 0.60 | 44115 | 8397 | 10878 | 32884 |
| 150 | 3 | 2 | 0.64 | 0.68 | 0.65 | 0.66 | 0.62 | 0.23 | 35506 | 17006 | 19397 | 24365 |
| 150 | 3 | 8 | 0.64 | 0.68 | 0.65 | 0.66 | 0.62 | 0.23 | 35503 | 17009 | 19396 | 24366 |
| 150 | 3 | 24 | 0.68 | 0.67 | 0.70 | 0.68 | 0.66 | 0.32 | 34936 | 17576 | 15007 | 28755 |
| 150 | 3 | 68 | 0.73 | 0.68 | 0.71 | 0.69 | 0.67 | 0.34 | 35453 | 17059 | 14813 | 28949 |
| 150 | 81 | 2 | 0.76 | 0.75 | 0.71 | 0.73 | 0.70 | 0.38 | 39205 | 13307 | 15945 | 27817 |
| 150 | 81 | 8 | 0.79 | 0.77 | 0.73 | 0.75 | 0.72 | 0.43 | 40406 | 12106 | 14802 | 28960 |
| 150 | 81 | 24 | 0.82 | 0.82 | 0.74 | 0.78 | 0.74 | 0.48 | 43234 | 9278 | 15544 | 28218 |
| 150 | 81 | 68 | 0.82 | 0.80 | 0.75 | 0.77 | 0.74 | 0.48 | 42248 | 10264 | 14314 | 29448 |
| 150 | 2187 | 2 | 0.84 | 0.81 | 0.77 | 0.79 | 0.77 | 0.53 | 42564 | 9948 | 12384 | 31378 |
| 150 | 2187 | 8 | 0.86 | 0.83 | 0.78 | 0.81 | 0.78 | 0.56 | 43530 | 8982 | 12062 | 31700 |
| 150 | 2187 | 24 | 0.87 | 0.85 | 0.78 | 0.81 | 0.79 | 0.57 | 44410 | 8102 | 12414 | 31348 |
| 150 | 2187 | 68 | 0.87 | 0.84 | 0.78 | 0.81 | 0.79 | 0.57 | 44309 | 8203 | 12169 | 31593 |
| 150 | 96273 | 2 | 0.86 | 0.84 | 0.79 | 0.81 | 0.79 | 0.57 | 43983 | 8529 | 11906 | 31856 |
| 150 | 96273 | 8 | **0.89** | 0.85 | **0.81** | **0.83** | **0.81** | 0.61 | 44795 | 7717 | 10735 | 33027 |
| 150 | 96273 | 24 | **0.89** | 0.85 | **0.81** | **0.83** | **0.81** | 0.61 | 44669 | 7843 | 10795 | 32967 |

| | | | | | | | | | | | | |
|---|---|---|---|---|---|---|---|---|---|---|---|---|
| 150 | 96273 | 68 | 0.88 | 0.85 | 0.80 | 0.82 | 0.80 | 0.60 | 44445 | 8067 | 10936 | 32826 |
| 300 | 3 | 2 | 0.64 | 0.68 | 0.65 | 0.66 | 0.62 | 0.23 | 35452 | 17060 | 19373 | 24389 |
| 300 | 3 | 8 | 0.64 | 0.68 | 0.65 | 0.66 | 0.62 | 0.23 | 35481 | 17031 | 19380 | 24382 |
| 300 | 3 | 24 | 0.68 | 0.67 | 0.70 | 0.68 | 0.66 | 0.32 | 35204 | 17308 | 15269 | 28493 |
| 300 | 3 | 68 | 0.73 | 0.72 | 0.70 | 0.71 | 0.68 | 0.35 | 37798 | 14714 | 16014 | 27748 |
| 300 | 81 | 2 | 0.76 | 0.75 | 0.71 | 0.73 | 0.70 | 0.39 | 39322 | 13190 | 15956 | 27806 |
| 300 | 81 | 8 | 0.79 | 0.77 | 0.73 | 0.75 | 0.72 | 0.43 | 40571 | 11941 | 14952 | 28810 |
| 300 | 81 | 24 | 0.82 | 0.82 | 0.74 | 0.78 | 0.74 | 0.48 | 43283 | 9229 | 15514 | 28248 |
| 300 | 81 | 68 | 0.82 | 0.81 | 0.75 | 0.77 | 0.74 | 0.48 | 42291 | 10221 | 14345 | 29417 |
| 300 | 2187 | 2 | 0.84 | 0.81 | 0.77 | 0.79 | 0.77 | 0.53 | 42587 | 9925 | 12388 | 31374 |
| 300 | 2187 | 8 | 0.86 | 0.83 | 0.78 | 0.80 | 0.78 | 0.56 | 43477 | 9035 | 12052 | 31710 |
| 300 | 2187 | 24 | 0.87 | 0.85 | 0.78 | 0.81 | 0.79 | 0.57 | 44413 | 8099 | 12415 | 31347 |
| 300 | 2187 | 68 | 0.87 | 0.84 | 0.78 | 0.81 | 0.79 | 0.57 | 44258 | 8254 | 12287 | 31475 |
| 300 | 96273 | 2 | 0.87 | 0.84 | 0.79 | 0.81 | 0.79 | 0.57 | 44138 | 8374 | 11913 | 31849 |
| 300 | 96273 | 8 | **0.89** | **0.86** | **0.81** | **0.83** | **0.81** | **0.62** | 44914 | 7598 | 10593 | 33169 |
| 300 | 96273 | 24 | **0.89** | 0.85 | **0.81** | **0.83** | **0.81** | 0.61 | 44749 | 7763 | 10795 | 32967 |
| 300 | 96273 | 68 | 0.88 | 0.85 | 0.80 | 0.82 | 0.80 | 0.60 | 44458 | 8054 | 10899 | 32863 |
| 500 | 3 | 2 | 0.64 | 0.68 | 0.65 | 0.66 | 0.62 | 0.23 | 35497 | 17015 | 19396 | 24366 |
| 500 | 3 | 8 | 0.64 | 0.68 | 0.65 | 0.66 | 0.62 | 0.23 | 35517 | 16995 | 19406 | 24356 |
| 500 | 3 | 24 | 0.68 | 0.67 | 0.70 | 0.68 | 0.66 | 0.32 | 34974 | 17538 | 15017 | 28745 |
| 500 | 3 | 68 | 0.73 | 0.68 | 0.71 | 0.70 | 0.67 | 0.35 | 35881 | 16631 | 14676 | 29086 |
| 500 | 81 | 2 | 0.76 | 0.75 | 0.71 | 0.73 | 0.70 | 0.39 | 39207 | 13305 | 15883 | 27879 |
| 500 | 81 | 8 | 0.79 | 0.77 | 0.73 | 0.75 | 0.72 | 0.43 | 40537 | 11975 | 14911 | 28851 |
| 500 | 81 | 24 | 0.82 | 0.82 | 0.74 | 0.78 | 0.74 | 0.48 | 43224 | 9288 | 15507 | 28255 |
| 500 | 81 | 68 | 0.82 | 0.80 | 0.75 | 0.77 | 0.74 | 0.48 | 42248 | 10264 | 14365 | 29397 |
| 500 | 2187 | 2 | 0.84 | 0.81 | 0.77 | 0.79 | 0.77 | 0.53 | 42655 | 9857 | 12403 | 31359 |
| 500 | 2187 | 8 | 0.86 | 0.83 | 0.78 | 0.81 | 0.78 | 0.56 | 43527 | 8985 | 12064 | 31698 |
| 500 | 2187 | 24 | 0.87 | 0.85 | 0.78 | 0.81 | 0.79 | 0.57 | 44490 | 8022 | 12407 | 31355 |
| 500 | 2187 | 68 | 0.87 | 0.84 | 0.78 | 0.81 | 0.79 | 0.57 | 44284 | 8228 | 12222 | 31540 |
| 500 | 96273 | 2 | 0.87 | 0.84 | 0.79 | 0.81 | 0.79 | 0.57 | 44100 | 8412 | 11880 | 31882 |
| 500 | 96273 | 8 | **0.89** | 0.85 | **0.81** | **0.83** | **0.81** | 0.61 | 44865 | 7647 | 10715 | 33047 |
| 500 | 96273 | 24 | **0.89** | 0.85 | **0.81** | **0.83** | **0.81** | 0.61 | 44791 | 7721 | 10732 | 33030 |
| 500 | 96273 | 68 | 0.88 | 0.85 | 0.80 | 0.82 | 0.80 | 0.60 | 44522 | 7990 | 11033 | 32729 |

* Ntrees: number of trees, MaxDecSpt: maximum number of decision splits, PredSelRan: number of predictors to select at random for each split.

**Table S5.** Variation data points selected from the MutationTaster benchmark test dataset which are found be challenging in terms of predicting their effects, considering the prediction performance of multiple variant effect predictors from the literature and ASCARIS.

| UniProt accession | wt_residue | position | mut_residue | label |
|---|---|---|---|---|
| A5D8T8 | T | 151 | M | 0 |
| A5D8W1 | T | 885 | M | 0 |
| A6NCV1 | G | 86 | D | 0 |
| A6NHA9 | C | 252 | Y | 0 |
| A6NHA9 | S | 240 | F | 0 |
| O00170 | R | 304 | Q | 1 |
| O00337 | Q | 237 | K | 0 |
| O60403 | L | 40 | Q | 0 |
| O60437 | Q | 1573 | E | 0 |
| O60566 | Q | 921 | H | 1 |
| O75712 | I | 141 | V | 1 |
| O75712 | N | 166 | S | 1 |
| O75752 | E | 266 | A | 0 |
| O75752 | G | 271 | R | 0 |
| O95050 | F | 254 | C | 0 |
| O95452 | T | 5 | M | 1 |
| O95477 | D | 1289 | N | 1 |
| P00966 | R | 86 | C | 1 |
| P01185 | V | 67 | A | 1 |
| P02647 | A | 199 | P | 1 |
| P02656 | K | 78 | E | 1 |
| P02671 | E | 545 | V | 1 |
| P02708 | V | 201 | M | 1 |
| P03950 | K | 41 | E | 1 |
| P03950 | R | 55 | K | 1 |
| P03950 | S | 52 | N | 1 |
| P04062 | D | 448 | H | 1 |
| P04424 | V | 178 | M | 1 |
| P07902 | D | 113 | N | 1 |
| P07902 | E | 308 | K | 1 |
| P07902 | I | 198 | M | 1 |
| P07902 | K | 229 | N | 1 |
| P07902 | L | 227 | P | 1 |
| P07902 | L | 282 | V | 1 |
| P07902 | P | 265 | A | 1 |
| P07902 | Q | 38 | P | 1 |
| P07902 | Q | 9 | H | 1 |
| P07902 | R | 123 | Q | 1 |
| P07902 | R | 148 | Q | 1 |
| P07902 | R | 223 | S | 1 |
| P07902 | R | 262 | P | 1 |
| P07902 | T | 23 | A | 1 |
| P07902 | V | 125 | A | 1 |
| P07992 | F | 231 | L | 1 |
| P08246 | A | 61 | V | 1 |
| P08246 | S | 126 | L | 1 |
| P09874 | V | 762 | A | 0 |

| | | | | |
|---|---|---|---|---|
| P09936 | S | 18 | Y | 0 |
| P12259 | R | 334 | T | 1 |
| P12644 | E | 93 | G | 1 |
| P12644 | R | 287 | H | 1 |
| P15848 | C | 405 | Y | 1 |
| P19075 | G | 73 | A | 0 |
| P19526 | L | 242 | R | 0 |
| P21549 | E | 141 | D | 1 |
| P21549 | G | 42 | E | 1 |
| P21549 | I | 244 | T | 1 |
| P21549 | L | 150 | P | 1 |
| P22607 | A | 391 | E | 1 |
| P23560 | V | 66 | M | 0 |
| P24043 | A | 2587 | V | 0 |
| P24752 | A | 333 | P | 1 |
| P29033 | D | 159 | V | 1 |
| P29033 | S | 113 | R | 1 |
| P29317 | T | 940 | I | 1 |
| P30566 | S | 438 | P | 1 |
| P30793 | A | 196 | S | 1 |
| P30793 | K | 224 | R | 1 |
| P30793 | M | 221 | T | 1 |
| P30793 | R | 249 | S | 1 |
| P30954 | I | 103 | M | 0 |
| P31371 | S | 99 | N | 1 |
| P35670 | M | 645 | R | 1 |
| P35916 | H | 890 | Q | 0 |
| P38398 | E | 1038 | G | 0 |
| P41181 | A | 190 | T | 1 |
| P41181 | L | 22 | V | 1 |
| P41181 | T | 125 | M | 1 |
| P43026 | L | 373 | R | 1 |
| P43629 | P | 203 | S | 0 |
| P49748 | F | 458 | L | 1 |
| P55075 | K | 89 | E | 1 |
| P55291 | R | 92 | W | 1 |
| P68133 | D | 3 | Y | 1 |
| P78324 | R | 107 | S | 0 |
| P78363 | A | 1038 | V | 1 |
| P78363 | E | 1036 | K | 1 |
| Q01484 | T | 3744 | N | 1 |
| Q03692 | G | 18 | E | 1 |
| Q0ZLH3 | T | 54 | I | 1 |
| Q13009 | G | 247 | R | 0 |
| Q13336 | E | 44 | K | 0 |
| Q13618 | K | 459 | R | 1 |
| Q13705 | V | 494 | I | 1 |
| Q13797 | G | 507 | E | 0 |
| Q14624 | I | 85 | N | 0 |
| Q14674 | S | 614 | R | 0 |
| Q16600 | C | 209 | G | 0 |
| Q2KHM9 | E | 375 | D | 0 |

| | | | | |
|---|---|---|---|---|
| Q3YBM2 | A | 134 | T | 0 |
| Q5ST30 | R | 947 | Q | 0 |
| Q5T4I8 | A | 13 | D | 0 |
| Q5TD94 | R | 556 | H | 0 |
| Q5VTT5 | G | 662 | R | 0 |
| Q5VVB8 | F | 111 | V | 0 |
| Q6KF10 | A | 249 | E | 1 |
| Q6KF10 | K | 424 | R | 1 |
| Q6KF10 | P | 327 | H | 1 |
| Q6KF10 | Q | 253 | L | 1 |
| Q6T423 | R | 300 | T | 0 |
| Q6UXH1 | D | 182 | E | 0 |
| Q6UXH8 | R | 158 | C | 1 |
| Q6ZU80 | H | 732 | R | 0 |
| Q7Z3V5 | L | 573 | H | 0 |
| Q7Z3Z4 | Q | 327 | L | 0 |
| Q7Z6Z6 | L | 140 | F | 0 |
| Q86WV6 | R | 293 | Q | 0 |
| Q8IUA7 | K | 1306 | T | 0 |
| Q8IWI9 | P | 1523 | A | 0 |
| Q8IWZ6 | H | 323 | R | 1 |
| Q8N884 | P | 261 | H | 0 |
| Q8N960 | L | 602 | V | 0 |
| Q8NCM8 | G | 2461 | V | 1 |
| Q8NET1 | R | 36 | W | 0 |
| Q8NET6 | A | 271 | V | 0 |
| Q8NFJ9 | L | 518 | P | 1 |
| Q8NG31 | M | 598 | T | 0 |
| Q8NGC1 | I | 99 | N | 0 |
| Q8NGI0 | H | 264 | R | 0 |
| Q8NGS8 | S | 18 | F | 0 |
| Q8NH85 | A | 274 | V | 0 |
| Q8TAM1 | R | 34 | P | 1 |
| Q8TAM1 | V | 11 | G | 1 |
| Q8TB36 | Q | 218 | E | 1 |
| Q8TC05 | T | 103 | I | 0 |
| Q8WTS1 | E | 7 | K | 1 |
| Q92851 | L | 285 | F | 1 |
| Q96P65 | L | 344 | S | 0 |
| Q96QU1 | D | 435 | A | 0 |
| Q9BWD1 | K | 211 | R | 0 |
| Q9BX63 | M | 299 | I | 1 |
| Q9BXJ7 | T | 41 | I | 1 |
| Q9BYE9 | L | 1164 | M | 0 |
| Q9BYK8 | T | 2170 | M | 0 |
| Q9BZE9 | L | 252 | Q | 0 |
| Q9C0J9 | P | 384 | R | 0 |
| Q9GZV9 | R | 176 | Q | 1 |
| Q9H1B5 | T | 801 | R | 1 |
| Q9H324 | A | 25 | T | 1 |
| Q9H346 | I | 251 | T | 0 |
| Q9HAQ2 | R | 638 | W | 0 |

| | | | | |
|---|---|---|---|---|
| Q9HBY0 | T | 171 | K | 0 |
| Q9HCJ1 | M | 48 | T | 1 |
| Q9HCJ1 | P | 5 | T | 1 |
| Q9HCL2 | E | 131 | G | 0 |
| Q9NPC4 | G | 187 | D | 0 |
| Q9NRG9 | Q | 15 | K | 1 |
| Q9NVE7 | A | 547 | V | 0 |
| Q9NYK6 | D | 136 | E | 0 |
| Q9P2D6 | D | 1242 | G | 0 |
| Q9UBS4 | I | 264 | V | 0 |
| Q9UKP4 | T | 307 | M | 0 |
| Q9UKU7 | H | 289 | Q | 1 |
| Q9UMX9 | L | 374 | F | 0 |
| Q9Y267 | P | 560 | R | 0 |
| Q9Y281 | A | 35 | T | 1 |
| Q9Y3N9 | D | 296 | N | 0 |
